# BindFlow: a free, user-friendly pipeline for absolute binding free energy calculations using free energy perturbation or MM(PB/GB)SA

**DOI:** 10.1101/2025.09.25.678545

**Authors:** Alejandro Martínez León, Lucas Andersen, Jochen S. Hub

## Abstract

We present BindFlow, a Python-based software for automated absolute binding free energy (ABFE) calculations at the free energy perturbation (FEP) or at the molecular mechanics Poisson-Boltzmann/generalized Born surface area [MM(PB/GB)SA] level of theory. BindFlow is free, open-source, user-friendly, easily customizable, runs on workstations or distributed computing platforms, and provides extensive documentation and tutorials. BindFlow uses GROMACS as molecular dynamics engine and provides built-in support for the small-molecule force fields GAFF, OpenFF, and Espaloma. We test BindFlow by computing affinities for 139 ligand/target pairs, involving eight different targets including six soluble proteins, one membrane protein and one non-protein host– guest system. Quantified by Pearson, Kendall, and Spearman correlations coefficients, we find that the agreement of BindFlow predictions with experiments are overall similar to gold standards in the field. Interestingly, we find that MM(PB/GB)SA achieves correlations that, for some systems and force fields, approach those obtained with FEP, while requiring only a fraction of the computational cost. This study establishes BindFlow as a validated and accessible tool for ABFE calculations.

## Introduction

During the early stages of drug discovery, a large number of chemical compounds are evaluated with the aim of finding potent binders, a process that is both time-consuming and resource-intensive.^1,2^ Thus, computational models are widely used to estimate the binding free energy of small molecules to biological targets in order to prioritize compounds for followup synthesis and experimental evaluation. ^3–6^ In the 1950s, methods collectively known as alchemical free energy perturbation were introduced. ^7^ The term “free energy perturbation” (FEP) referred originally to a specific class of alchemical methods^8^ but has since been more generally applied to alchemical binding free energy methods, a notion followed in this study. FEP enabled the estimation of free energy differences at a fraction of the computational cost compared to conventional MD simulations. Nevertheless, FEP remained computationally demanding for high-throughput studies, rationalizing the development of computationally more efficient yet more approximate endpoint free energy techniques such as the molecular mechanics Poisson-Boltzmann/generalized Born surface area [MM(PB/GB)SA] methods,^9^ which achieved varying degrees of success.^10^ Recent advances in biomolecular simulation methods^6,11^ combined with the growth of computational power have enabled increasingly accurate affinity predictions,^6,12^ making both FEP and MM(PB/GB)SA calculations more routine.

Binding free energy estimation have been categorized into relative (RBFE) and absolute binding free energy (ABFE) calculations. ^11^ RBFE is typically preferred for ranking congeneric molecular series^13^ or for evaluating the effects of mutations on ligand binding.^14^ Alchemical RBFE methodologies have seen substantial advancements, resulting in numerous implementations.^15–29^ These developments have enabled alchemical RBFE methods to achieve remarkable accuracy, often with a root mean-squared error (RMSE) of 1–2 kcal mol^−1^ relative to experimental values, ^30^ making them highly effective for drug discovery applications.^13–15,31–37^

ABFE calculations yield the binding free energy relative to a standard state of the com-pound in solution. ^38^ ABFE calculations are particularly useful for studying highly diverse molecular sets,^4,39–41^ for binding pose validation,^42,43^ or for multi-target selectivity prediction.^44,45^ Alchemical ABFE methods have demonstrated their power in addressing complex challenges across various projects.^4,34,39,40,44–48^ Several tools for alchemical ABFE calculations have been proposed recently including ABFE workflow,^49^ A3FE,^50^ BAT2^51^ (successor to BAT.py^23^), SAFEP,^52^ FEP+,^46^ BFEE2^53,54^ (successor to BFEE^55^), HTBAC,^56^ XFEP,^17^ CHARMM-GUI,^29^ and FEPSetup. ^57^ Since such tools automate significant portions of the binding free energy calculation pipeline, large-scale comparisons with experimental data have become accessible, reveling that alchemical ABFE predictions are approaching experimental accuracy.^58^ However, the preparation, execution, and analysis of ABFE simulations remain at times tedious, system-dependent, often reliant on expert intervention, and, in some cases, associated with substantial costs due to software licensing. Fully automating the binding free energy pipeline has the potential to overcome these challenges, enabling high-throughput calculations for modern drug design campaigns while enhancing reproducibility and accessibility.

We aim to close this gap by introducing BindFlow, a free open-source software that automates the entire ABFE pipeline. BindFlow implements two ABFE methods: (i) the endpoint free energy method MM(PB/GB)SA^9^ based on a simulation of the receptor–ligand complex only, also referred to as single trajectory approach, and (ii) a double-decoupling alchemical free energy method that uses thermodynamic integration (TI)^59^ or the multistate Bennett acceptance ratio (MBAR)^60^ frameworks for free energy estimation. BindFlow was inspired by Alibay et al. ^5^ and Ries et al. ^49^. BindFlow is highly user-friendly, provides extensive tutorials and documentation, natively supports the small-molecule force fields GAFF, OpenFF, and Espaloma. It schedules tasks either on a local desktop or to SLURM-based distributed computing environments, while being resilient against rare hardware failures or simulation instabilities. For advanced users, it allows full controls over the pipeline and MD settings, and it provides various options of further customization.

Below, we first present implementation principles of BindFlow and concepts of the BindFlow API (application programming interface). Next, we validate BindFlow by computing ABFEs for 139 diverse ligands binding to eight different receptors including six soluble proteins, one membrane protein, and one host–guest system (Fig. 1b). The calculations include, both, FEP and MM(PB/GB)SA methods using three widely used open-source small-molecule force fields: GAFF-2.11, ^61^ OpenFF-2.0.0, ^62^ and Espaloma-0.3.1, ^63^ referred to GAFF, OpenFF, and Espaloma in the following.

**Figure 1.**
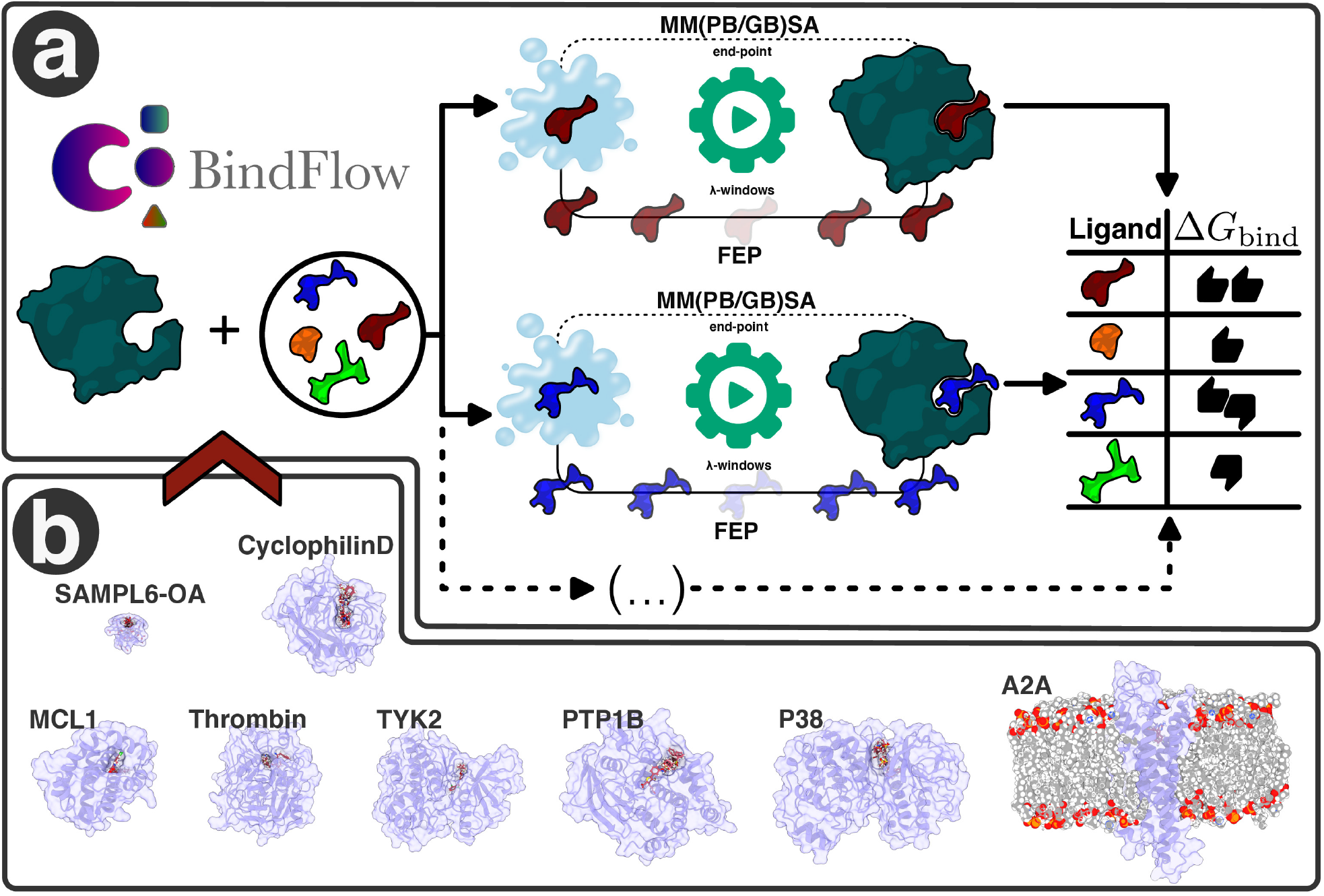
(**a**) Schematic representation of the BindFlow workflow and (**b**) the systems used in this study for validating ABFE calculations.

Overall, our results show that BindFlow delivers predictive performance on par with established gold standards in the field. Strikingly, we observe that for certain targets and force fields, the computationally efficient MM(PB/GB)SA method achieves correlations with experiment that approach those of the more rigorous FEP calculations—yet at only a fraction of the computational cost. By fully automating the binding free energy workflow and minimizing user intervention, BindFlow enhances not only efficiency but also reproducibility. Together, these features make BindFlow a practical, accessible, and reliable platform for large-scale binding affinity prediction, with strong potential for accelerating modern drug discovery campaigns.

### Theory

The theory of FEP and MM(PB/GB)SA have been reviewed frequently,^9,10,12,60,64–66^ hence we provide only a brief summary of the underlying concepts.

### MM(PB/GB)SA

Following the thermodynamic cycle in Fig. 2a, MM(PB/GB)SA estimates the binding free energy Δ*G*_bind_ as follows: ^9,10,65,66^

**Figure 2.**
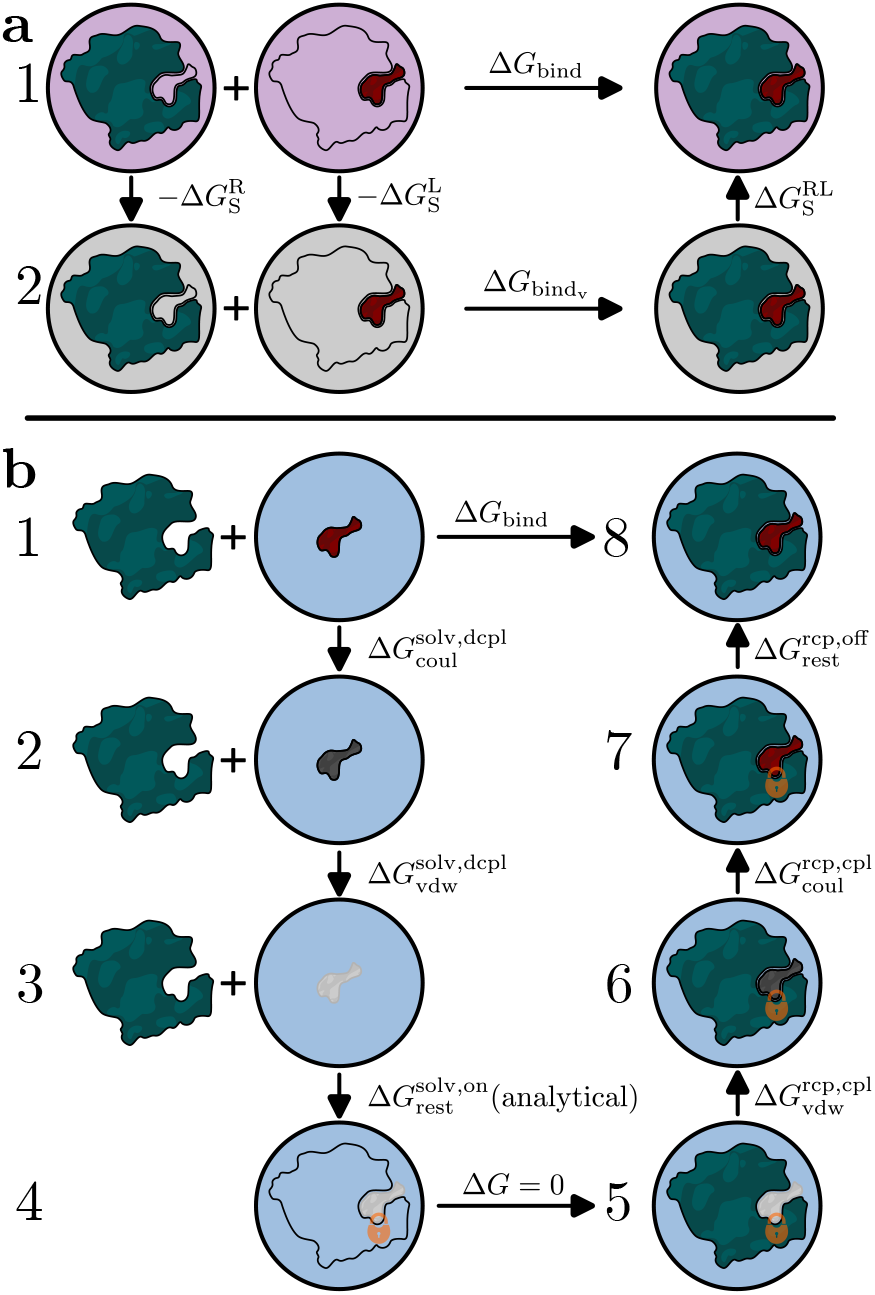
Thermodynamic cycles for (**a**) MM(PB/GB)SA and (**b**) FEP. 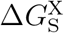 denotes the solvation free energy for species X, where superscripts R, L, and RL denote the receptor, ligand, and receptor–ligand complex, respectively. 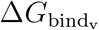 is the binding free energy in vacuum. Superscripts “solv” and “rcp” indicate states with the ligand in solvent or in the receptor, respectively. indicates. Superscripts “dcpl” and “cpl” indicate decoupling or coupling processes, while “coul” and “vdw” specify transitions of Coulomb or Lennard-Jones interactions. Superscripts “on/off” specify the activation or deactivation of Boresch restraints (subscript “rest”). The light pink, gray, and light blue backgrounds represent the implicit solvation model, vacuum, and explicit water model, respectively.

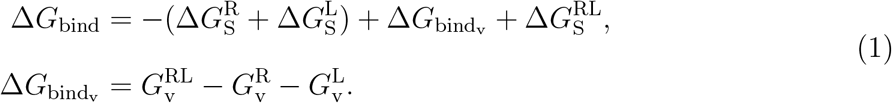

Here, 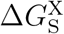 denotes the solvation free energy for species X (X = R, L, or RL), where superscripts R, L, and RL denote receptor, ligand, and receptor–ligand complex, respectively. The solvation free energy for species X is decomposed into polar and non-polar terms, 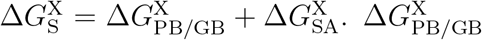 is obtained by solving the Poisson-Boltzmann (PB) equation or using the generalized Born (GB) model, ^10^ while 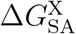 is derived from the solvent-accessible surface area (SASA).^67^

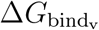 denotes the binding free energy in vacuum, computed from the free energies 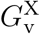 of receptor, ligand, or receptor–ligand complex. The free energies are estimated via

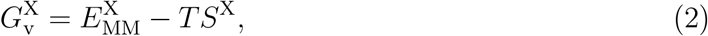

where 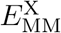 is the molecular mechanics (MM) potential energy of species X comprising bonded and non-bonded terms, as defined by the MM force field. *T* is the temperature, and *S*^X^ is the entropy of species X. By defining the free energy of species X as

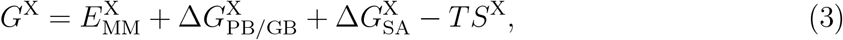

Δ*G*_bind_ may be written as the free energy difference between products and reactants,

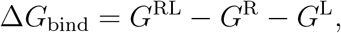

thereby rationalizing by the subscripts of Eq. 3 the name MM(PB/GB)SA.^9,10,65,66^

BindFlow computes the energy contributions to Δ*G*_bind_ by averaging over multiple MD simulation frames using the package gmx_MMPBSA.^65^ The entropic term may be estimated using normal mode analysis, interaction entropy (IE),^68^ or a cumulant approximation to the second order of the exponential average (C2). ^69^ These entropy estimations have been implemented by gmx_MMPBSA^65^ and may therefore be used by BindFlow.

BindFlow employs a single-trajectory protocol, that is, all contributions from ligand, receptor, or receptor–ligand complex are computed from MD simulations of the complex. This approach reduces the computational cost and facilitates the cancellation of bonded terms in 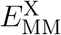. However, the approach neglects contributions from structural changes of the ligand or receptor upon binding.^10,65^

### Free energy perturbation (FEP)

Alchemical FEP estimates binding free energies by constructing a thermodynamic cycle. Such cycles involve decoupling the ligand from solvent (Fig. 2b, Steps 1–3) and coupling it back in the binding site (Fig. 2b, Steps 5–7) in the presence of restraints. The restraints maintain the ligand in the binding pocket, thereby improving sampling and unambiguously defining the reference standard state. BindFlow employs Boresch restraints. ^38^ The free energy for activating Boresch restraints in the solvent are calculated analytically (Fig. 2b, Steps 3–4) and the free energy for removing the restraints in the receptor environment are derived numerically (Fig. 2b, Steps 7–8).

Free energy differences Δ*G* between two states (e.g., between states 2 and 1 in Fig. 2b) are estimated from simulations across a series of intermediate alchemical states, specified by a parameter *λ*. Bindflow calculates Δ*G* values using either thermodynamic integration (TI)^64^ or the multistate Bennett acceptance ratio (MBAR)^60^ method. TI computes the Δ*G* values by integrating the mean work along the *λ* parameter:^64^

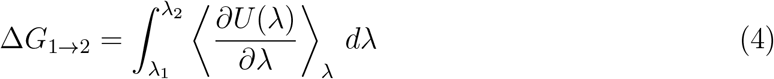

where ⟨… ⟩_*λ*_ denotes the ensemble average at a given value *λ* and *U* (*λ*) is the potential energy. In practice, the integral is solved numerically using a finite number of *λ* points. MBAR estimates Δ*G* values using information from all alchemical states. MBAR requires solving the following set of equations self-consistently to obtain the reduced free energies *f*_*i*_:^60^

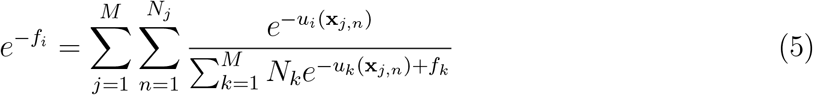

Here, *M* is the number of *λ*-states, *N*_*j*_ is the number of samples from the *λ*-state *j*, and *u*_*i*_(**x**_*j,n*_) is the reduced potential energy of the *n*^th^ sample **x**_*j,n*_ from the simulation of *λ*-state *j* evaluated with the energy function of *λ*-state *i*.

### Implementation

#### General concept of BindFlow

As illustrated in Fig. 3, BindFlow streams the entire binding free energy pipeline, including building of the simulations systems, definition of force fields and MD parameters, careful multi-step equilibration routines, launching of productions simulations, and analysis. For FEP calculations, specifically, production simulations include the definition of Boresch restraints as well as the setup, equilibration, and launching of all *λ*-windows. The pipeline is organized into tasks with well-defined dependencies, which are deployed to the computing environment. The environment may be a desktop computer, a parallelized high-performance computer, or another distributed computing environment. Task scheduling is carried out by Snakemake,^70^ a robust task manager that has been widely used for defining complex bioinformatics analyses, but has to our knowledge been less used in the MD community. Snakemake constructs a direct acyclic graph (DAG), which specifies the dependencies among Bindflow tasks to be executed asynchronously, thereby optimally using the available hardware. Bind-Flow uses GROMACS^71^ as molecular dynamics engine.

**Figure 3.**
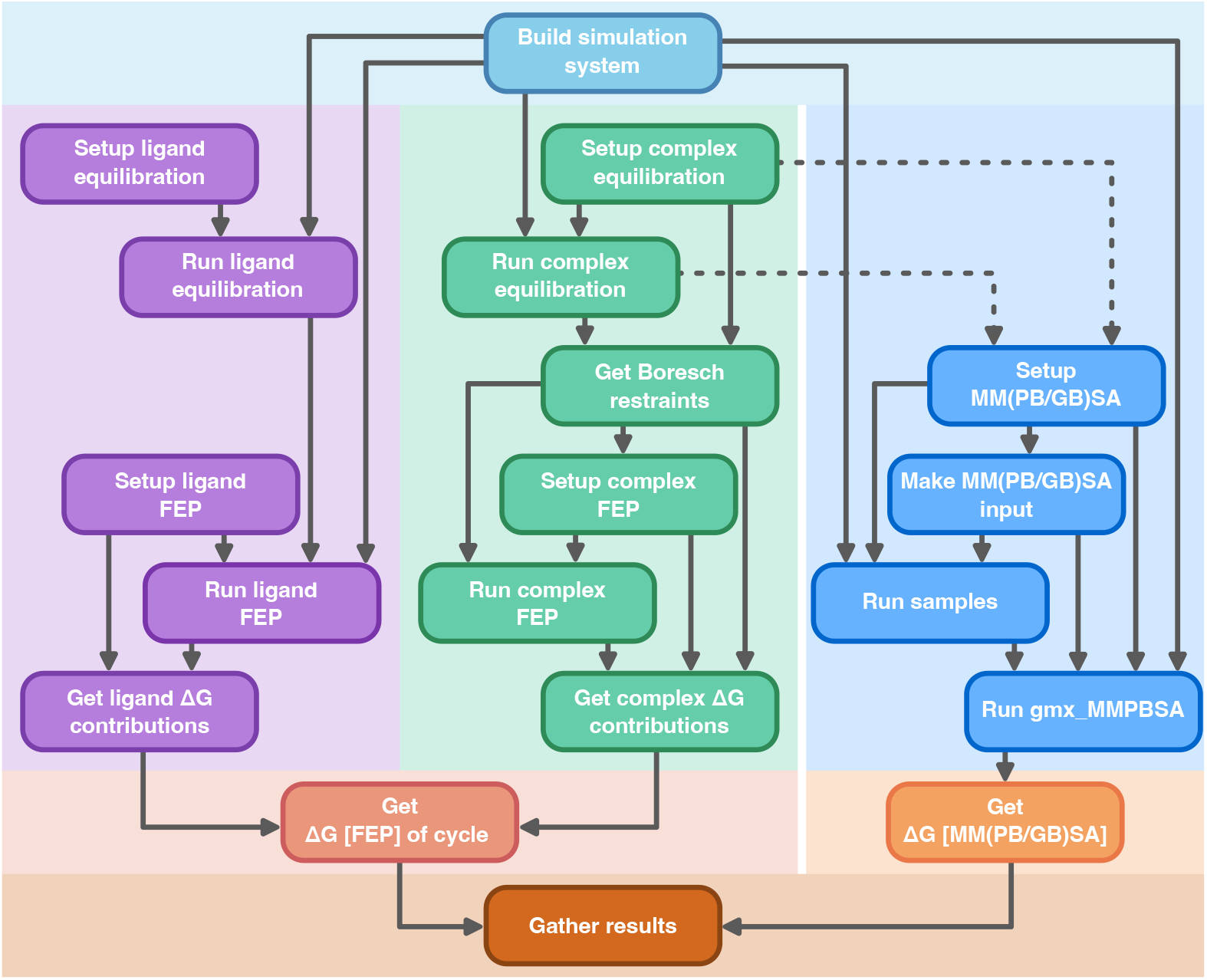
Task dependency graph for BindFlow. Each rectangle represents a task (or group of tasks), while arrows indicate dependencies between them. Some tasks are shared between methodologies. Purple boxes correspond to ligand-in-solvent FEP simulations, and green boxes to protein–ligand complex FEP simulations. The darkest blue boxes denote MM(PB/GB)SA simulations, which reuse some tasks from the protein–ligand complex FEP workflow. All three approaches share the initial system-building step (light blue) and converge at the final result-gathering stage to report the binding free energy, Δ*G*_bind_.

BindFlow may compute Δ*G*_bind_ from a protein PDB file together with a set of ligands provided as MOL files. However, for more complex systems including cofactors or membranes, additional input files may be provided.

The pipeline is implemented into a single function *calculate* within the module *runners*. The function *calculate* takes general parameters such as the protein and ligand structures and, through the keyword argument *global_config*, more specific definitions such as the computational environment, or parameters for simulations analysis. Listing 1 presents a minimalistic Python script for running an FEP pipeline in an HPC environment, while Listing S1 presents a more extended example for running an MM(PB/GB)SA pipeline on a desktop computer. Parameters passed with *global_config* may be specified within Python (Listing 1) or, more conveniently, using JSON or YAML files (Listing S1). While BindFlow implements default workflows, the user has full controls over equilibration and production settings, GROMACS parameters, and task deployment (Listing S7). Full documentation is provided online. Two classes for deployment are provided, namely for a desktop computer and for the SLURM queuing system,^72^ however, alternative deployment managers may be added by the user via the abstract base class (ABC) *Scheduler*, for instance for using cloud computing services. Thus, BindFlow is highly user-friendly, yet allows extensive customization by the user.

**Listing 1:**
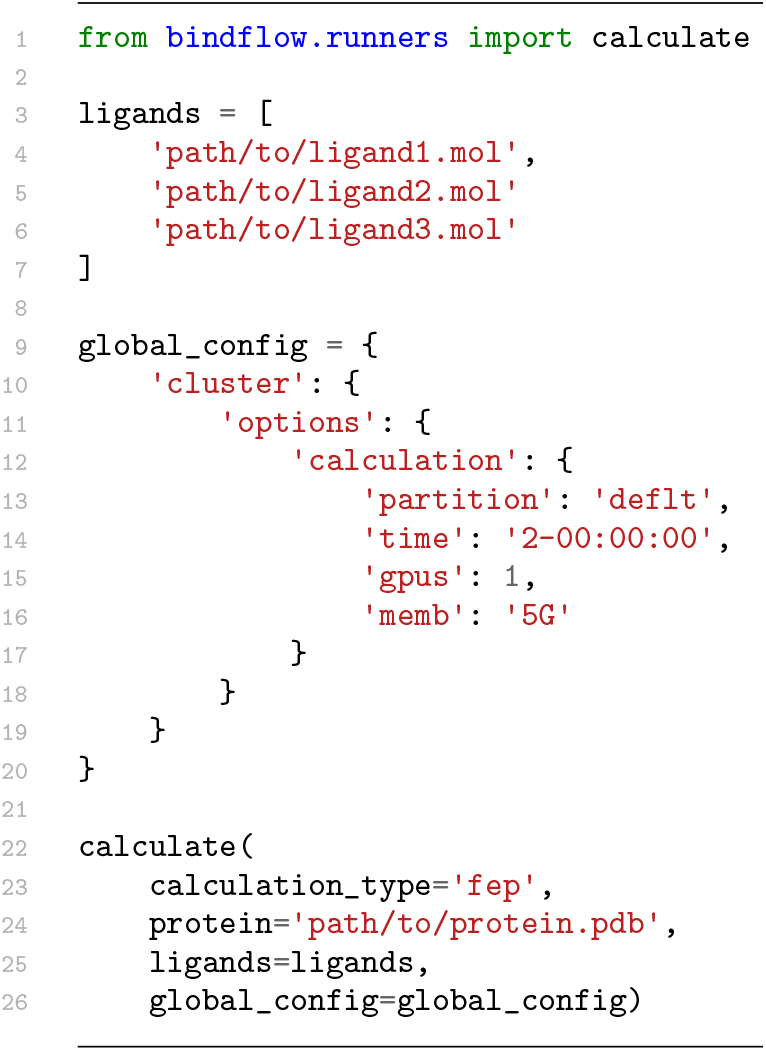
Minimal Python code for running an FEP campaign with BindFlow, as specified by *calculation type*. The *protein* specifies the path to the protein PDB file, and the *ligands* the list of ligand MOL files. The dictionary *global_config* specifies options for the computing environment.

BindFlow has been forked from ABFE Workflow,^49^ yet has been largely rewritten with the following aims in mind: options for extensive customization, efficient resource utilization, built-in support for several small-molecule force fields, flexibility with respect to user-provided force fields, comprehensive documentation, implementation of MM(PB/GB)SA, handling of cofactors, as well as support for membrane proteins and non-protein receptors. BindFlow is released as Open Source under the GPL-3.0 license.

#### Input files and force fields

For a typical workflow, the protein structure is provided as PDB file and a set of ligands as MOL files, where the ligand coordinates are aligned with the protein binding pocket. Optionally, a cofactor as MOL file or a membrane as PDB file may be provided. Alternatively, to enable the use of custom force fields, the user may provide GROMACS topology (TOP) and structure files (GRO). Running membrane protein systems requires additional care, as described in the online documentation. Examples for system component definitions are given in Listings S2, S3, S4, S5, and S6.

By default, proteins are described with Amber99sb-ildn, ^73^ membranes with SLipids2020,^74^ cofactors and ligands with OpenFF-2.0.0, ^62^ and water molecules with the TIP3P model.^75^ Other protein force fields may be selected from the GROMACS distribution or from a user-specified path. Alternative water models may be chosen. The small-molecule force fields OpenFF,^62^ GAFF,^61^ and Espaloma^63^ are natively supported by BindFlow via the TOFF^76^ package.

## Results

### Validation set involving 139 receptor–ligand pairs and three small-molecule force fields

We validated BindFlow by computing the binding affinity of 139 ligand/receptor pairs, for which high-quality experimental affinity data is available (Fig. 1b). The receptors included six soluble proteins (Cyclophilin D, MCL1, Thrombin, TYK2, PTP1B, P38), the transmembrane GPCR protein A2A, and the non-protein host–guest system SAMPL6-OA that has been used for a binding affinity prediction challenge.^78^ Affinity data for P38, PTP1B, TYK2, Thrombin, and MCL1 have been widely utilized for validating RBFE^15–18^ and ABFE calculations,^17,46,79^ allowing us to compare BindFlow results with the literature (see below). These systems involve various challenges for binding affinity calculations. (i) Some ligands binding to MCL1, PTP1B, Thrombin, or SAMPL6-OA are charged (Figs. S25, S22, S24, S27). (ii) Ligand binding to PTP1B or P38 may proceed via slow induced-fit conformational transitions, suggesting that simulations may not sufficiency sample the apo state and, thereby, partly miss the free energy cost for the induced-fit transition.^46^ (iii) Binding affinities may be influenced by cofactors: for P38, PTP1B, TYK2, and Thrombin, crystallographic water molecules may interact with the ligand (Table 1, 6^th^ column). In A2A, a sodium ion interacts with the ligand, suggesting that imprecise modeling of a salt bridge by the force field may add considerable uncertainty. (iv) Among the ten ligands binding to Cyclophilin D (Fig. S26), four comprise more than 30 heavy atoms, which could potentially increase sampling challenges, while (v) eight ligands contain exotic ring structures, posing challenges for the accurate force field representation.^5,49,50^ Thus, our test sets involve highly diverse ligand/receptor pairs that cover various common challenges during binding affinity calculations. All ligands are shown in Figs. S20–S27.

**Table 1:** Simulation systems used for validating BindFlow.

| System | Number of Atoms | Force Field |  | Number of Ligands | Cofactor(s) | Dynamic Range<br>[kcal mol<sup>-1</sup>] | Reference |
| --- | --- | --- | --- | --- | --- | --- | --- |
|  |  | Protein | Ligands |  |  |  |  |
| P38 | 86,376 | Amber14sb | Espaloma-0.3.1<br>GAFF-2.11<br>OpenFF-2.0.0 | 29 | 3H<sub>2</sub>O | 3.80 | Hahn et al.<sup>30</sup> |
| A2A | 84,996 | Amber14sb | Espaloma-0.3.1<br>GAFF-2.11<br>OpenFF-2.0.0 | 10 | Na<sup>+</sup> | 2.69 | Deflorian et al.<sup>13</sup> |
| PTP1B | 74,246 | Amber14sb | Espaloma-0.3.1<br>GAFF-2.11<br>OpenFF-2.0.0 | 22 | 4H<sub>2</sub>O | 5.17 | Hahn et al.<sup>30</sup> |
| TYK2 | 66,425 | Amber14sb | Espaloma-0.3.1<br>GAFF-2.11<br>OpenFF-2.0.0 | 13 | 2H<sub>2</sub>O | 3.47 | Hahn et al.<sup>30</sup> |
| Thrombin | 49,471 | Amber14sb | Espaloma-0.3.1<br>GAFF-2.11<br>OpenFF-2.0.0 | 23 | 3H<sub>2</sub>O | 5.87 | Hahn et al.<sup>30</sup> |
| MCL1 | 34,829 | Amber14sb | Espaloma-0.3.1<br>GAFF-2.11<br>OpenFF-2.0.0 | 25 | None | 4.19 | Hahn et al.<sup>30</sup> |
| Cyclophilin D | 31,861 | Amber99sb-ildn | Espaloma-0.3.1<br>GAFF-2.11<br>OpenFF-2.0.0 | 10 | None | 8.49 | Alibay et al.<sup>5</sup>, Ries et al.<sup>49</sup> |
| SAMPL6-OA | 9,204 | Espaloma-0.3.1 | Espaloma-0.3.1 | 7 | None | 3.79 | Rizzi et al.<sup>77</sup>, Isik et al.<sup>78</sup> |

We computed binding for these 139 ligand/receptor pairs in triplicates using FEP or MM(PB/GB)SA, except for the Cyclophilin D set, where five FEP replicates were used instead. For all protein targets, we computed binding affinities using three popular small-molecule force fields: GAFF-2.11,^61^ OpenFF-2.0.0, ^62^ and Espaloma-0.3.1.^63^ For the SAMPL6-OA system, only Espaloma-0.3.1 was used. ^63^ Cyclophilin D was described with the Amber99sbildn force field to allow comparison with a previous study. ^5,49,50^ Among our MM(PB/GB)SA calculations, we compared results from the Poisson-Boltzmann with results from the generalized Born model, and, for each solvation model, evaluated the effects from using either no entropy correction or using the IE or C2 entropy correction. Together, our validation sets comprise 1215 FEP calculations and 7254 MM(PB/GB)SA calculations.

We quantified the agreement between calculated Δ*G*_calc_ and experimental Δ*G*_exp_ affinities using the Pearson *ρ*, Kendall *τ*, and Spearman *r*_S_ correlation coefficient. Here, Pearson *ρ* quantifies the linear correlation between calculated and experimental values, accounting for magnitudes rather than just ranks, though it remains sensitive to outliers. In contrast, Kendall *τ* and Spearman *r*_*S*_ quantify agreement in ranking and are far less sensitive to outliers. Whereas Spearman *r*_S_ quantifies whether the ligand ranking according to Δ*G*_exp_ agrees with the ligand ranking according to Δ*G*_calc_, Kendall *τ* quantifies the concordance between pairs of ligands. Upon comparing correlation coefficients from different receptors, it is critical to note that Pearson *ρ* is highly sensitive to the dynamic range among the ligands, whereas Kendall *τ* and Spearman *r*_*S*_ are less sensitive to the dynamic range (Table 1, 7^th^ column). In this study, we strongly rely on Kendall *τ* as key quality measure for Δ*G*_bind_ calculations since it has been shown to be a robust estimator, ^80^ while reporting Pearson *ρ* and Spearman *r*_S_ as complementary measures. In addition to the correlation coefficients, we computed the root mean-squared error (RMSE), mean signed error (MSE), and mean unsigned error (MUE) of Δ*G*_calc_ relative to Δ*G*_exp_ (see SI methods). To focus on the global correlation between Δ*G*_calc_ and Δ*G*_exp_ across all ligand-receptor pairs, we furthermore subtracted from each set its corresponding MSE from Δ*G*_calc_ to yield offset-corrected 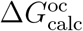 values and, respectively, offset-corrected RMSE (ocRMSE) and offset-corrected MUE (ocMUE, see Methods). Thereby, ocRMSE and ocMUE ignore systematic offsets between Δ*G*_calc_ and Δ*G*_exp_, for instance owing to insufficiently sampling of the apo state or owing to a systematic bias in protein–ligand interactions as modeled by the force field. ocRMSE and ocMUE values may be compared with results from RBFE calculations since the latter are blind to such systematic offsets. Uncertainties of the statistical measures were derived by bootstrapping among the ligands for each set. Thus, critically, the errors bars for statistical measures are not only caused by limited sampling but furthermore stem from the limited number of 7 to 29 ligands per receptor.

### Comparison of free energy perturbation (FEP) results with experiments

Figure 4 presents correlation plots between offset-corrected calculated affinities 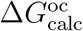 ob-tained with GAFF and experimental affinities Δ*G*_exp_. Values are either listed for seven protein targets individually (small panels) or accumulated from these seven targets (large panel). Insets report Pearson *ρ*, Kendall *τ*, ocMUE, and ocRMSE for each data set. Correlations plots prior to the offset correction are shown in Figs. S2 and S3. Correlation plots obtained with OpenFF or Espaloma are shown in Figs. S6 and S7. Notably, we removed a single extreme outlier given by the ligand lig 4 binding to Thrombin (Fig. S24) from the statistical quality measures to avoid a bias from a single receptor–ligand pair (see Fig. S3). Lig 4 comprises a cationic amidinium moiety 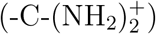 forming a salt bridge to Asp-189 of Thrombin (Fig. S28). We speculate that the greatly overestimated affinity of lig 4 may be a consequence of overly stable amidinium–carboxylate bridge modeled by the three force fields.

For most receptors and force fields, the affinities were overestimated, as shown by mostly negative MSE values between typically −4 and 0 kcal mol^−1^ (Figs. S2, S3, see also Fig. 5d). Such negative MSE values have been rationalized by insufficient sampling of the apo state, thereby partly missing the free energy cost of the induced-fit conformations transition upon ligand binding.^46^ An extreme case was PTP1B with MSE values between −14 and −10 kcal mol^−1^, pointing to a major free energy cost for the induced-fit conformational transition that was not captured during FEP simulation, in line with the value of approximately −10 kcal mol^−1^ reported by Chen et al. ^46^.

**Figure 4.**
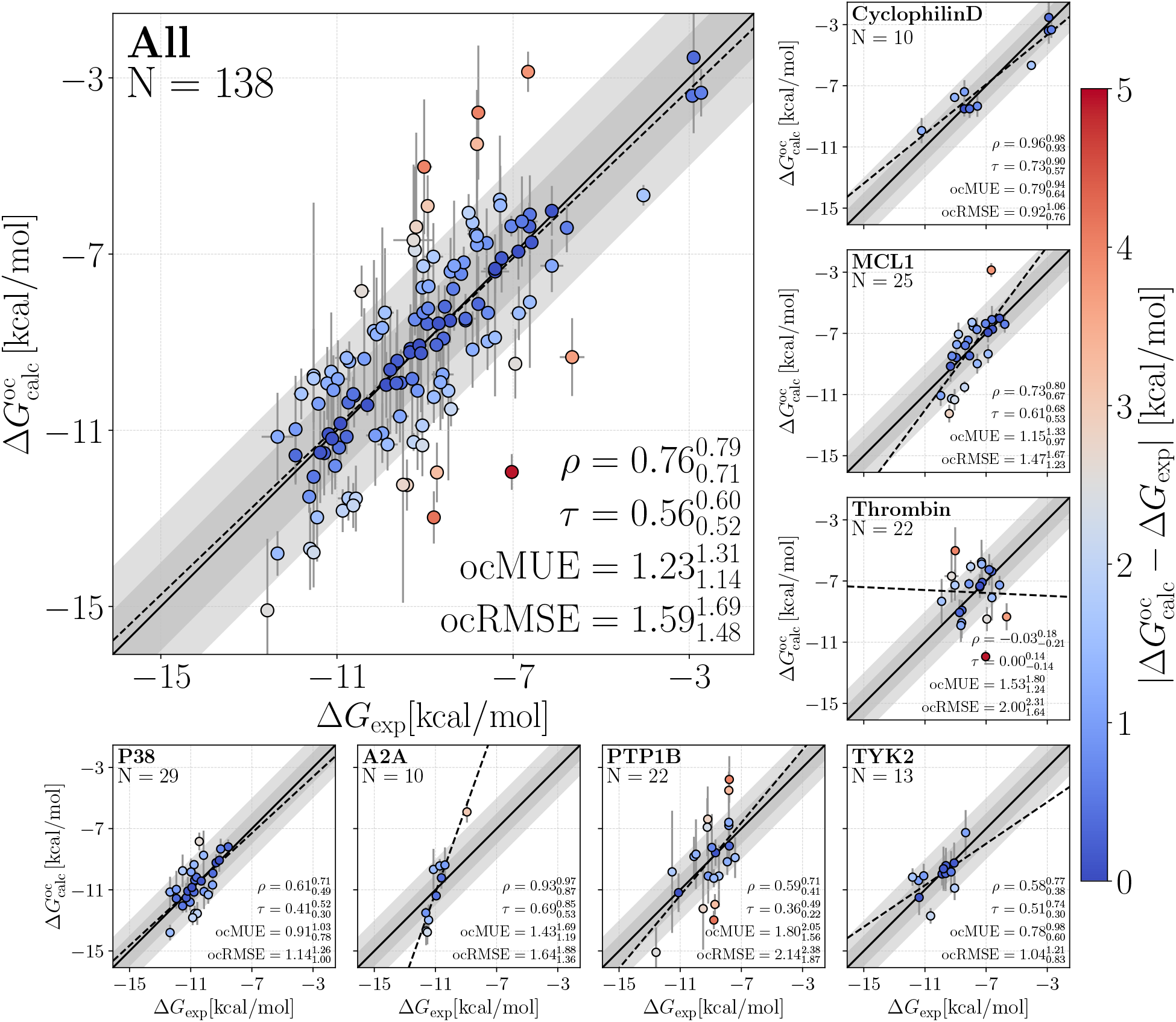
Offset-corrected calculated affinities 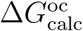 versus experimental affinities Δ*G*_exp_ from FEP with GAFF-2.11. Results shown for individual set (small panels, see labels for protein name and number of ligands *N*) and collected from all sets (large panel). Insets show Pearson *ρ*, Kendall *τ*, osMUE, and ocRMSE (last two in kcal mol^−1^) for each data set with its corresponding 68 % confident interval. Colors of dots indicate the absolute deviation between Δ*G*_calc_ and Δ*G*_exp_ (color bar). Dark and light gray diagonal regions indicate 1 or 2 kcal mol^−1^ deviations, respectively. Dashed lines are linear fits shown to guide the eye. A single outlier (lig 4, Thrombin) has been removed. Error bars show uncertainties obtained via three independent replicates. Figure style following Ref. 79.

**Figure 5.**
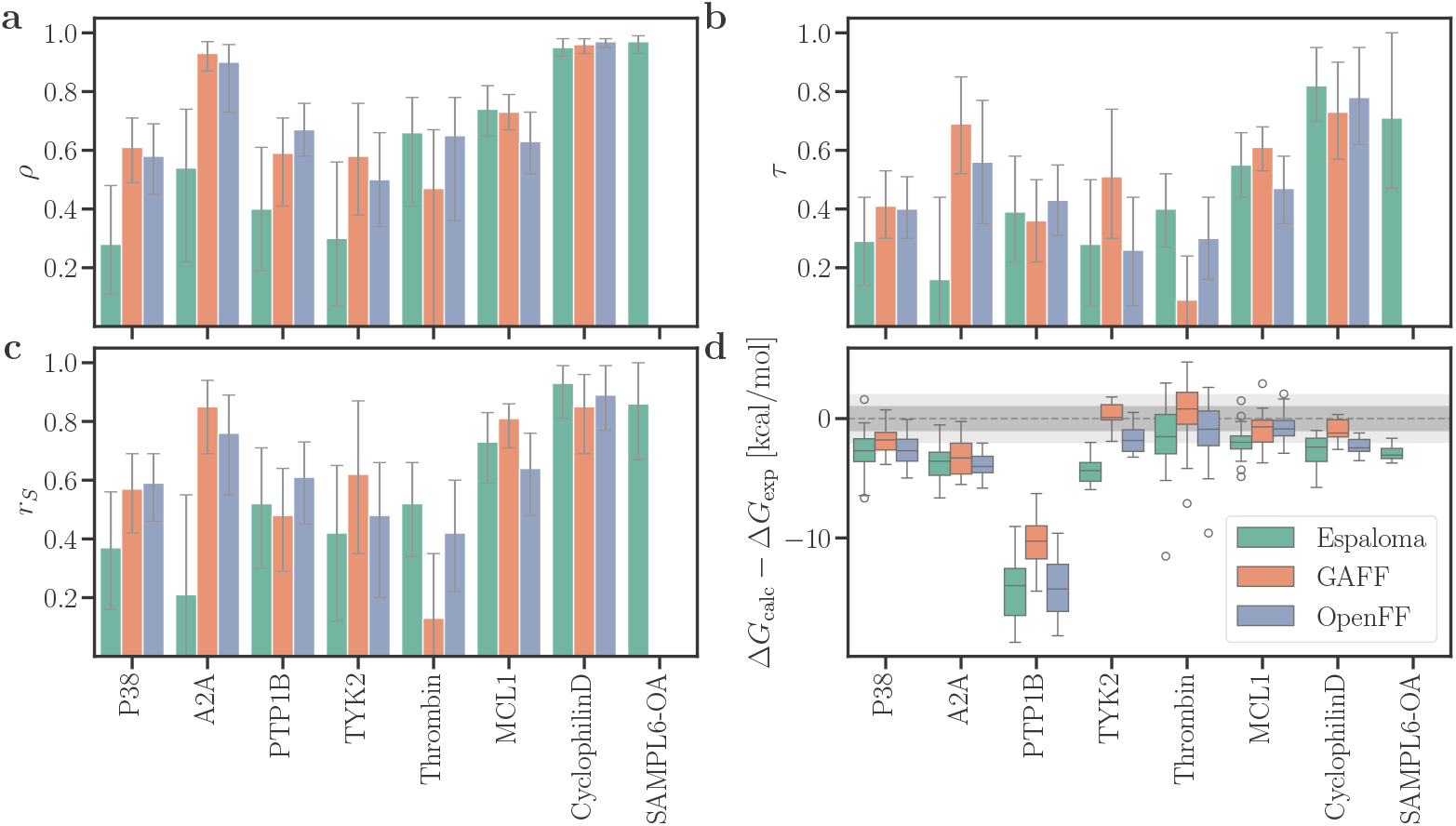
(**a**) Pearson *ρ*, (**b**) Kendall *τ*, (**c**) Spearman *r*_*S*_, and (**d**) deviations between calculated and experimental binding free energies Δ*G*_calc_ − Δ*G*_exp_ from FEP using Espaloma-0.3.1 (green), GAFF-2.11 (orange), and OpenFF-2.0.0 (blue). Results are shown for seven different data sets (labels at abscissa). Error bars represent the 68 % confident interval obtained from bootstrapping among the ligands from each set. For the Δ*G*_calc_ − Δ*G*_exp_, dark and light gray horizontal regions indicate 1 or 2 kcal mol^−1^ deviations, respectively. Box plots present the median (50^th^ percentile) as a line within the box, and the lower and upper box edges correspond to the first (25^th^ percentile) and third (75^th^ percentile) quartiles. Whiskers extend to the smallest and largest data points within 1.5 *×* interquartile range (IQR = Q3 − Q1), while outliers beyond this range are shown as circles.

After removing the MSE form Δ*G*_calc_ to obtain 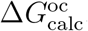, statistical quality measures for the agreement with experiments still greatly depended on the protein target, with ocMUE values far below 1 kcal mol^−1^ for Cyclophilin D up to approximately 2 kcal mol^−1^ for PTP1B. Excellent agreement was found for SAMPL6-OA (Fig. S7). These variations were likewise reflected by Kendall *τ* which spanned 0.82 for Cyclophilin D with Espaloma down to poor *τ* for Thrombin. Pearson *ρ* and ocRMSE values reported similar trends. The wide range of correlations obtained with different receptor and different force fields are summarized by Fig. 5.

### Comparison of FEP results from BindFlow with the literature

Ross et al. ^81^ evaluated the reproducibility of experimental RBFE and reported an overall RMSE (weighted by number of ligands on each set) of 0.91 kcal mol^−1^ (95% CI: [0.83, 1.11]) and a weighted average Kendall *τ* = 0.71 (95% CI: [0.65, 0.74]). Thus, upon comparing Δ*G*_calc_ with Δ*G*_exp_, the considerable intrinsic uncertainty of the Δ*G*_exp_ must be kept in mind. Accordingly, an RMSE of ∼ 1 kcal mol^−1^ and Kendall *τ* of ∼ 0.7 between Δ*G*_calc_ and Δ*G*_exp_ would indicate excellent agreement.

In addition, Ross et al. ^81^ validated the FEP+ software for RBFE calculations in combination with the OPLS4 force field^82^ against protein–ligand sets different from the ones used here. Beyond the differences in validation sets, their study employed RBFEs, whereas we performed ABFEs, which are generally more challenging. For these reasons, a direct, quantitative comparison of RMSE and Kendall *τ* is not meaningful. Nevertheless, placing the results side by side helps to contextualize the current state of the art in FEP calculations. FEP+ achieved an RMSE of 1.25 (95% CI: [1.17, 1.33]) and a Kendall *τ* of 0.51 (95% CI: [0.48, 0.55]). With Espaloma, GAFF, and OpenFF, we obtained slightly higher ocRMSE values, which may indicate that the closed-source OPLS4 force field^82^ performs somewhat better than the currently freely available force fields. However, the Kendall *τ* values obtained here were even slightly higher than those from FEP+, suggesting that our ABFE calculations are competitive in terms of ligand ranking.

Next, we systematically compared ABFE results from BindFlow with FEP+ by restricting the analysis to ligands that were present in both the dataset of Chen et al. ^46^ and our BindFlow simulations, resulting in 89 ligands across the P38, PTP1B, TYK2, and MCL1 sets. Both BindFlow and FEP+ systematically overestimated affinities for PTP1B (Fig. S11, top row). Thus, to enable a quantitative comparison of ABFE values, we removed PTP1B from the datasets and focused the comparison on the remaining 67 ligands. FEP+ yielded systematically stronger binding affinities on these three sets compared to BindFlow, as quantified by the MSE of 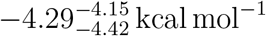 for FEP+, versus 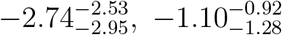, and 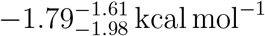 for Espaloma, GAFF, and OpenFF, respectively (Fig. S11, middle row). Keeping the PTP1B set and removing the constant offset for each set by subtracting the set-specific MSE from Δ*G*_calc_, FEP+ yielded a considerably lower ocRMSE and ocMUE values compared to BindFlow when used with any of the three force fields. FEP+ also yielded higher correlation coefficients (Fig. S11, bottom row). Thus, in the light of these data, FEP+ may achieve superior ranking at the cost of overestimating the absolute binding affinity. We speculate that the superior ranking by FEP+ is primarily a consequence of using the OPLS4 force field, although alternative sampling algorithms may also play a role. Within this subset of ligands, GAFF achieved better agreement than Espaloma and OpenFF with experimental data, with 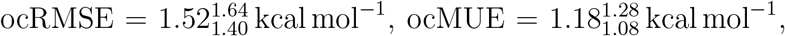, and 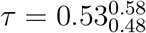 (Fig. S11, bottom row).

Deflorian et al. ^13^ reported RBFE calculations for the A2A system using the FEP+ software. We converted their RBFE values into ABFEs by referencing the experimental binding affinity of ligand 4k and correcting for the MSE. From their data, we ob-tained 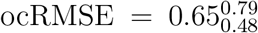 and 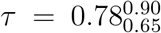. Evidently, since all ligands were similarly composed of three aromatic rings (Fig. S21), RBFE required relatively small perturbations, rationalizing the high *τ* and exceptionally low ocRMSE. Although our ABFE calculations involved by far larger perturbations, we obtained decent agreement (Fig. S14) with GAFF (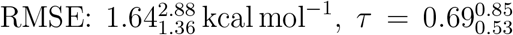, Fig. 4) and with OpenFF (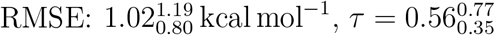, Fig. S6). Only with Espaloma, we obtained a rather poor *τ* of only 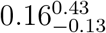 (Fig. S7), which might indicate inaccurate modeling of the ligand interactions with the Na^+^ cofactor inside the A2A binding pocket.

The Cyclophilin D set was first introduced by Alibay et al. ^5^ for ABFE benchmarking and was recently employed to validate ABFE workflow^49^ and A3FE.^50^ Excellent agreement with experiments was found across the three pipelines —ABFE workflow, A3FE, and BindFlow—, underscoring that the Cyclophilin D set was the least challenging receptor–ligand set considered in this study (Figs. S12, 5).

The P38 and TYK2 systems have been used by several authors for validating binding free energy calculations using different methodologies.^17,46,48,79^ Figure S13 compares results from BindFlow with previous studies ^17,46,48,79^. Here, Lin et al. ^17^ and Chen et al. ^46^ achieved the best ranking of ligands, however, at the cost of overestimating the absolute binding free energies as shown by large negative MSE. Among our results, GAFF achieved comparable ranking at a less negative MSE.

Taken together, the reasonable agreement with previous data validates BindFlow’s FEP pipeline and demonstrates competitive agreement with experimental data for a wide range of ligands and receptors.

### Comparison of MM(PB/GB)SA with experiments

MM(PB/GB)SA provides a computationally efficient alternative to FEP. Specifically, the MM(PB/GB)SA pipeline used here employed only ∼3 ns of MD simulations; corresponding to a 180-fold lower computing cost compared to the FEP pipeline used here. However, whereas the MMGBSA calculation from the MD frames is highly efficient, the MMPBSA calculation may take considerable additional computing time owing to the cost of Poisson-Boltzmann calculations. As expected from previous studies, ^10^ absolute binding free energies obtained with MMGBSA (Figs. S4, S5, 6d, S15d, S16d) or MMPBSA (Figs. S17d, S18d, and S19d) revealed poor agreement with experimental values. For instance, MMGBSA without entropy correction overestimated the absolute binding affinities to protein targets by approximately 25 to 75 kcal mol^−1^ (Fig. 6d). Even upon correcting the binding affinity with a target-specific offset, ocRMSE values remained large (Figs. S8, S9, S10).

**Figure 6.**
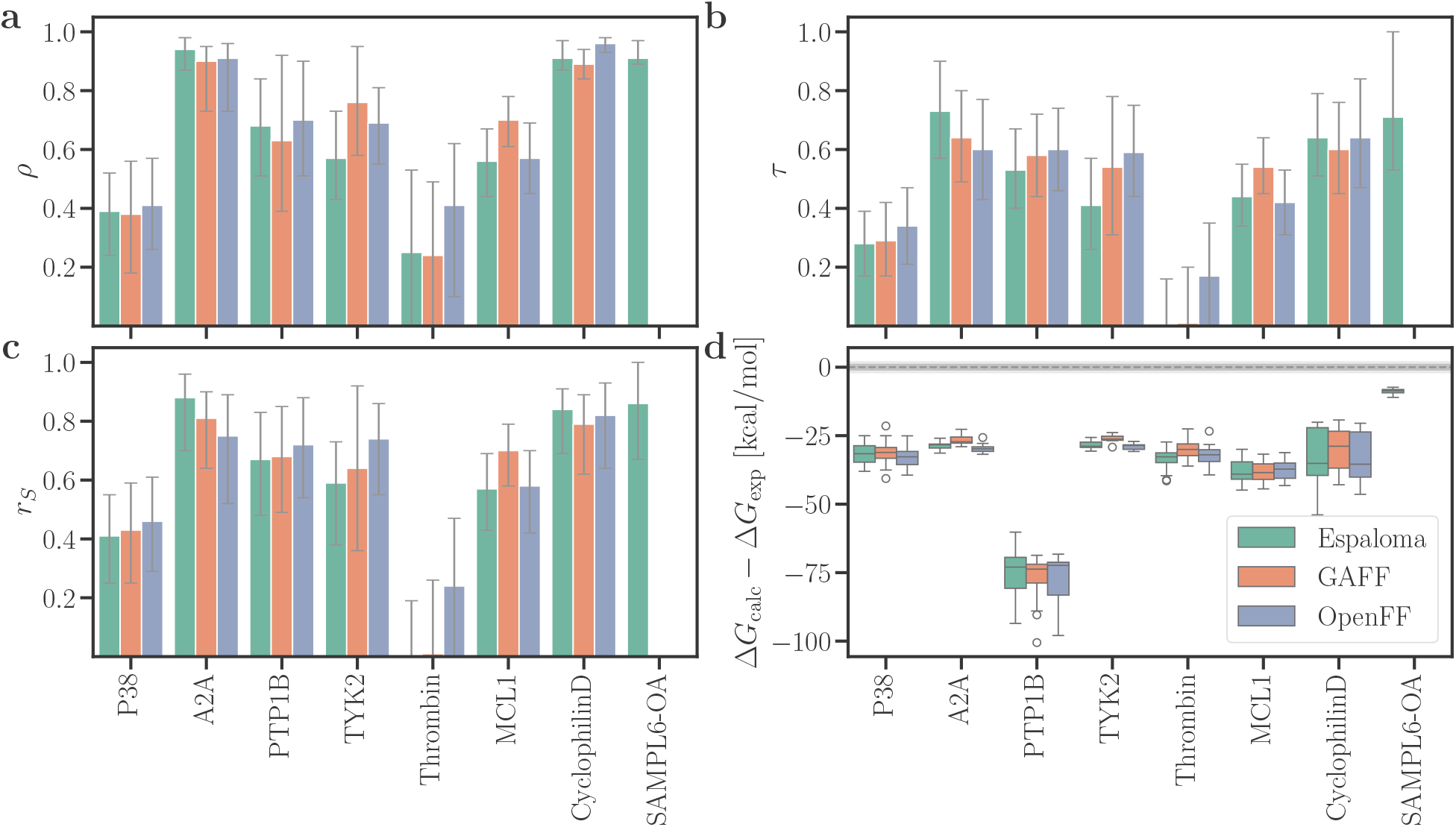
Pearson *ρ*, Kendall *τ*, Spearman *r*_*S*_, and deviations between calculated and experimental binding free energies (Δ*G*_calc_ − Δ*G*_exp_) from MMGBSA results without entropy correction. Presentation style according to Fig. 5.

However, MMGBSA without entropy correction achieved overall excellent Pearson, Kendall, and Spearman correlation coefficients for A2A, Cyclophilin D, and SAMPL6-OA, while reaching still good correlation coefficients for PTP1B, TYK2, and MCL1 (Fig. 6a–c). In contrast, correlation coefficients for P38 or Thrombin were rather poor. The three force fields GAFF, OpenFF, and Espaloma achieved similar ranking (Figs. S10, S8, S9 and also 6a–c). Thus, while MMGBSA is not suitable for obtaining absolute binding free energies (Fig. 6d), it may provide for certain receptors reasonable ranking among ligands and, thereby, serve as a useful tool for screening large datasets of potential binders.

Using the Poisson-Boltzmann instead of the generalized Born solvation model had only moderate effects on the correlation coefficients (compare Fig. S17 with Fig. 6), suggesting that the computationally more efficient generalized Born model yields a good starting point for computational studies. Using the IE entropy correction had only a small effects on the correlation coefficients (compare Fig. S19 with S16), whereas the C2 entropy correction decreased the correlation coefficients (compare Fig. S18 with S15). Hence, in line with previous findings,^83^ entropy corrections should be used with care or tested for the receptor of interest, because entropy corrections may deteriorate the ranking by MM(PB/GB)SA.

### Ligand ranking by FEP versus MM(PB/GB)SA and by different force fields

Figure 7 summarizes the overall performance of FEP versus MM(PB/GB)SA among the three force fields, as collected from all ligands–receptor pairs, thus including easy receptors such as Cyclophilin D and challenging receptors such as Thrombin (see also Table S1). Here, the matrices in Figs. 7b/d visualize whether one combination of method/force field, such FEP/Espaloma, outperforms another combination, such as MMGBSA/GAFF (see Methods for the applied significance test). The following conclusions are drawn from the analysis: (i) In terms of ocMUE, FEP by far outperformed MMGBSA (Figs. 7d). These findings suggest that errors caused by major approximations underlying MMGBSA remain after correcting Δ*G*_calc_ by the MSE. (ii) In terms of ranking by Kendall *τ*, FEP overall outperformed MMG-BSA (Figs. 7a/b). An exception was given by FEP/Espaloma that showed only insignificant differences relative to MMGBSA (Figs. 7b). (iii) For a given method (FEP or MMGBSA), the three force field showed no statistically significant differences (Figs. 7b/d), except among the poor ocMUE by MMGBSA. Thus, simulations with additional ligands will be required to test whether the slightly lower *τ* and slightly larger ocMUE obtained with Espaloma are significant or whether they were caused by the limited set of receptor–ligand pairs considered in this study.

**Figure 7.**
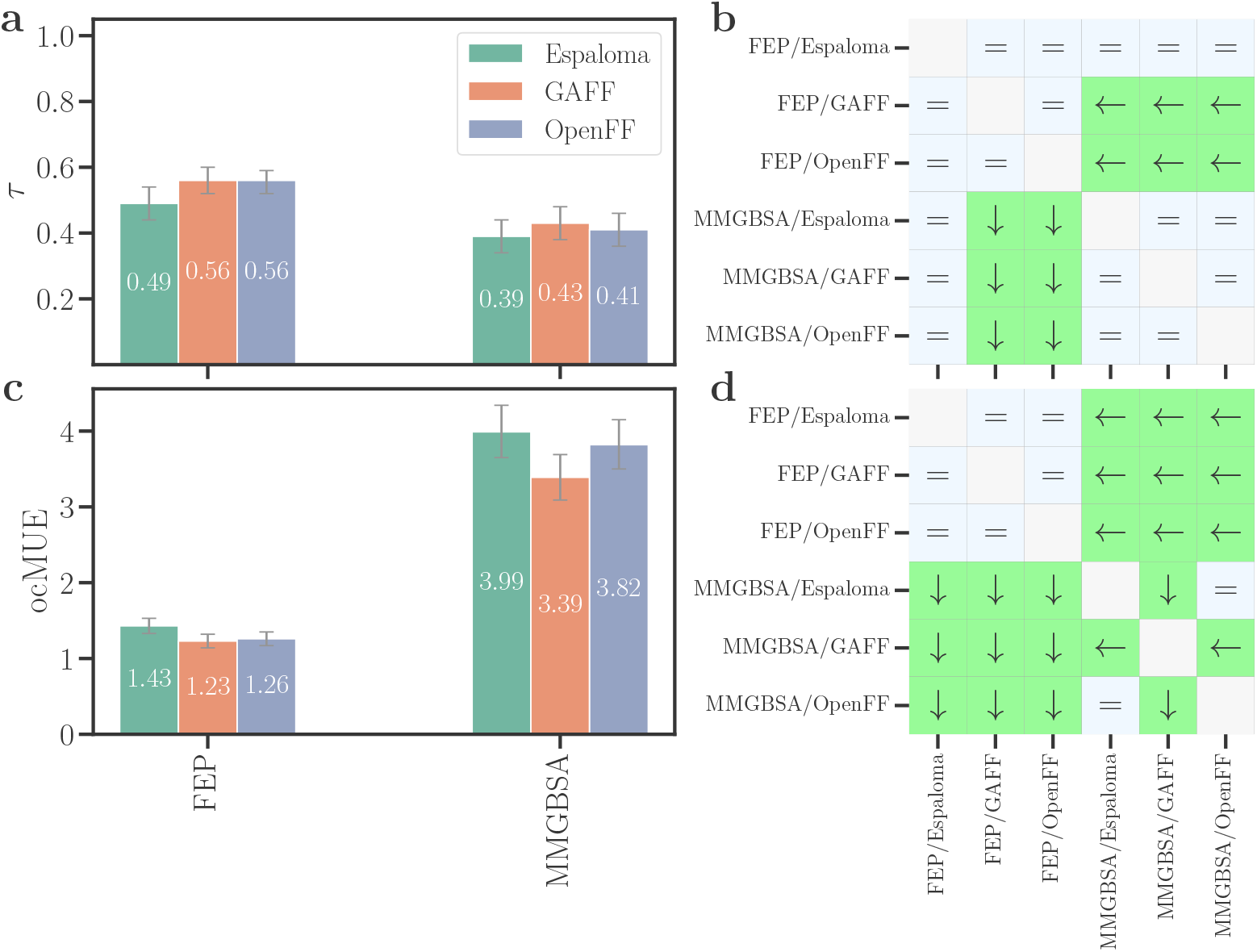
Summary of agreement between FEP or MMGBSA calculations and experimental data quantified by (**a**/**b**) Kendall *τ* and (**b**/**c**) offset-corrected mean unsigned error (ocMUE) in kcal mol^−1^, summarized from all receptors. Error bars show 68 % confident intervals obtained from bootstrapping among the ligands from all sets. Pairwise significance difference matrices are for (**b**) *τ* and (**d**) ocMUE. An arrow indicates a statistically significant difference pointing to the combination of method/force field with the better statistical metric. Equal symbols indicate an insignificant difference. The set SAMPL6-OA is excluded to ensure consistence in the analysis across force fields. The extreme outlier lig 4 from the Thrombin set was excluded.

## Discussion

We introduced BindFlow, a free and user-friendly software for absolute binding free energy (ABFE) calculations. BindFlow offers fully automated pipelines for FEP and MM(GB/PB)SA, requiring minimal user intervention while providing extensive configuration options for advanced users. Comprehensive online tutorials and documentation support accessibility. The software is developed according to best coding practices, ensuring transparency, reproducibility, and a solid foundation for future community-driven development.

Beyond the benchmark simulations presented here, BindFlow has recently been applied successfully for combining ∼60.000 MM(GB/PB)SA calculations with Bayesian active learning^41^ as well as for interpreting X-ray crystallographic data from the antiviral drug tecovirimat binding to the phospholipase F13 from monkeypox virus.^43^ These use cases demonstrate that BindFlow is ready for production use.

We validated BindFlow by computing Δ*G*_bind_ values for 139 ligands binding to a one of eight different receptors using FEP together with MBAR or using MM(GB/PB)SA. Overall, we found that BindFlow yields competitive Δ*G*_bind_ compared to the standards in the field. As expected from previous studies, the accuracy of Δ*G*_bind_ values and ligand ranking greatly depend on the target and on the chemical moieties of the specific ligand.

For the 139 receptor–ligand pairs, in terms of accuracy of Δ*G*_bind_ predictions quantified by ocMUE or ocRMSE, FEP by far outperformed MM(GB/PB)SA. In terms of ligand ranking, FEP still outperformed MMGBSA significantly; however, for several receptors studied here, ranking by MMGBSA was remarkably good. Only for P38 and Thrombin, MMGBSA achieved poor ranking. Thus, our study underscores that MMGBSA may provide a computationally inexpensive method for pre-screening large sets of ligands prior to validation by the more expensive FEP. However, future studies will need to establish whether favorable correlations between MM(GB/PB)SA and experiments, as observed here, also hold for non-congeneric sets of ligands.

BindFlow will be useful to systematically test and improve small molecule force fields. For instance, we spotted that the affinity of lig 4 binding to Thrombin was greatly overestimated by 7 to 13 kcal mol^−1^, possibly caused by an overly strong amidinium–aspartate salt bridge (Fig. S28). This interpretation is supported by the fact that larger partial charges used by Espaloma correlated with an even stronger computed affinity. Since amidinium moieties are quite common in drugs, these findings point towards options for further force field refinements.

Future implementations may further enhance the capabilities of BindFlow. (i) Enhanced sampling techniques, such as Hamiltonian replica exchange, ^84–86^ may accelerate convergence and, thereby, reduce the computational cost. Enhanced sampling may, specifically, improve the sampling of the apo state and lead to smaller MSE values. (ii) The adaptive allocation of computational resources may focus sampling to *λ* windows with increased sampling challenges.^50^ (iii) Charged ligand systems (e.g., MCL1, Thrombin, and PTP1B) showed satisfactory accuracy in our validations. However, finite-size effects in charge-imbalanced systems, primarily due to the use of particle-mesh Ewald (PME) method,^87,88^ remain a concern.^89,90^ Charge corrections ^91,92^ such as the co-alchemical ion approach^12^ represent viable solutions to address this issue. (iv) Water exchange between the binding site and bulk solvent represents a slow sampling process that is difficult to capture during the short MD simulations performed in FEP.^12^ Monte Carlo-based water swap methods in a grand canonical ensemble,^93,94^ methods based on inhomogeneous fluid solvation theory,^95^ or specialized hydration-shell generators such as SOLVATE^96,97^ have shown promise in treating the explicit water. Future versions of BindFlow may include such methodologies.

## Conclusions

We presented BindFlow, an open-source and user-friendly pipeline for absolute binding free energy (ABFE) calculations using FEP and MM(PB/GB)SA. Validation on 139 receptor– ligand pairs showed that BindFlow achieves predictive performance comparable to standards in the field, while offering full automation and extensive customization. FEP achieved more accurate Δ*G*_bind_ predictions and ligand ranking compared to MM(PB/GB)SA. However, MM(PB/GB)SA provided remarkable ligand rankings for several systems at a fraction of the computational cost, highlighting its value for large-scale computational pre-screening campaigns. Thus, we anticipated that the combination of MM(PB/GB)SA and FEP will be powerful for balancing efficiency and accuracy in future BindFlow applications. By combining automation, flexibility, accessibility, and documentation, BindFlow lowers the entry barrier for routine ABFE calculations and provides a platform for modern drug discovery or for systematically improving small-molecule force fields.

## Methods

### Building simulation systems

Simulations followed largely the current default settings of BindFlow. The setup protocols are fully automated, yet may be easily adapted by the user via the *global input* keyword, which is a Python dictionary. This dictionary may be conveniently defined with a YAML file (Listing S7). The simulations presented here were set up as follows.

The task *Build Simulation System* (Fig. 3) assembles the simulation systems and defines force fields. Starting structures were taken from previous studies. ^5,13,30,49,77,78^ Soluble proteins, SAMPLE6-OA, and ligands were placed in a dodecahedral simulation box, keeping a distance of 1.5 nm between solute and box boundary. The systems were solvated by explicit water with the GROMACS solvate module and approximately 150 mM of NaCl were added, thereby neutralizing the system. A minimum distance of 1 nm between ions and nonsolvent molecules was used. The structure of the membrane protein A2A was taken from Ref. 13. BindFlow does not build simulation systems of membrane proteins. Thus, the A2A system was built using CHARMM-GUI^98^ by placing the protein in a simulation box of a hexagonal prism and by embedding it in a membrane of 172 1-palmitoyl-2-oleoyl-sn-glycero-3-phosphocholine (POPC) lipids.

Ligand interactions were described with Espaloma-0.3.1, ^63^ GAFF-2.11,^61^ or OpenFF-2.0.0.^62^ Ligand topologies were generated within BindFlow using the TOFF Python library. ^76^ Cyclophilin D was described with Amber99sb-ildn ^73^ as used by previous studies.^5,49,50^ All other proteins were described with Amber14sb, ^99^ using OpenMM^100^ and ParmEd^101^ for generating force field topologies. The host molecule octa acid of the SAMPL6-OA system was described with Espaloma-0.3.1^63^ and the topologies were generated with TOFF.^76^ Water was modeled with the TIP3P model^75^ and ion parameters were taken from the definitions provided with Amber99sb-ildn. ^73^

### Simulation parameters

The tasks *Setup Complex Equilibration, Setup Ligand Equilibration, Setup Complex FEP, Setup Ligand FEP*, and *Setup Complex MM(PB/GB)SA* (Fig. 3) define MD parameters and set up directory structures for the initial equilibration, FEP or MM(GB/PB)SA simulations.

All simulations were carried out with GROMACS, version 2022.4. ^71^ The geometry of water acting as cofactor was constrained with LINCS, ^102^ while the geometry of all other water molecules were constrained with SETTLE.^103^ All other bonds were constrained with LINCS^102^ if not stated otherwise. Hydrogen mass repartitioning was used with a mass repartition factor of 2.5. During production simulations a time step of 4 fs was used. Dispersive interactions and short-range repulsion were modeled using a Lennard-Jones potential with a 1 nm cutoff. Electrostatic interactions were calculated with the particle-mesh Ewald^87,88^ (PME) method, applying a real-space cutoff of 1 nm.

The temperature was maintained at 298.15 K. For the initial equilibration of membrane protein–ligand complexes, the temperature is maintained using velocity rescaling (*τ*_*t*_ = 1 ps). Here, the protein and the Na^+^ cofactor were coupled to the same thermostat. During all other simulations, the temperature was maintained using Langevin dynamics (*τ*_*t*_ = 2 ps). The pressure was maintained at 1 bar for membrane protein–ligand complexes using a semi-isotropic stochastic cell rescaling^104^ with a time constant *τ*_*p*_ = 5 ps during the initial equilibration. The same barostat was used during the perturbation phase for membrane protein–ligand complexes, but the pressure was maintained at 1 atm (1.01325 bar) and *τ*_*p*_ = 2 ps was used. For all other simulations the pressure was controlled at 1 atm with the isotropic Berendsen barostat during equilibration (*τ*_*t*_ = 1 ps)^105^ and the Parrinello-Rahman barostat during production (*τ*_*p*_ = 2 ps).^106^

### Initial multi-step equilibration protocol

After minimizing the energy with the steepest-descent algorithm, each system was equilibrated with a multi-step protocol (see SI methods).

### FEP calculations

Δ*G*_bind_ from FEP was computed according the thermodynamic cycle shown in Fig. 2b. Coulomb interactions of the ligand in solvent were decoupled over 11 *λ* points (Fig. 2b, transition 1→2), followed by decoupling of Lennard-Jones interactions over 11 *λ* points (transition 2→3). Both, inter- and intramolecular interactions were decoupled. A 10 ns equilibrium simulation together with the software MDRestraintsGenerator ^107^ were used to obtain the optimal Boresch restraints,^38^ the free energy cost for introducing restraints to the ligand in solvent (transition 3→4), as well as the initial frame for the complex decoupling simulations. Inter- and intramolecular Lennard-Jones interactions of the ligand in the receptor were activated over 21 *λ* points (transition 5→6), followed by 11 *λ* points for activating the Coulomb interactions (transition 6→7). Finally, Boresch restraints were removed for the ligand in the receptor over 11 *λ* points (transition 7→8).

For each *λ* window, the system was energy minimized and equilibrated with a three-step protocol (see SI methods). Each window was simulated for 10 ns using a stochastic dynamics integrator. During simulations that decouple Lennard-Jones interactions, a soft-core potential (*α* = 0.5, *σ* = 0.3, power = 1) was used. Free energy differences were computed with MBAR.^60^ As a control, free energies were additionally computed using thermodynamic integration (TI).^64^ BindFlow reports a warning if results from MBAR and TI differ by more than 0.5 kcal mol^−1^, as such differences may indicate poor convergence or simulation instabilities. Alchemlyb-2.0.0 was used for MBAR and TI evaluations.^108^

The binding free energy is computed via (Fig. 2B):

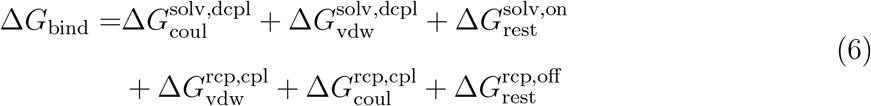

Here, 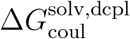 and 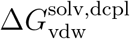 denote free energies for decoupling (dcpl) Coulomb and Lennard-Jones interactions of the ligand in solvent (solv), respectively. 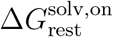 denotes the free energy for turning on Boresh restraints (rest) for the ligand in solvent, which is computed nalytically. 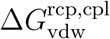 and 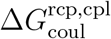 denote the free energy for activating (coupling, cpl) Lennard-Jones and Coulomb interactions for the ligand in the receptor (rcp), respectively. 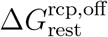 denotes the free energy cost for turning off the Boresch restraints for the ligand in the receptor.

BindFlow reports the statistical error obtained from MBAR ^60^ together with Gaussian error propagation. However, owing to autocorrelations, this value greatly underestimates the true uncertainty of Δ*G*_bind_. Thus, we carried out the whole pipeline including setup, equilibration, and FEP in three (five for Cyclophilin D ^5^) independent replicates and report the respective standard error of the mean as error bars in the correlation plots.

### MM(PB/GB)SA calculations

The initial multi-step equilibration for MM(PB/GB)SA calculation was carried out as described in the SI methods. Following Su et al. ^83^, MM(PB/GB)SA values were computed from multiple short simulations, rather than from a single long simulation. Accordingly, twenty starting frames were taken from a 950 ps equilibrium simulation, using one frame every 50 ps. From each frame, a 100 ps simulation was carried out writing a frame every 5 ps. Here, we follow Genheden and Ryde ^10^, who recommended using an output frequency of 1 to 10 ps. MM(PB/GB)SA values were computed with the gmx_MMPBSA software,^65^ yielding 20 samples of the binding affinity. Cofactors were defined as part of the receptor for the MM(PB/GB)SA calculation. For each complex, the pipeline was run in three independent replicates. Binding affinities reported here were computed by averaging over 60 samples from the three replicates with 20 samples per replica, and the error was derived as SEM and reported as error bars in correlation plots. PB and GB models were used, each without entropy correction or using the IE or C2 entropy corrections. The main text reports results from the GB model without entropy correction, while all other results are provided in the SI.

## Code and data availability statement

BindFlow is free software published under the GPL-3.0 license. The code is currently hosted at https://github.com/ale94mleon/BindFlow. Documentation is available at https://bindflow.readthedocs.io. Scripts and input files required for reproducing our results and analysis are accessible at: https://github.com/ale94mleon/bindflow-api-paper-si.

## Acknowledgement

We thank the authors of ABFE Workflow for publishing the software under a free license, and in particular Benjamin Ries and Aniket Magarkar for helpful discussions. We thank Mario Sergio Valdés-Trasanco for support with the gmx_MMPBSA software. This research was funded by the European Union’s Horizon 2020 research and innovation program under Marie Skłodowska Curie Grant 860592.

## Supporting Information Available

Supporting Information provides Supporting Methods, Listings S1–S7, Figures S1–S28 and Table S1.

## TOC Graphic

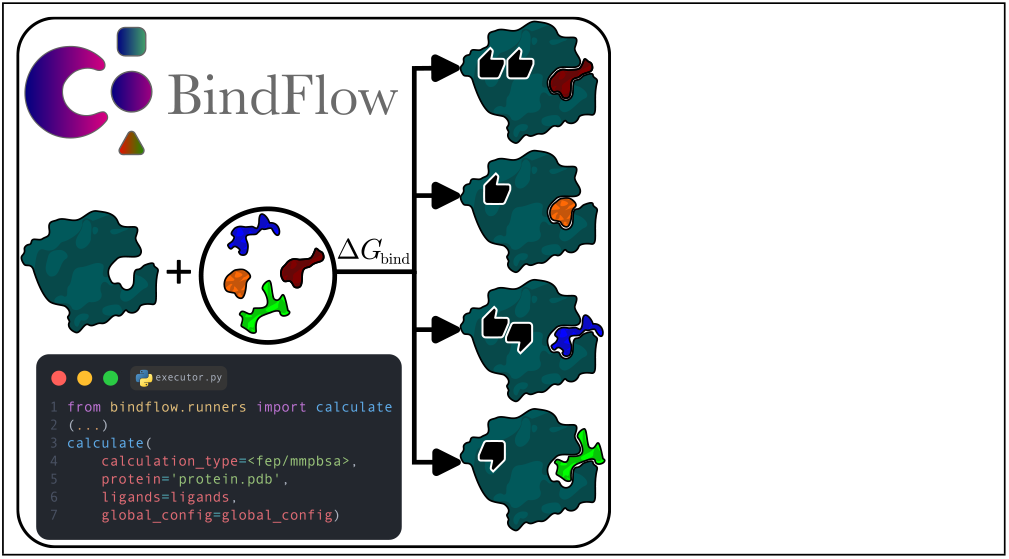

