## Supporting Information for "BindFlow: a free, user-friendly pipeline for absolute binding free energy calculations using free energy perturbation or MM(PB/GB)SA"

### Supporting Methods

#### Statistical measures of agreement with experiment

For each set, the following statistical measures were computed to quantify the agreement of calculated $\Delta G_{\text{calc},i}$ with experimental binding affinities $\Delta G_{\text{exp},i}$: Pearson ($\rho$), Spearman ($r_S$), and Kendall ($\tau$) correlation coefficients as well as root mean-squared error (RMSE), mean signed error (MSE), and mean unsigned error (MUE):

$$\begin{aligned}\text{RMSE} &= \left(\frac{1}{N}\sum_{i=1}^{N}\left(\Delta G_{\text{calc},i} - \Delta G_{\text{exp},i}\right)^2\right)^{1/2} &(1)\\ \text{MSE} &= \frac{1}{N}\sum_{i=1}^{N}(\Delta G_{\text{calc},i} - \Delta G_{\text{exp},i}) &(2)\\ \text{MUE} &= \frac{1}{N}\sum_{i=1}^{N}|\Delta G_{\text{calc},i} - \Delta G_{\text{exp},i}| &(3)\end{aligned}$$

Here, $N$ is the number of ligands probed for the specific set. To account for systematic offsets, for instance due to undersampling of the apo state or a force field bias, we computed the offset-corrected binding affinities on each set by subtracting its corresponded MSE:

$$\Delta G^{\text{oc}}_{\text{calc},i} = \Delta G_{\text{calc},i} - \text{MSE} \tag{4}$$

Offset-corrected RMSE (ocRMSE) and offset-corrected MUE (ocMUE) were calculated by replacing $\Delta G_{\text{calc},i}$ with $\Delta G^{\text{oc}}_{\text{calc},i}$ in Eqs. 1 and 3, respectively.

RMSE values of 1–2 $\text{kcal mol}^{-1}$ have previously been referred to as gold standard for $\Delta G_{\text{bind}}$ calculations, yet typically in the context of RBFE calculations.[1] RBFE calculations are blind to a systematic offset, however, they may be related to ABFE values by fitting to experimental affinities. Thereby, our ocRMSE and ocMUE metrics correspond to RMSE and MUE metrics in RBFE calculations.

The 68% confidence intervals (CIs) for $\rho$, $r_S$, $\tau$, RMSE, ocRMSE, MSE, MUE, and oc-

MUE were estimated using 10,000 rounds of bootstrapping from the corresponding receptor–ligand pairs within each set. In each bootstrap iteration, we resampled $N$ ligands (with replacement) from the original set of $N$ ligands and recalculated all statistical descriptors. The CI was then determined by ranking the 10,000 bootstrap estimates and taking the central 68% (i.e., discarding the lowest 16% and highest 16%). This procedure derived a confidence interval for each descriptor.

The error in the reported $\Delta G_{\mathrm{calc}}^{\mathrm{oc}}$ was determined through uncertainty propagation. The $\mathrm{SEM}_i$ was used as the measurement error of $\Delta G_{\mathrm{calc}}$.

To test whether differences in $\tau$ or ocMUE between methods [FEP versus MM(PB/GB)SA] or force fields (GAFF, OpenFF, or Espaloma) were statistically significant, we estimated a two-sided $p$-value from the bootstrap distribution. For each of 10,000 bootstrap resamples, we calculated the difference $\Delta$ between the two selected methods or force fields. The $p$-value was then defined as $2\min[P(\Delta \leq 0), P(\Delta \geq 0)]$ that is, twice the smaller of the probabilities of observing a non-positive or non-negative difference. This procedure tests whether the observed difference may have been occurred randomly. A difference was considered statistically significant if $p < \alpha = 0.05$.

#### Multi-step equilibration protocols for setting up FEP simulations

Equilibration of non-membrane systems started with a 1 ns NVT phase with a 2 fs integration time step and position restraints on the heavy atoms using a force constant of $2500\,\mathrm{kJ\,mol^{-1}\,nm^{-2}}$. Next, a 1.05 ns NVT phase and approximately 1 ns NPT phase were conducted, both with a 3 fs integration time step and the same position restraints, using the Berendsen barostat. Subsequently, a 5 ns NPT phase with the Parrinello-Rahman barostat and a 4 fs integration time step were performed without restraints.

For the ligand systems, the same protocol was applied, except that (i) the initial 1 ns NVT phase with a 2 fs time step was omitted and (ii) an addition final equilibration under NPT conditions was carried out for 5 ns.

Equilibration of the membrane protein–ligand complex was inspired by the scheme suggested by CHARMM-GUI.[2,3] The process started with minimization using the steepest-descent algorithm and position restraints on the heavy atoms using a force constant of $3000\,\mathrm{kJ\,mol^{-1}\,nm^{-2}}$. This was followed by two steps of 125 ps, each with NVT conditions with a 1 fs integration time step and position restraints of 3000 and $1500\,\mathrm{kJ\,mol^{-1}\,nm^{-2}}$, respectively. Next, a 125 ps NPT phase was conducted and the force constant reduced to $1000\,\mathrm{kJ\,mol^{-1}\,nm^{-2}}$. Subsequently, three steps of 500 ps each with NPT conditions were performed with a 2 fs integration time step, and the restraints were reduced to 500, 200, and $50\,\mathrm{kJ\,mol^{-1}\,nm^{-2}}$, respectively. In contrast to the equilibration simulations of soluble proteins, only bonds involving hydrogen atoms were constrained.

#### Equilibration during FEP simulation

Each $\lambda$ window was energy minimized using the steepest descent algorithm and, subsequently, equilibrated with a three-step protocol: A 10 ps NVT phase was carried out with a 2 fs time step and position restraints on the heavy atoms (force constant $2500\,\mathrm{kJ\,mol^{-1}\,nm^{-2}}$). Next, 100 ps NPT phase was conducted with a 4 fs time step using the same position restraints. Here, the Berendsen barostat was applied. Finally, a 500 ps NPT phase was carried using a 4 fs time step without restraints and using the Parrinello-Rahman barostat.

For the membrane protein–ligand complex, the same protocol was applied except that the cell rescaling barostat was applied.

For future projects, we recommend using the cell rescaling barostat throughout all simulations.

#### Multi-step equilibration protocols for MM(PB/GB)SA simulations

To enable MM(PB/GB)SA calculations at low computational cost, the simulation times for the equilibration protocol were reduced. For soluble complexes, the five steps of the equilibration were carried out for 10, 15, 22.5, and 60 ps, respectively. For the membrane system A2A, the seven steps were carried out for 5, 5, 5, 15, 15, and 45 ps, respectively.

#### Disk usage

BindFlow aims to minimize the disk usage during FEP and MM(PB/GB)SA calculations. In addition, after finishing the simulations, BindFlow provides post-processing archiving and unarchiving functionalities to reduce the required medium-term storage.

As a numerical example, for the P38 system that comprised 86,376 atoms, calculations with 29 ligands (triplicated calculations) required 320 GB disk space for FEP and 40 GB for MMGBSA during runtime, respectively.

By excluding log files (*.snakemake* directory and **.log* and **.err* files) and irrelevant GROMACS files (**.edr*, *mdout.mdp* and **.tpr*) during archiving and compressing all non-trajectory files, the disk space was reduced to 137 GB for FEP and 19 GB for MMGBSA (see Fig. S1). However, owing to BindFlow full automation, to reproduce the simulations, only the BindFlow version, input structures, run script, and configuration file are required; involving typically only few megabytes for long-term archive.

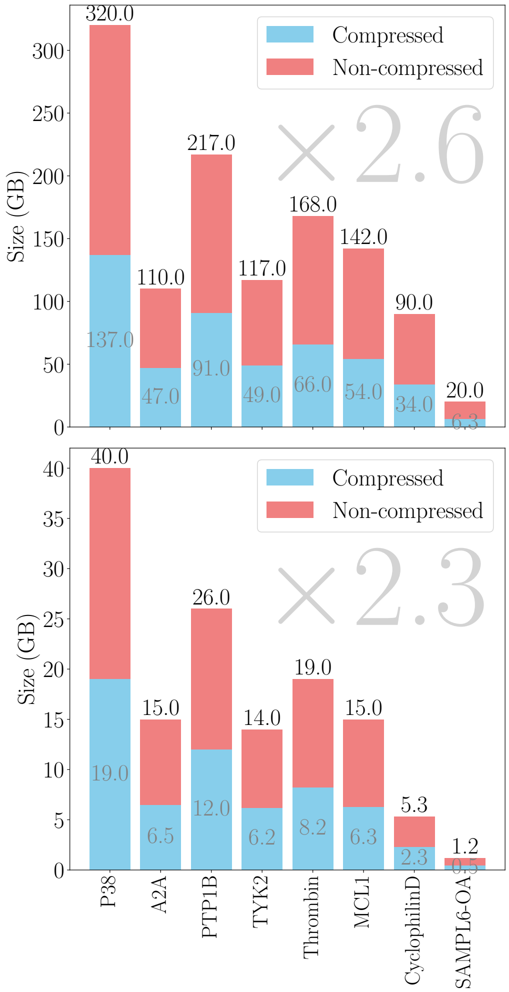


Figure S1: Disk space used by BindFlow for all simulation sets of this study. Top: for FEP. Bottom: for MM(PB/GB)SA. Red bars: disk space used during simulations. Blue bars: after compression of raw simulation data, yielding compression factors of 2.6 and 2.3 for FEP and MM(PB/GB)SA, respectively.

```python
import yaml
from bindflow.orchestration.generate_scheduler import FrontEnd
from bindflow.runners import calculate

ligands = [
    'path/to/ligand1.mol',
    'path/to/ligand2.mol'
    'path/to/ligand3.mol'
]

with open('path/to/config.yml', 'r') as c:
    global_config = yaml.safe_load(c)

calculate(
    calculation_type='mmpbsa',
    protein='path/to/protein.pdb',
    ligands=ligands,
    membrane='path/to/membrane.pdb',
    cofactor='path/to/cofactor.mol',
    cofactor_on_protein=True,
    water_model='amber/tip3p',
    hmr_factor=3,
    dt_max=0.004,
    threads=4,
    num_jobs=12,
    replicas=1,
    scheduler_class=FrontEnd,
    out_root_folder_path='mmpbsa',
    submit=True,
    global_config=global_config)
```

```yaml
extra_directives:
  dependencies:
    - module load gromacs/2022.4
    - export GMX_MAXBACKUP=-1
  mdrun:
    all:
      cpi: False
      stepout: 5000
      v: True
      ntmpi: 1
samples: 20
mdp:
  complex:
    equi:
      00_min:
        nsteps: 100000
      01_nvt:
          dt: 0.001
          nsteps: 5000
      02_nvt:
          dt: 0.001
          nsteps: 5000
      03_npt:
        dt: 0.001
        nsteps: 5000
      04_npt:
        dt: 0.002
        nsteps: 7500
      05_npt:
        dt: 0.002
        nsteps: 7500
      06_npt:
        dt: 0.003
        nsteps: 15000
      prod:
        dt: 0.004
        nsteps: 237500
        nstxout-compressed: 11875
    mmpbsa:
      prod:
        dt: 0.004
        nsteps: 25000
        nstxout-compressed: 1250
mmpbsa:
  general:
    c2_entropy: 1
    interaction_entropy: 1
  pb: {}
  gb: {}
```

Listing S1: MM(PB/GB)SA calculation of a membrane system executed in a desktop computer. **Left**: Calling of the *bindflow.runners.calculate* function with *threads* and *num_jobs* adjusted to the available frontend resources **Right**: Example configuration YAML file with customized parameters.

```
component = {
    "conf": "<valid configuration file (coordinates of the system)>",
    "top": "<GROMACS topology file>",
    "ff": {
        "code": "<force field code>",
        "type": "<force field type, only needed for ligand and cofactor>"
    },
    "is_water": "<bool, only needed for cofactor>",
    "custom_ff_path": "<the path to the '*.ff' directory of your custom force field>"
}
```

Listing S2: Structure of the optional dictionary used to define components such as ligands, proteins, membranes, or cofactors. *component* is passed to the main BindFlow function *bindflow.runners.calculate*. Consult the full online documentation of *bindflow.runners.calculate* for more details.

```
component = {
    "conf": "protein.gro",
    "top": "protein.top",
    "ff": {
        "code": "charmm36-jul2022",
    },
    "custom_ff_path": "/home/users/john-doe/FFs"
}
```

Listing S3: Example for including of a custom force field. The topology file may contain statements such as: `#include` “`/home/users/john-doe/FFs/charmm36-jul2022.ff/forcefield.itp`”.

```
component = {
    "conf": "water-cofactor.gro",
    "top": "water-cofactor.top",
    "is_water": True
}
```

Listing S4: Example for how to define specific water molecules as cofactors.

```
component = {
    "conf": "protein.pdb",
    "top": "protein.top",
    "ff": {
        "code": "AMBER94",
    },
}
```

Listing S5: Example for using a specific force fields, here Amber94, from the GROMACS distribution.

```
component = {
    "conf": "ligand.mol",
    "ff": {
        "code": "espaloma-0.3.2",
        "type": "espaloma"
    },
}
```

Listing S6: Example for how to generate Espaloma-0.3.2 force parameters for a small molecule.

```
cluster:
  options: # Depending on the Scheduler
    calculation:
      <cluster_options_for_calculation_jobs>
    job:
      <cluster_options_for_launcher_job>
extra_directives:
  dependencies:
    - <dependency_commands>
  mdrun:
    ligand:
      <mdrun_keywords_for_ligand_simulation>
    complex:
      <mdrun_keywords_for_complex_simulation>
    all:
      <mdrun_keywords_for_ligand_and_complex_simulation>
nwindows:
  ligand:
      vdw: <number_of_vdw_windows>
      coul: <number_of_coul_windows>
  complex:
      vdw: <number_of_vdw_windows>
      coul: <number_of_coul_windows>
      bonded: <<number_of_bonded_windows>
samples: <number_of_samples_for_mmpbsa>
mmpbsa:
  <mm(pb/gb)sa_options>
mdp:
  ligand:
    equi:
      <step>:
        <mdp_ligand_equi_step_options>
    fep:
      vdw:
        <step>:
          <mdp_ligand_fep_vdw_step_options>
      coul:
        <step>:
          <mdp_ligand_fep_coul_step_options>
  complex:
    equi:
      <step>:
        <mdp_complex_equi_step_options>
    fep:
      vdw:
        <step>:
          <mdp_complex_fep_vdw_step_options>
      coul:
        <step>:
          <mdp_complex_fep_coul_step_options>
      bonded:
        <step>:
          <mdp_complex_fep_bonded_step_options>
    mmpbsa:
      prod:
        <mdp_complex_mm(pb/gb)sa_options>
```

Listing S7: List of options that may be specified by the user, as passed to *global_config*.

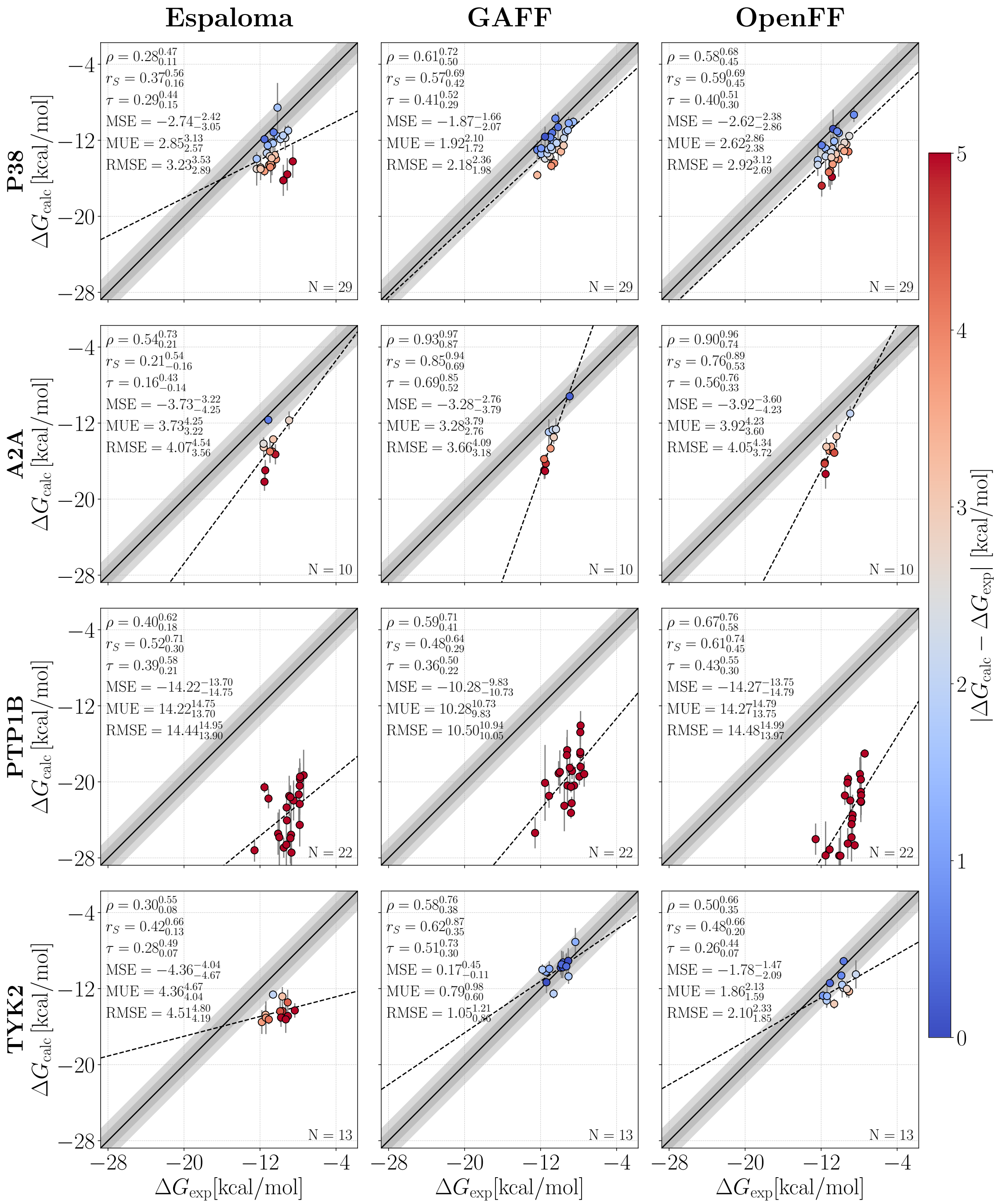


Figure S2: Correlation between $\Delta G_{\text{calc}}$ and $\Delta G_{\text{exp}}$ from FEP. Correlations are shown for (from left to right) Espaloma-0.3.1, GAFF-2.11, and OpenFF-2.0.0 and for (from top to bottom) P38, A2A, PTP1B, and TYK2. Colors of dots indicate the absolute deviation between $\Delta G_{\text{calc}}$ and $\Delta G_{\text{exp}}$ (see color bar). Dark and light gray diagonal regions indicate 1 and 2 $\text{kcal mol}^{-1}$ deviations, respectively. Dashed lines are linear fits shown to guide the eye. Insets report number of ligands $N$, Pearson $\rho$, Spearman $r_S$, Kendall $\tau$, MSE, MUE, and RMSE (last three in $\text{kcal mol}^{-1}$) for each data set with its corresponding 68 % confident interval. Error bars show uncertainties obtained via three independent replicates. The MSE was not removed from the data.

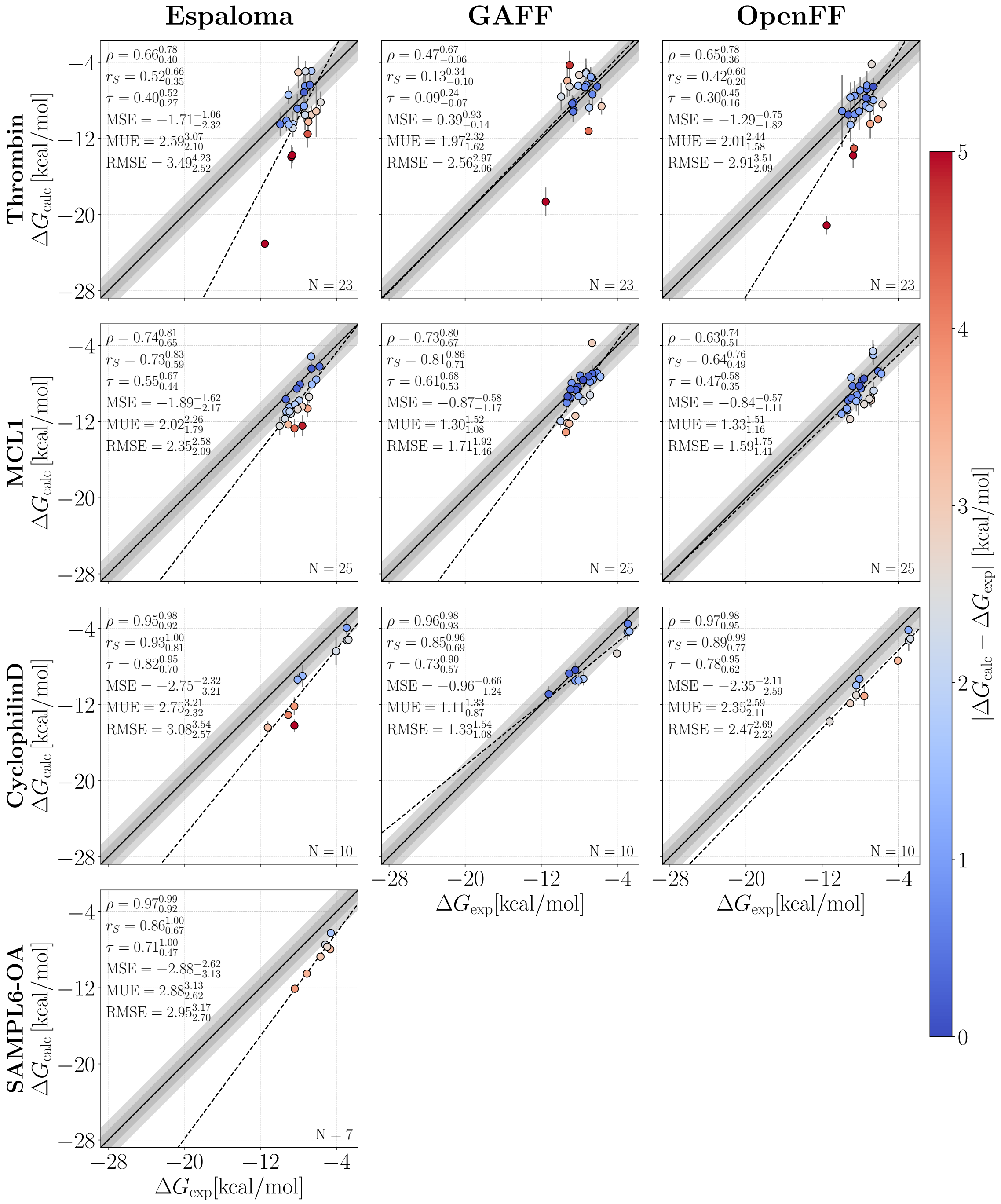


Figure S3: Correlation between $\Delta G_{\mathrm{calc}}$ and $\Delta G_{\mathrm{exp}}$ from FEP for (from top to bottom) Thrombin, MCL1, Cyclophilin D, and SAMPL6-OA. Presentation according to Fig. S2.

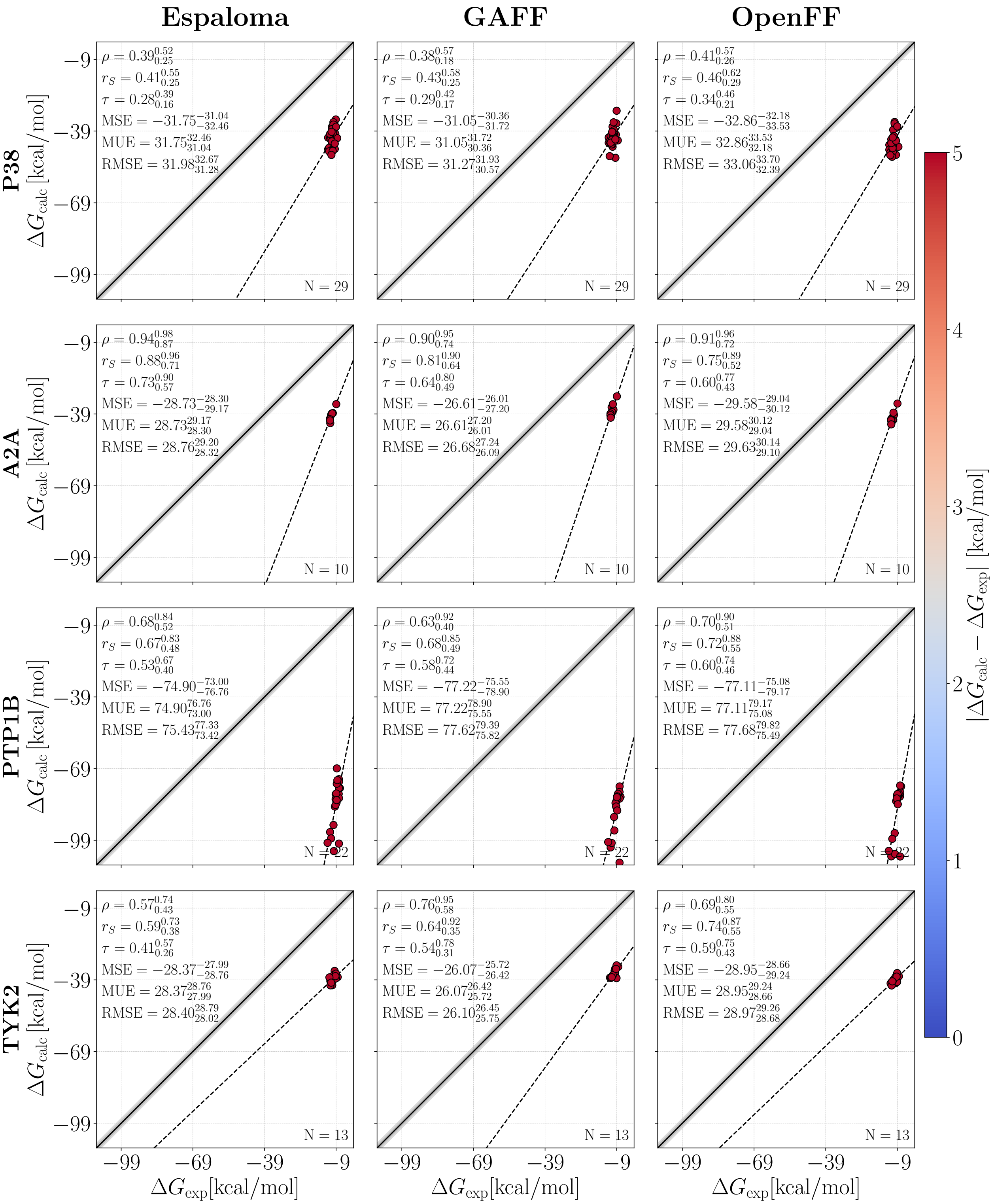


Figure S4: Correlation between $\Delta G_{\mathrm{calc}}$ and $\Delta G_{\mathrm{exp}}$ from MMGBSA without entropy correction for (from top to bottom) P38, A2A, PTP1B, and TYK2. Presentation according to Fig. S2.

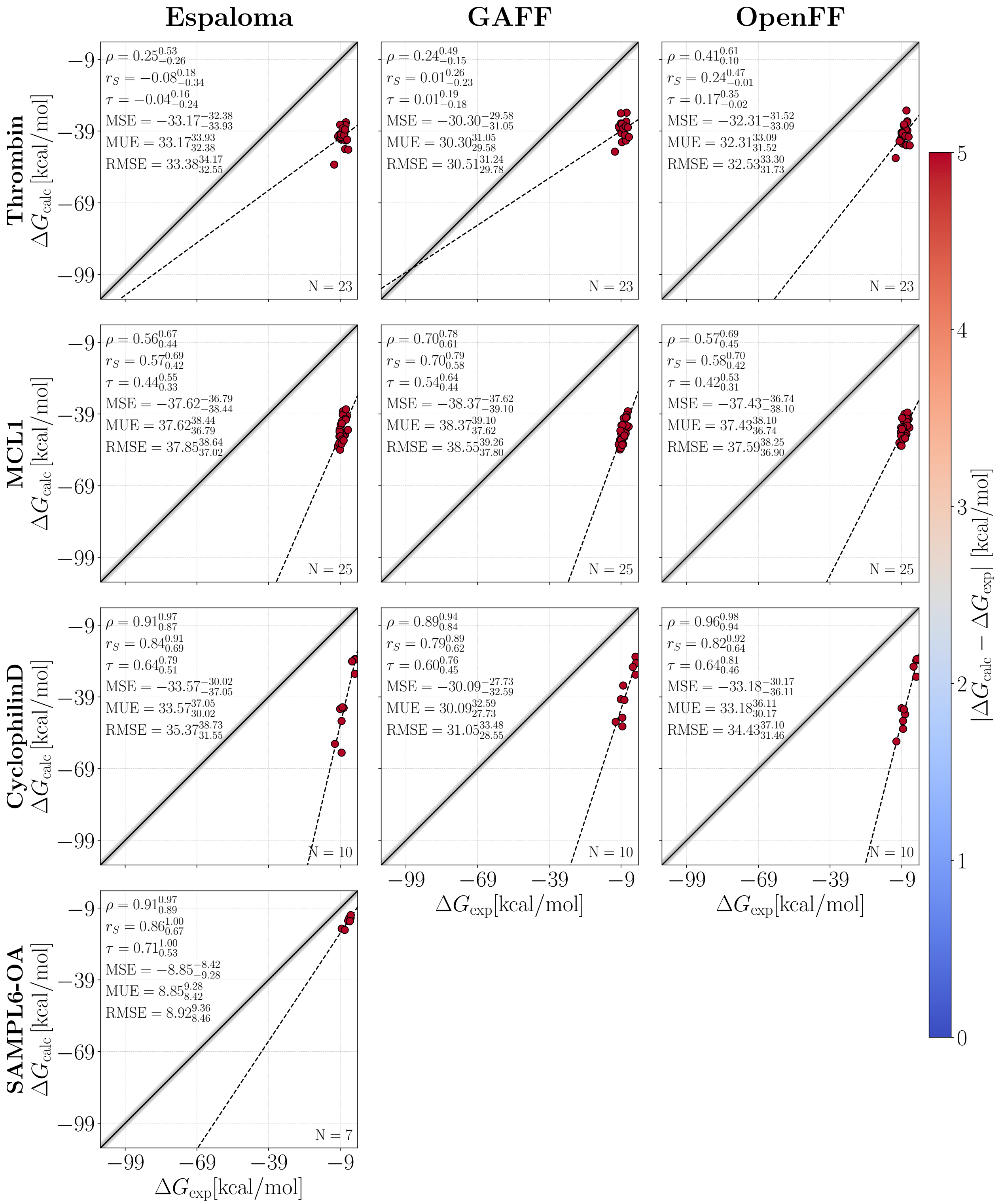


Figure S5: Correlation between $\Delta G_{\mathrm{calc}}$ and $\Delta G_{\mathrm{exp}}$ from MMGBSA without entropy correction for (from top to bottom) Thrombin, MCL1, Cyclophilin D, and SAMPL6-OA. Presentation according to Fig. S2.

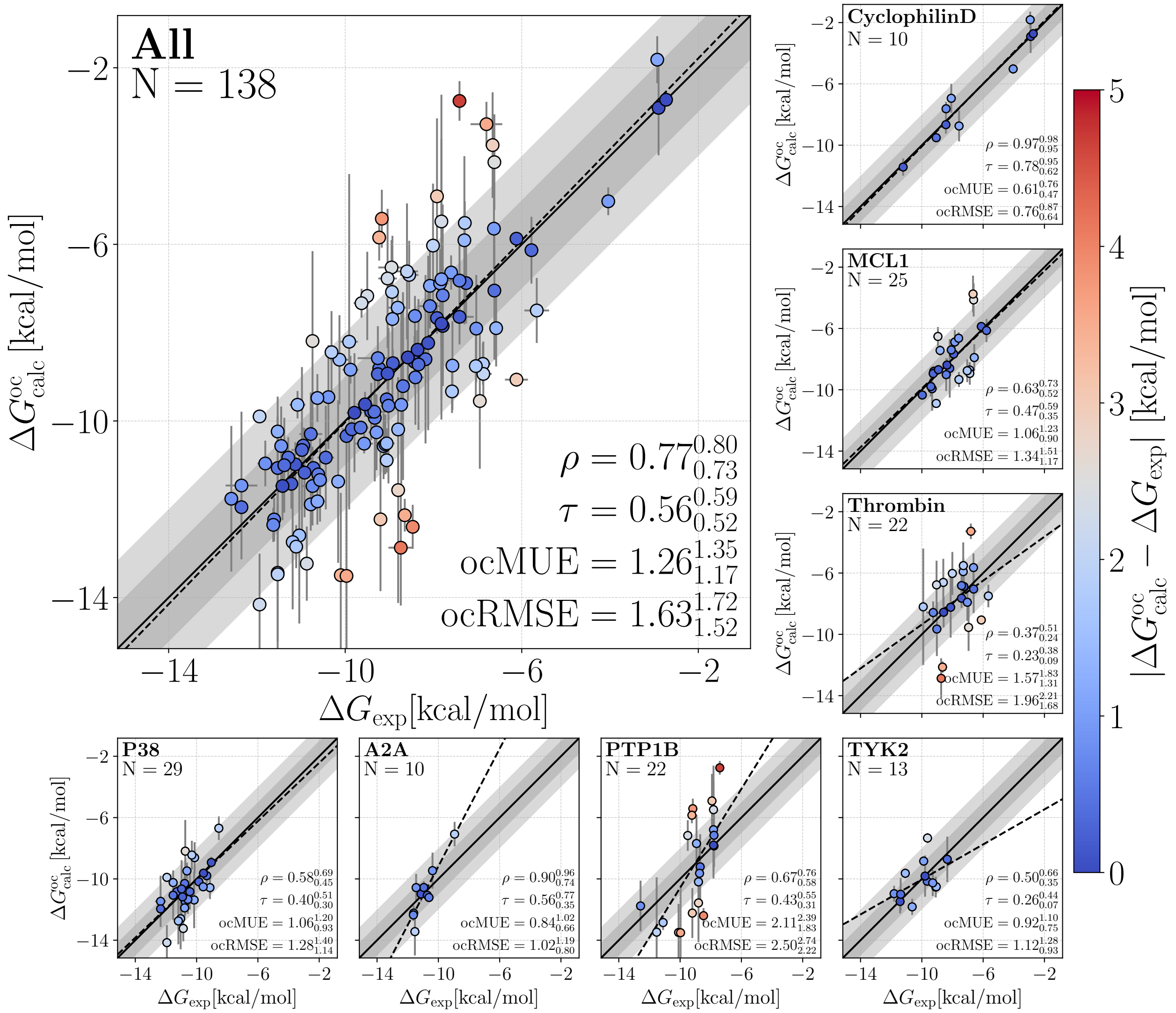


Figure S6: Offset-corrected calculated affinities $\Delta G_{\mathrm{calc}}^{\mathrm{oc}}$ versus experimental affinities $\Delta G_{\mathrm{exp}}$ from FEP with OpenFF-2.0.0. Presentation style according to Fig. 4.

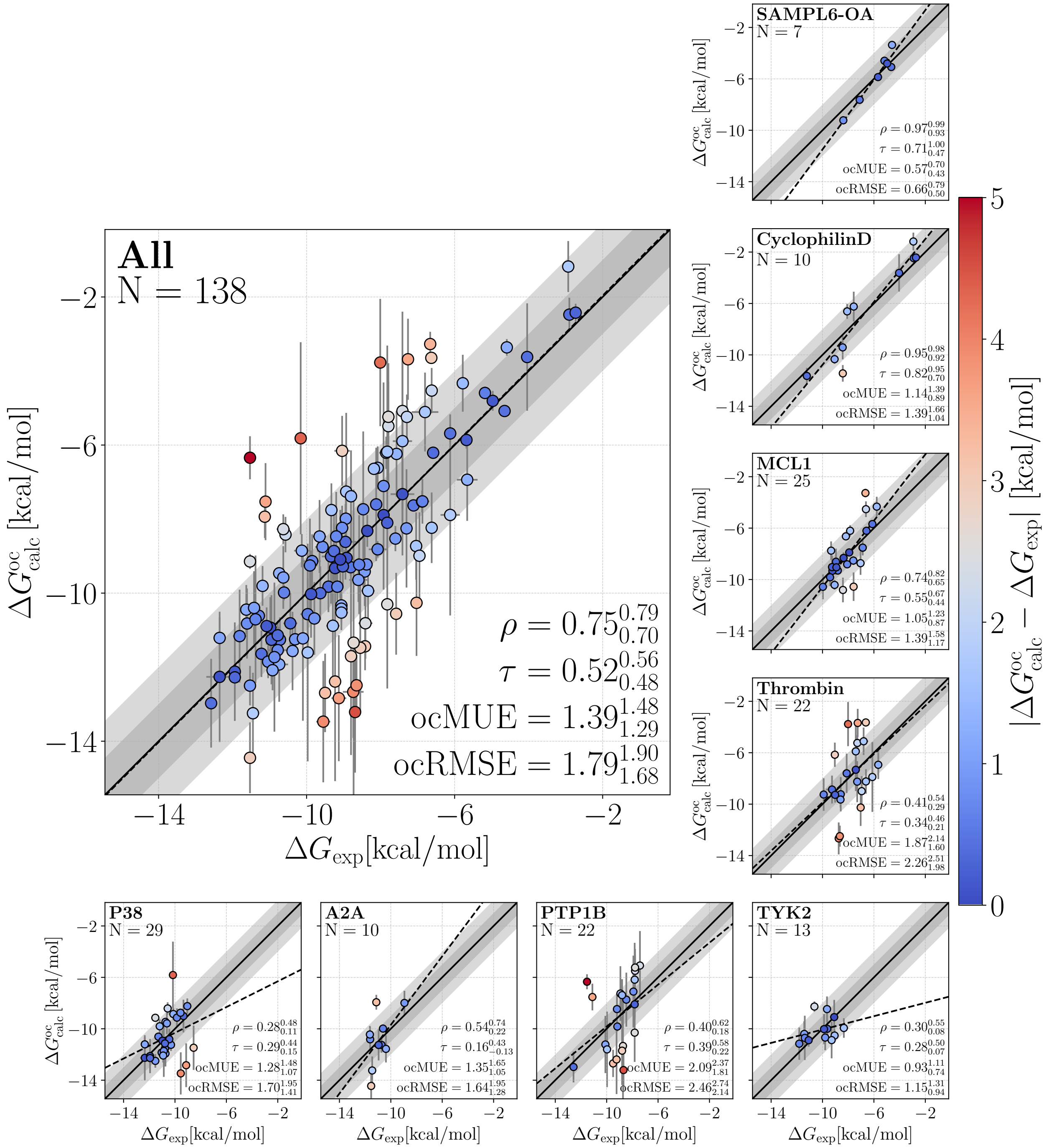


Figure S7: Offset-corrected calculated affinities $\Delta G^{oc}_{calc}$ versus experimental affinities $\Delta G_{exp}$ from FEP with Espaloma-0.3.1. Presentation style according to Fig. 4.

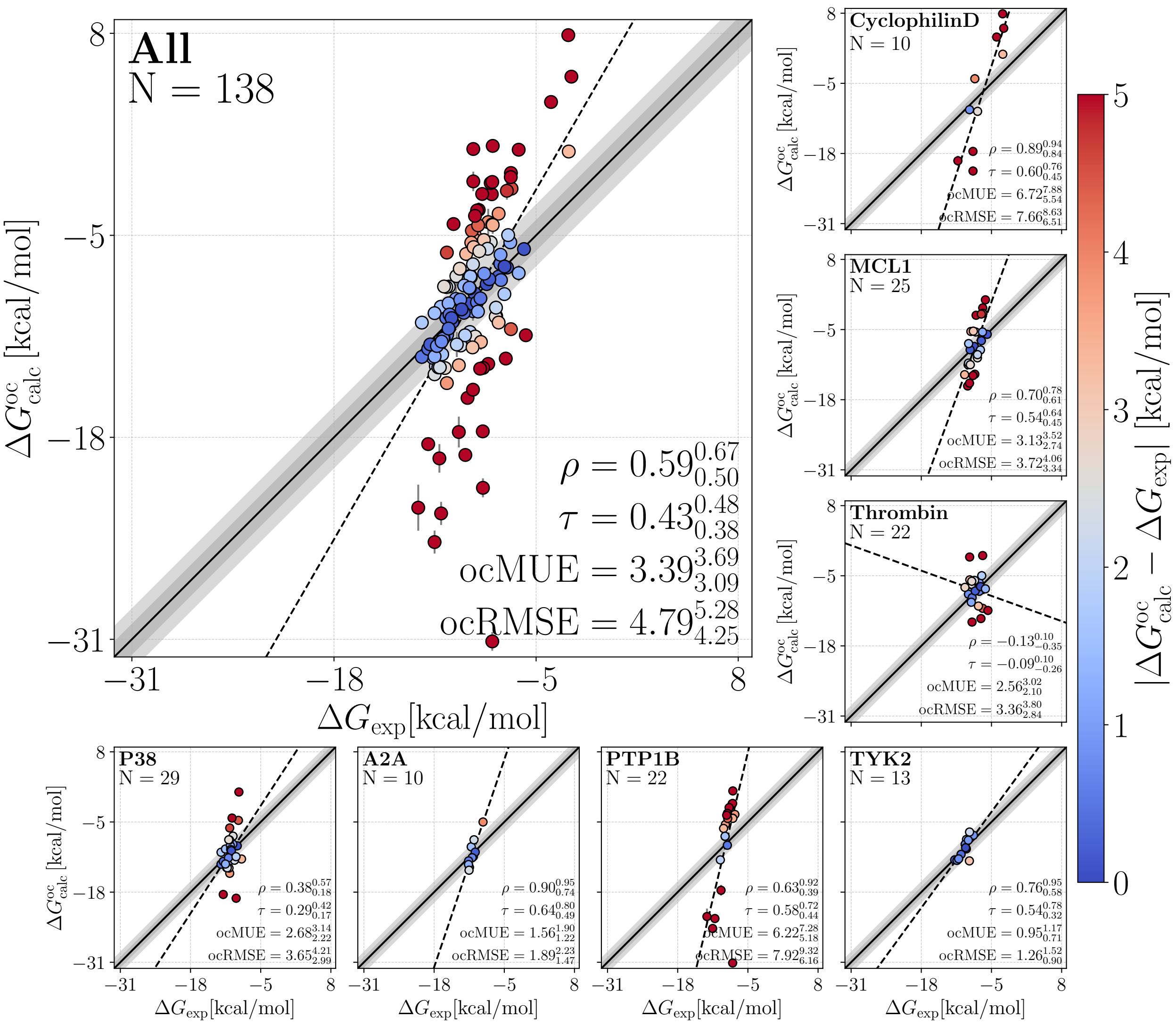


Figure S8: Offset-corrected calculated affinities $\Delta G^{\mathrm{oc}}_{\mathrm{calc}}$ versus experimental affinities $\Delta G_{\mathrm{exp}}$ from MMGBSA without entropy correction and using GAFF-2.11. Presentation style according to the main text Fig. 4.

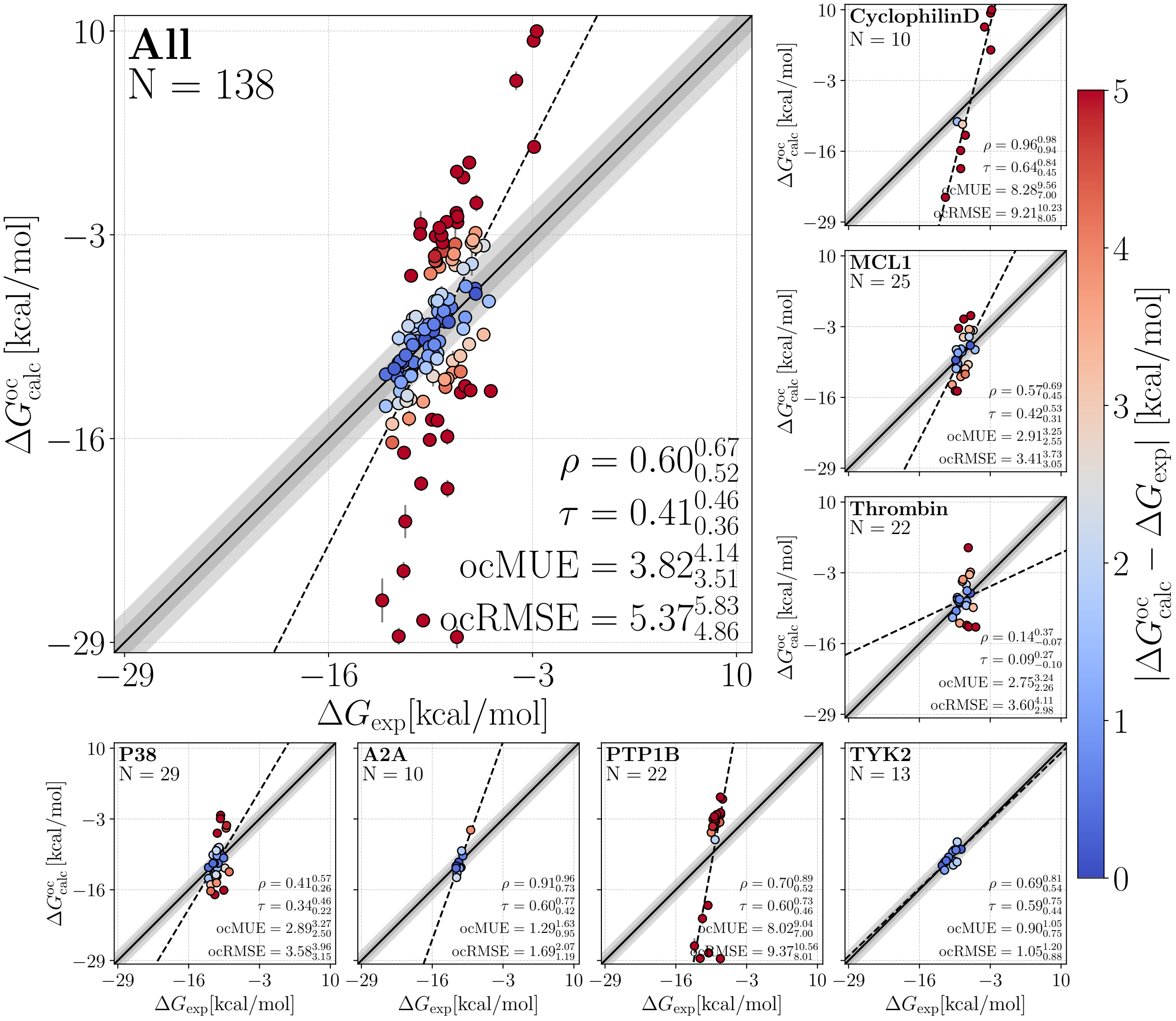


Figure S9: Offset-corrected calculated affinities $\Delta G_{\mathrm{calc}}^{\mathrm{oc}}$ versus experimental affinities $\Delta G_{\mathrm{exp}}$ from MMGBSA without entropy correction and using OpenFF-2.0.0. Presentation style according to the main text Fig. 4.

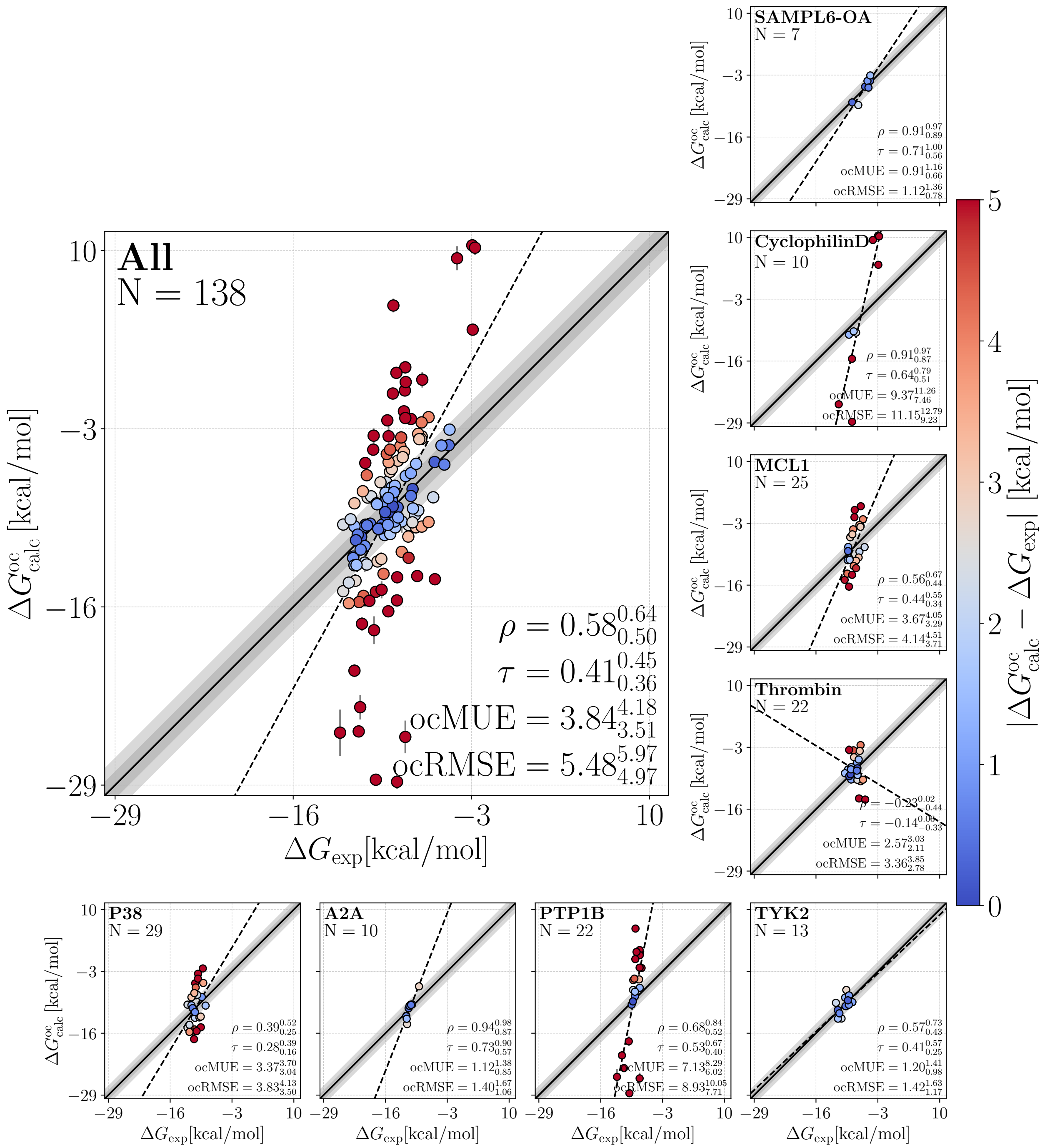


Figure S10: Offset-corrected calculated affinities $\Delta G_{\mathrm{calc}}^{\mathrm{oc}}$ versus experimental affinities $\Delta G_{\mathrm{exp}}$ from MMGBSA without entropy correction and using Espaloma-0.3.1. Presentation style according to the main text Fig. 4.

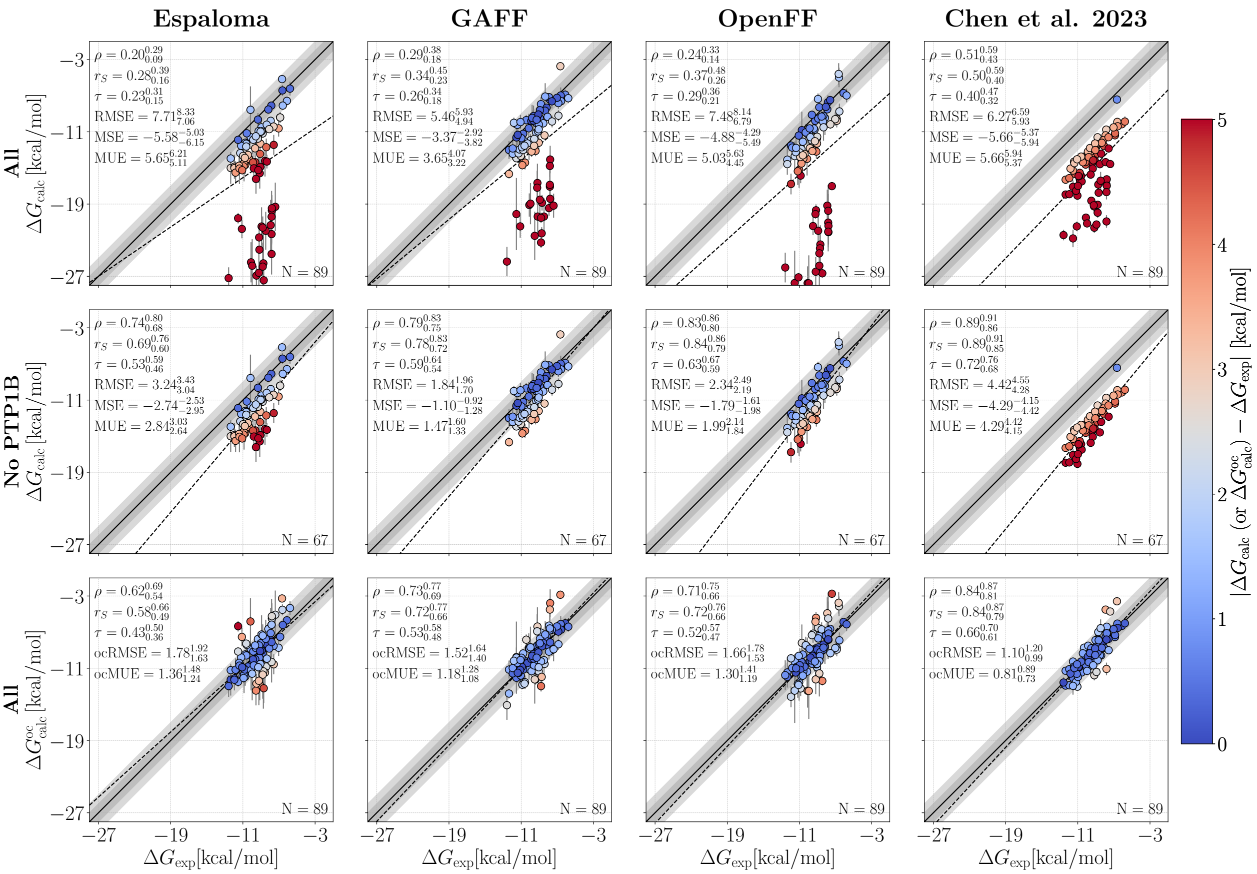


Figure S11: Comparison with Chen et al.[4]. Correlation between $\Delta G_{\mathrm{calc}}$ (or $\Delta G^{\mathrm{oc}}_{\mathrm{calc}}$) and $\Delta G_{\mathrm{exp}}$ from FEP. Correlations are shown for (from left to right) Espaloma-0.3.1, GAFF-2.11, OpenFF-2.0.0, and the study of Chen et al.[4] To enable quantitative comparison, metrics were computed from the same set of ligands of P38, PTP1B, TYK2 and MCL1 sets. The top row shows all sets, the middle row removes ligands from the PTP1B set, and the bottom row includes all sets with the set-specific MSE subtracted from each set. Colors of dots indicate the absolute deviation between $\Delta G_{\mathrm{calc}}$ (or $\Delta G^{\mathrm{oc}}_{\mathrm{calc}}$) and $\Delta G_{\mathrm{exp}}$ (see color bar). Dark and light gray diagonal regions indicate 1 and 2 $\mathrm{kcal\,mol^{-1}}$ deviations, respectively. Dashed lines are linear fits shown to guide the eye. Insets report number of ligands $N$, Pearson $\rho$, Spearman $r_S$, Kendall $\tau$, RMSE (or ocRMSE), MSE (or ocMSE), and MUE for each data set with its corresponding 68 % confident interval.

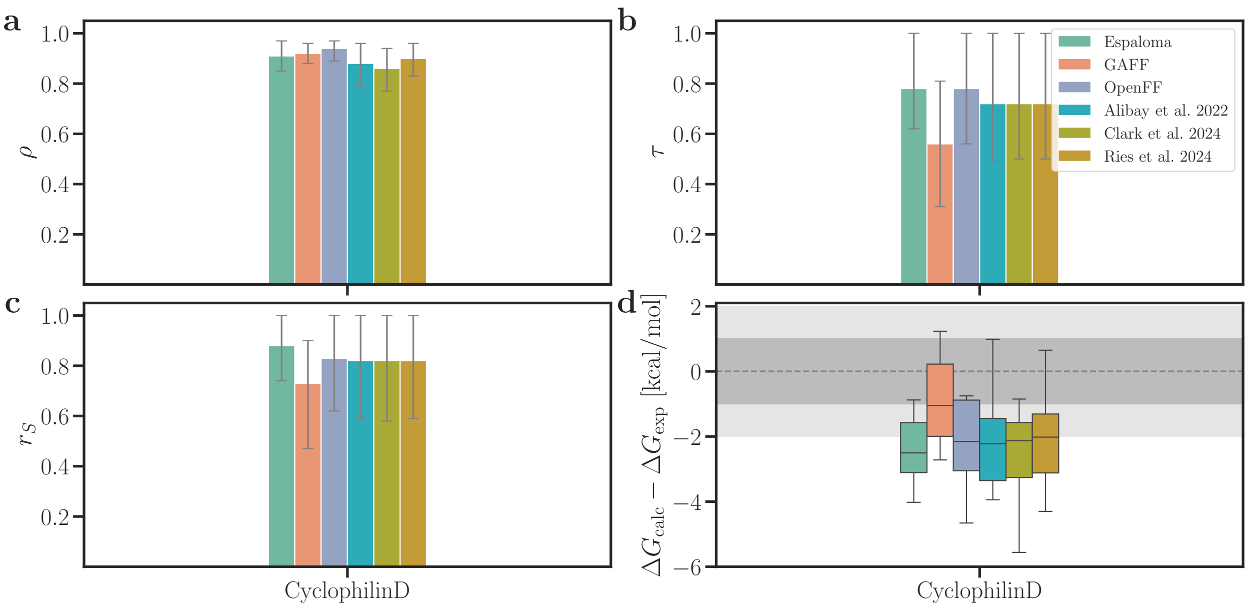


Figure S12: Comparison of statistical metrics for cyclophilin D from BindFlow using Espaloma, GAFF, or OpenFF with previous studies: Alibay et al.[5], Clark et al.[6], and Ries et al.[7] using GAFF-2 force field variants (see legend for color code). To enable quantitative comparison, metrics were computed from the same set of ligands and using three FEP replicates. Presentation style according to Fig. 5.

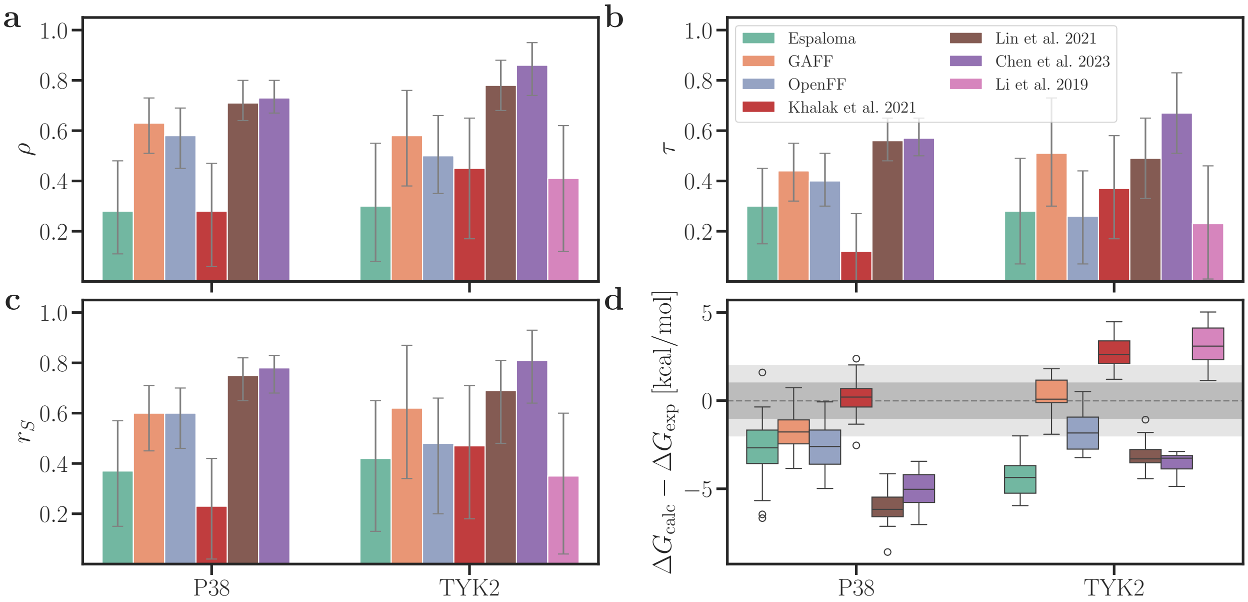


Figure S13: Comparison of statistical metrics for P38 and TYK2 from BindFlow using Espaloma, GAFF, or OpenFF with previous studies: Khalak et al.[8], Lin et al.[9], Chen et al.[4], Li et al.[10] (see legend for color code). In contrast to BindFlow, Khalak et al.[8] used non-equilibrium simulations together with Crook's Theorem for obtaining ABFE values. To enable quantitative comparison, metrics were computed from the same set of ligands. Presentation style according to Fig. 5.

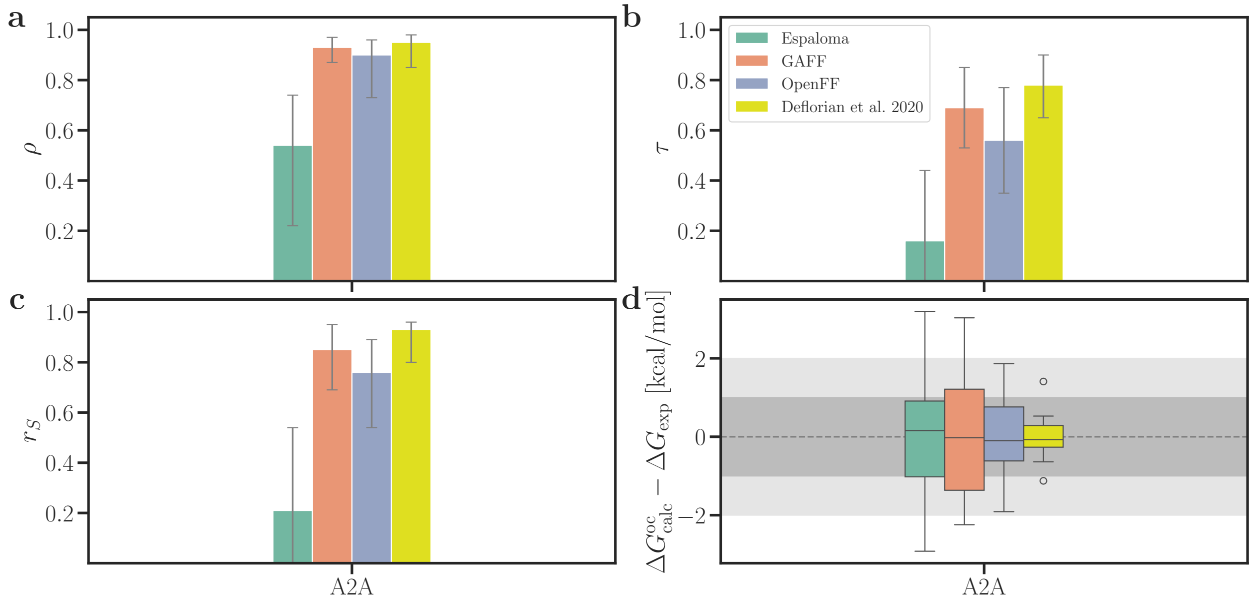


Figure S14: Comparison of statistical metrics for A2A from BindFlow using Espaloma, GAFF, or OpenFF with RBFE calculations by Deflorian et al.[11]. To enable quantitative comparison, metrics were computed from the same set of ligands. Presentation style according to Fig. 5.

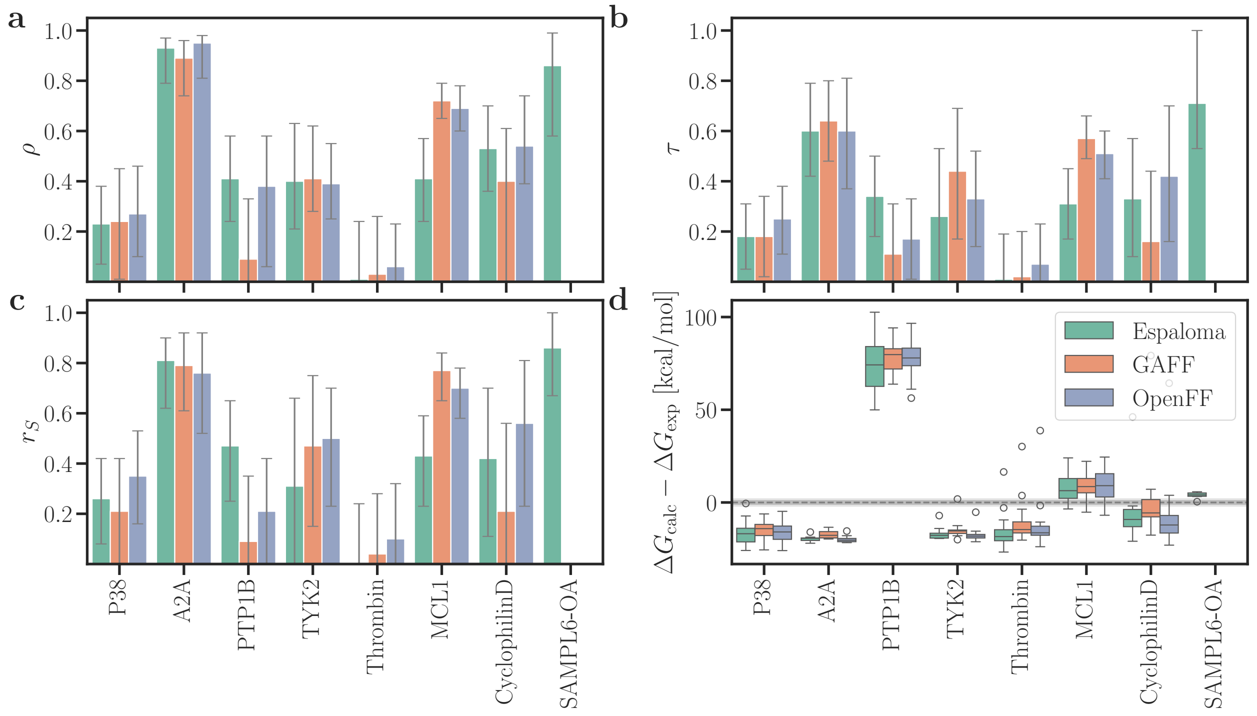


Figure S15: Statistical metrics from MMGBSA with C2 entropy correction. Presentation style according to Fig. 5.

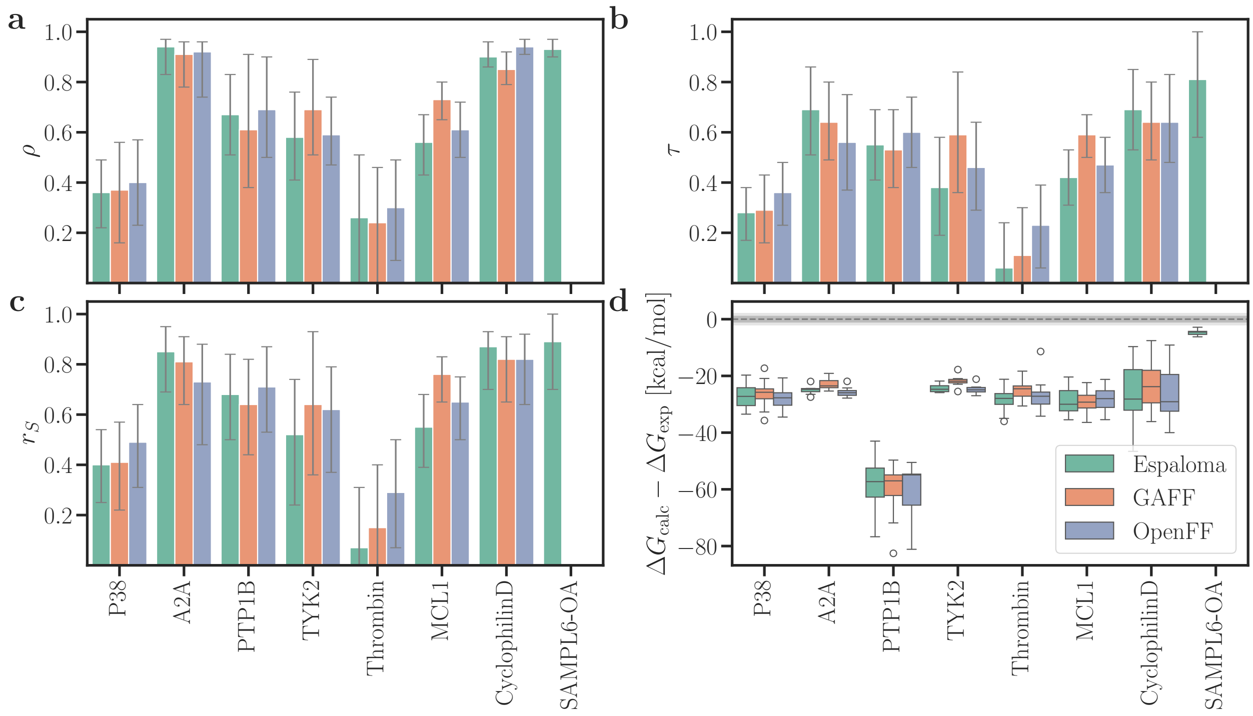


Figure S16: Statistical metrics from MMGBSA with IE entropy correction. Presentation style according to Fig. 5.

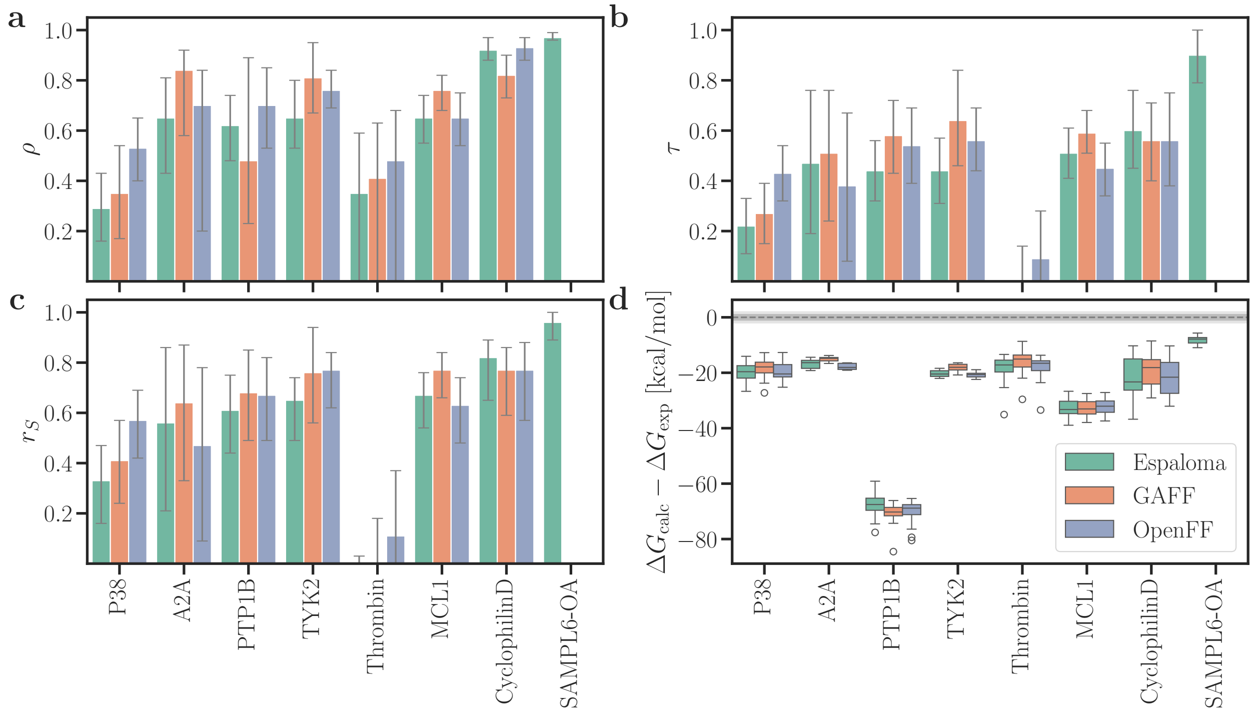


Figure S17: Statistical metrics from MMPBSA without entropy correction. Presentation style according to Fig. 5.

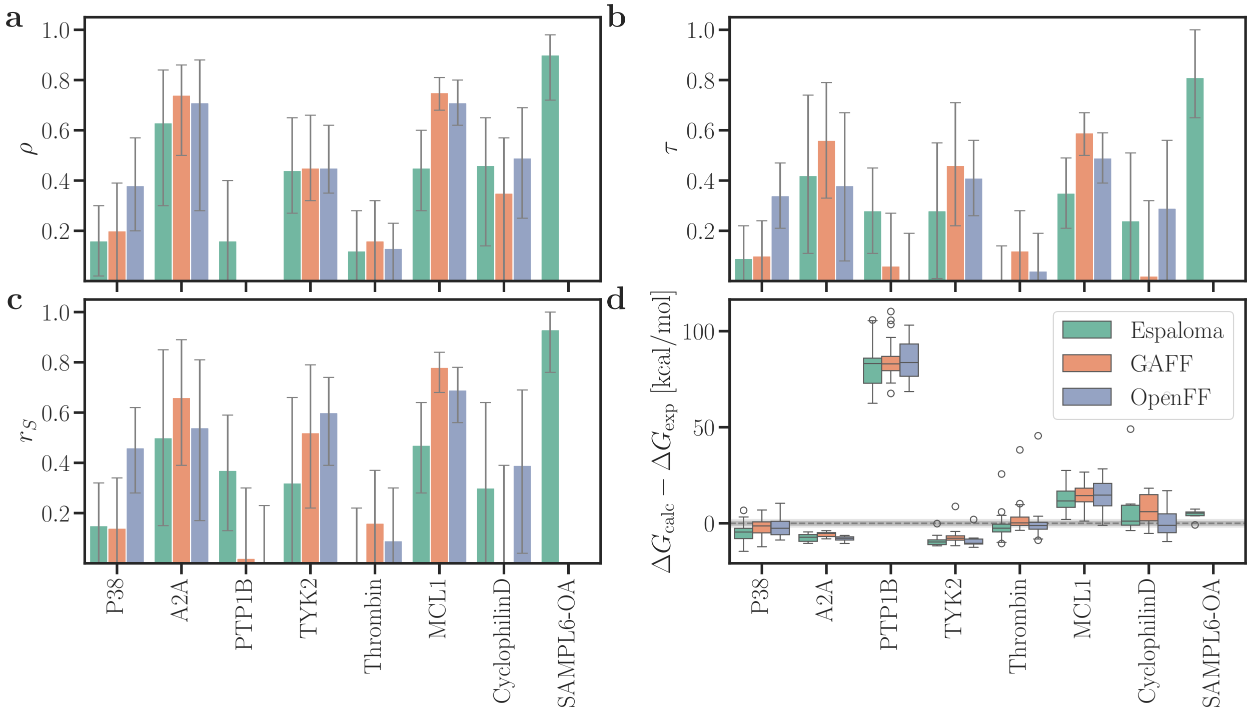


Figure S18: Statistical metrics from MMPBSA with C2 entropy correction. Presentation style according to Fig. 5.

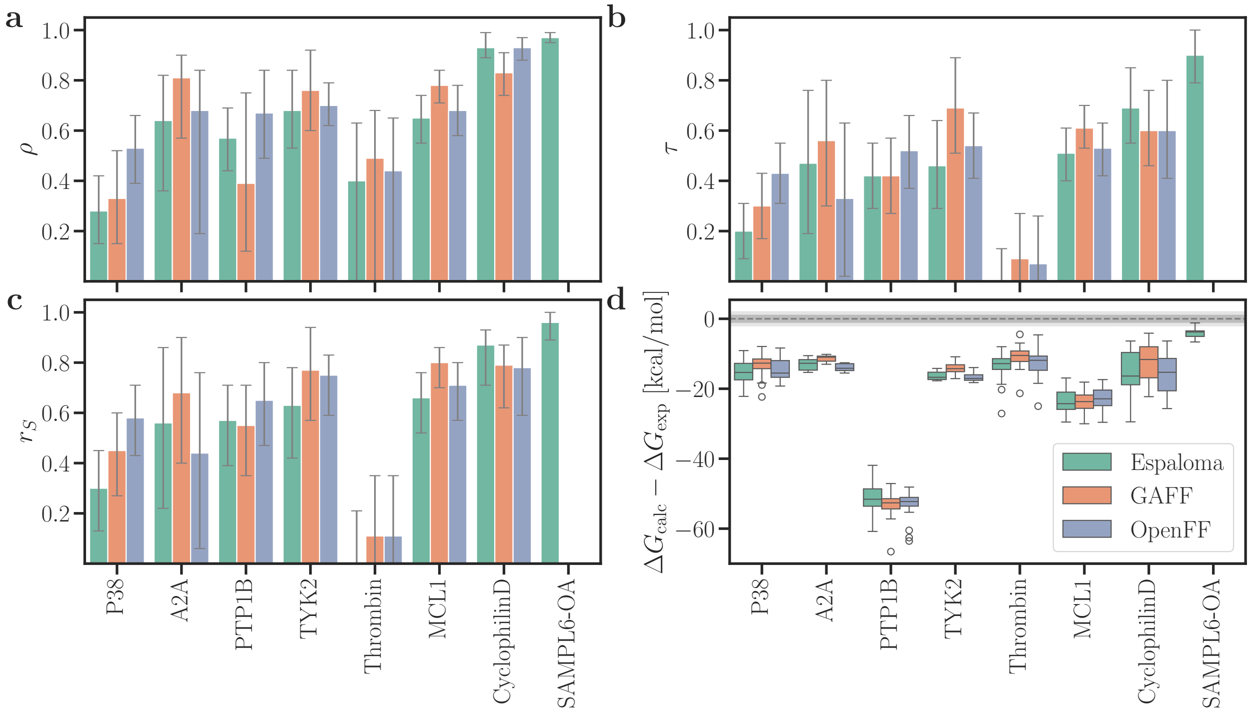


Figure S19: Statistical metrics from MMPBSA with IE entropy correction. Presentation style according to Fig. 5.

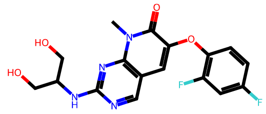

lig_p38a_2aa
ic50 = 1.58e+02 nM

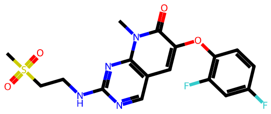

lig_p38a_2bb
ic50 = 2.23e+02 nM

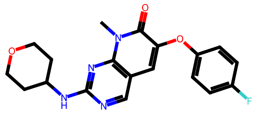

lig_p38a_2e
ic50 = 1.40e+01 nM

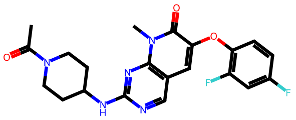

lig_p38a_2ee
ic50 = 1.00e+00 nM

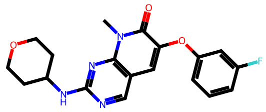

lig_p38a_2f
ic50 = 5.94e+02 nM

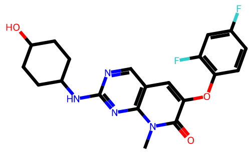

lig_p38a_2ff
ic50 = 4.00e+00 nM

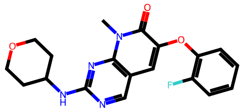

lig_p38a_2g
ic50 = 1.50e+01 nM

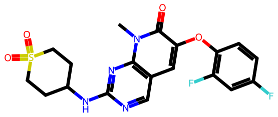

lig_p38a_2gg
ic50 = 1.50e+01 nM

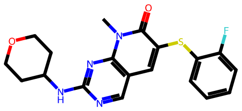

lig_p38a_2h
ic50 = 1.11e+02 nM

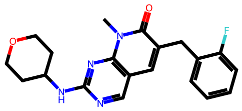

lig_p38a_2i
ic50 = 4.20e+01 nM

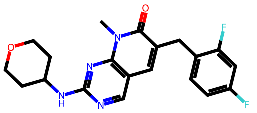

lig_p38a_2j
ic50 = 4.00e+01 nM

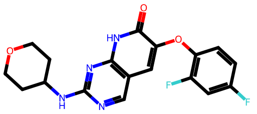

lig_p38a_2k
ic50 = 1.80e+01 nM

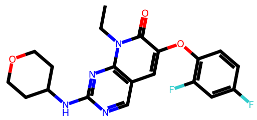

lig_p38a_2l
ic50 = 1.00e+00 nM

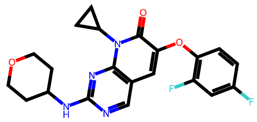

lig_p38a_2m
ic50 = 9.00e+00 nM

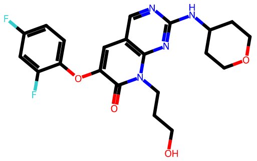

lig_p38a_2n
ic50 = 2.00e+00 nM

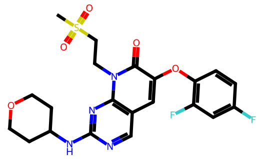

lig_p38a_2o
ic50 = 2.00e+01 nM

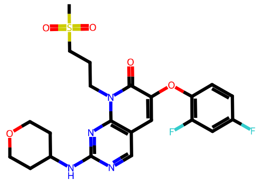

lig_p38a_2p
ic50 = 6.00e+00 nM

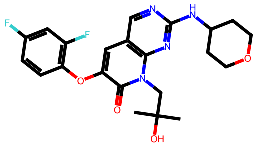

lig_p38a_2r
ic50 = 1.20e+01 nM

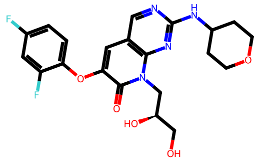

lig_p38a_2s
ic50 = 7.00e+00 nM

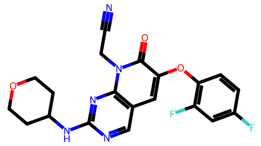

lig_p38a_2t
ic50 = 2.00e+00 nM

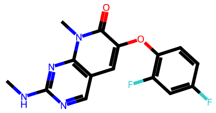

lig_p38a_2v
ic50 = 2.60e+02 nM

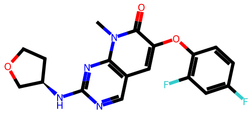

lig_p38a_2x
ic50 = 3.10e+01 nM

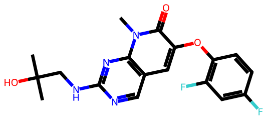

lig_p38a_2y
ic50 = 6.70e+01 nM

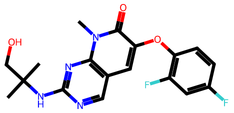

lig_p38a_2z
ic50 = 2.50e+01 nM

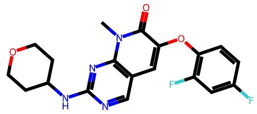

lig_p38a_3fln
ic50 = 1.00e+01 nM

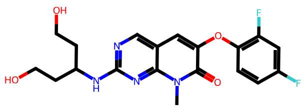

lig_p38a_3flw
ic50 = 1.40e+01 nM

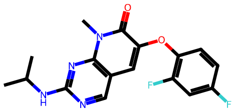

lig_p38a_3fly
ic50 = 4.00e+00 nM

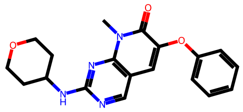

lig_p38a_3flz
ic50 = 1.06e+02 nM

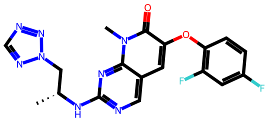

lig_p38a_3fmh
ic50 = 1.10e+01 nM

Figure S20: Ligands of the P38 set.

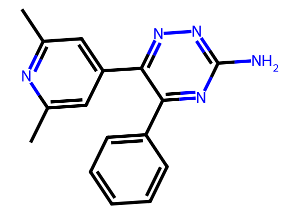
4g
kd = 7.76e-09 M

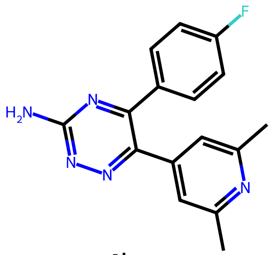
4h
kd = 1.55e-08 M

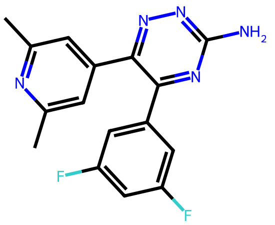
4i
kd = 2.75e-08 M

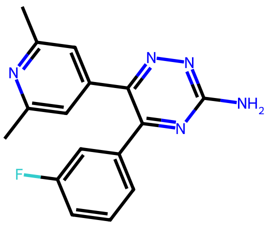
4j
kd = 1.05e-08 M

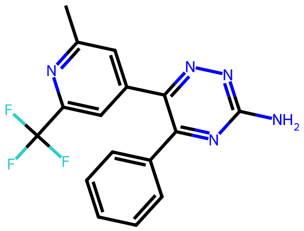
4k
kd = 3.47e-09 M

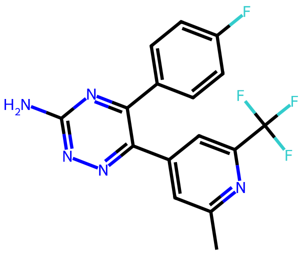
4l
kd = 4.57e-09 M

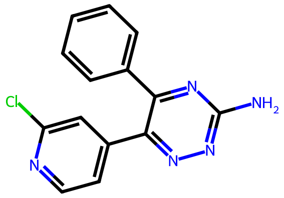
4m
kd = 1.86e-08 M

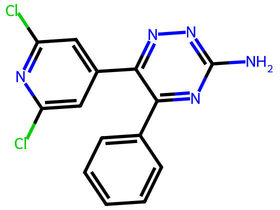
4n
kd = 3.39e-09 M

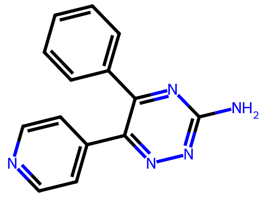
4q
kd = 3.09e-07 M

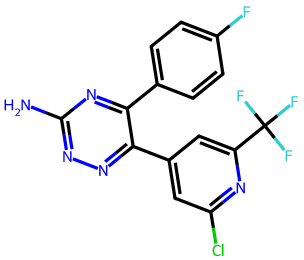
4r
kd = 3.98e-09 M

Figure S21: Ligands of the A2A set.

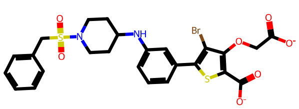
lig_20667_2qbp
kd = 4.00e-03 uM

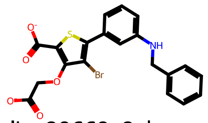
lig_20669_2qbr
kd = 4.70e-01 uM

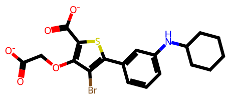
lig_20670_2qbs
kd = 2.10e-01 uM

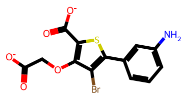
lig_23466
kd = 1.70e+00 uM

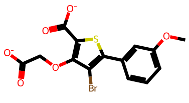
lig_23467
kd = 4.00e+00 uM

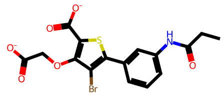
lig_23468
kd = 2.00e+00 uM

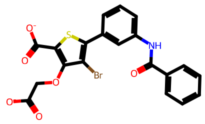
lig_23469
kd = 2.10e+00 uM

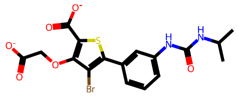
lig_23470
kd = 2.10e+00 uM

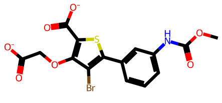
lig_23471
kd = 2.00e+00 uM

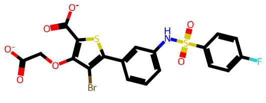
lig_23472
kd = 4.40e-01 uM

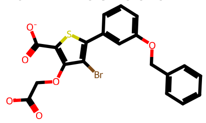
lig_23473
kd = 6.80e-01 uM

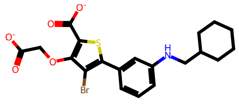
lig_23474
kd = 2.00e-01 uM

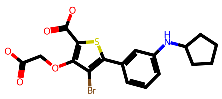
lig_23475
kd = 3.90e-01 uM

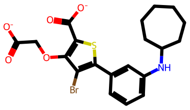
lig_23476
kd = 1.20e-01 uM

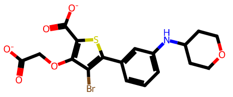
lig_23477
kd = 3.10e-01 uM

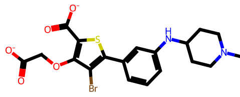
lig_23479
kd = 1.90e-01 uM

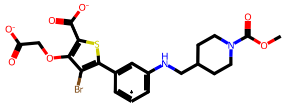
lig_23480
kd = 3.90e-01 uM

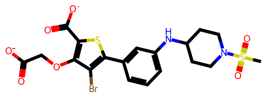
lig_23482
kd = 4.40e-02 uM

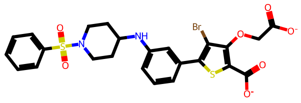
lig_23483
kd = 5.50e-02 uM

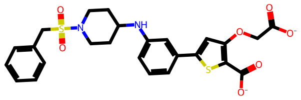
lig_23484
kd = 2.00e+00 uM

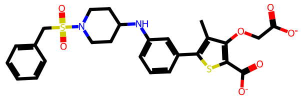
lig_23485
kd = 8.00e-03 uM

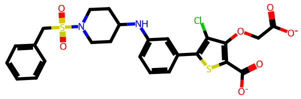
lig_23486
kd = 6.80e-04 uM

Figure S22: Ligands of the PTP1B set.

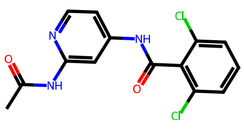

lig_ejm_31
kd = 9.60e-02 uM

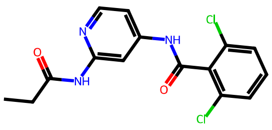

lig_ejm_42
kd = 6.40e-02 uM

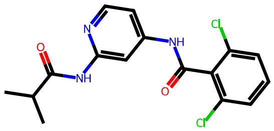

lig_ejm_43
kd = 8.40e-01 uM

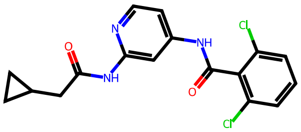

lig_ejm_45
kd = 9.40e-02 uM

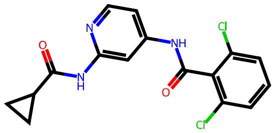

lig_ejm_46
kd = 4.80e-03 uM

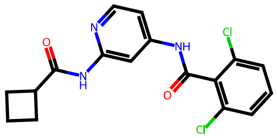

lig_ejm_47
kd = 7.40e-02 uM

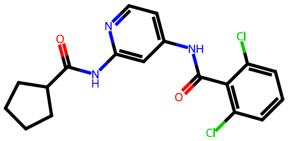

lig_ejm_48
kd = 2.40e-01 uM

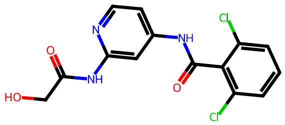

lig_ejm_50
kd = 2.50e-01 uM

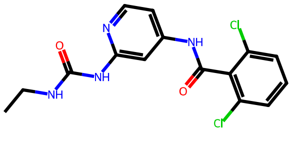

lig_ejm_54
kd = 1.80e-02 uM

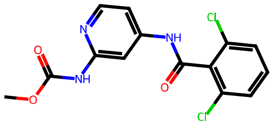

lig_ejm_55
kd = 1.70e-01 uM

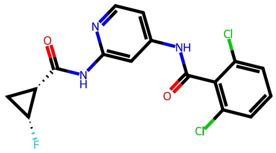

lig_jmc_23
kd = 2.50e+00 nM

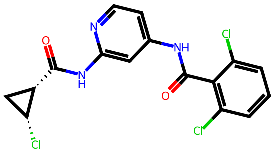

lig_jmc_27
kd = 5.10e+00 nM

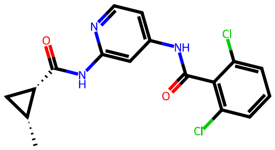

lig_jmc_28
kd = 8.50e+00 nM

Figure S23: Ligands of the TYK2 set.

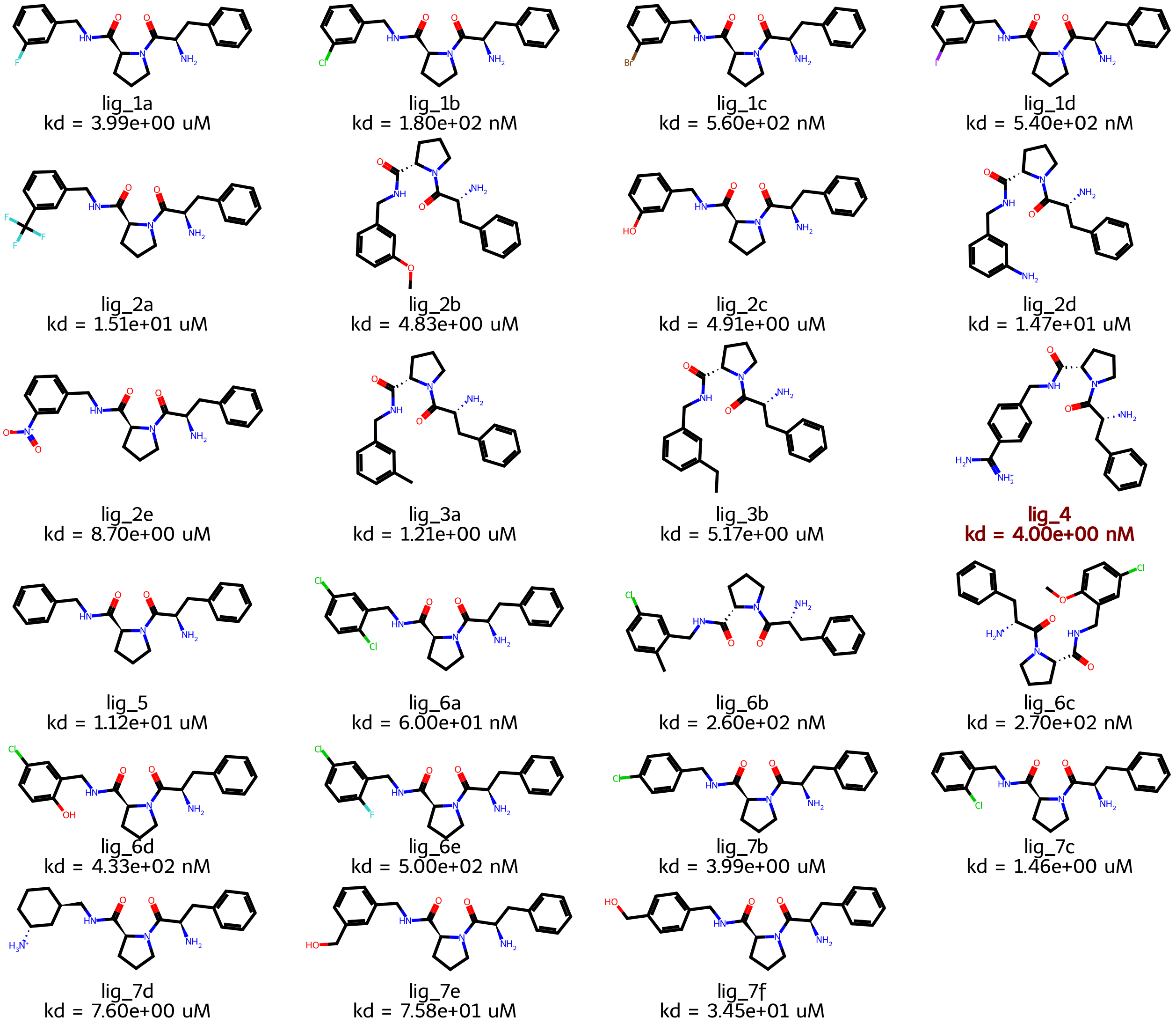


Figure S24: Ligands of the Thrombin set. The outlier lig_4 with bold red font.

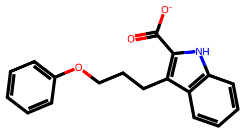

lig_27
kd = 3.50e+01 uM

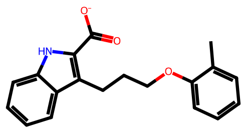

lig_28
kd = 1.50e+01 uM

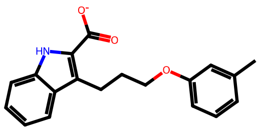

lig_30
kd = 1.90e+00 uM

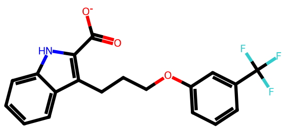

lig_31
kd = 1.70e+00 uM

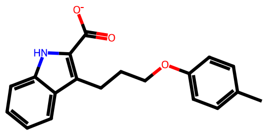

lig_32
kd = 1.60e+01 uM

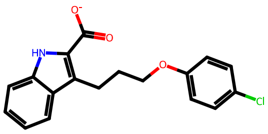

lig_33
kd = 9.80e+00 uM

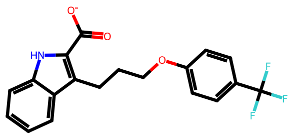

lig_34
kd = 9.90e+00 uM

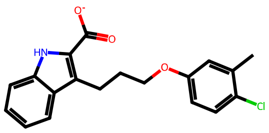

lig_35
kd = 3.80e-01 uM

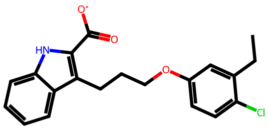

lig_36
kd = 1.10e+00 uM

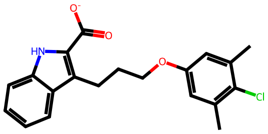

lig_37
kd = 3.00e-01 uM

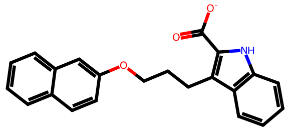

lig_43
kd = 7.50e+00 uM

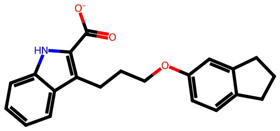

lig_46
kd = 2.90e+00 uM

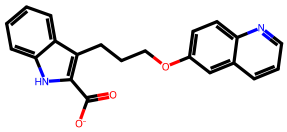

lig_47
kd = 6.20e+01 uM

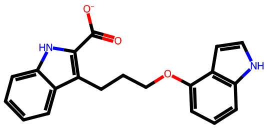

lig_48
kd = 1.40e+01 uM

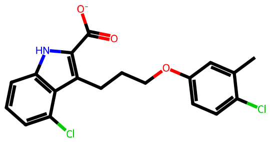

lig_49
kd = 8.10e-01 uM

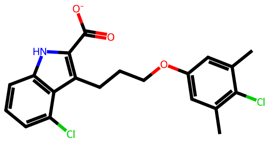

lig_50
kd = 1.60e-01 uM

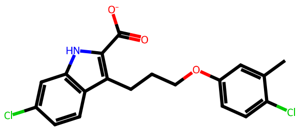

lig_52
kd = 1.90e-01 uM

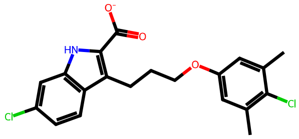

lig_53
kd = 5.50e-02 uM

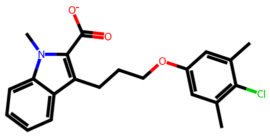

lig_56
kd = 1.80e-01 uM

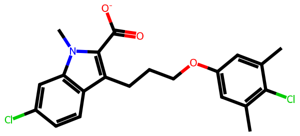

lig_58
kd = 1.40e-01 uM

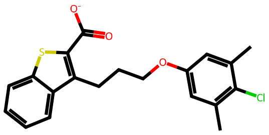

lig_60
kd = 3.20e-01 uM

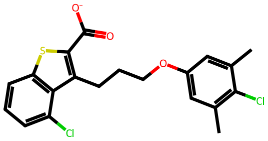

lig_61
kd = 1.30e+00 uM

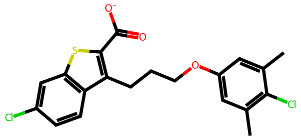

lig_63
kd = 2.50e-01 uM

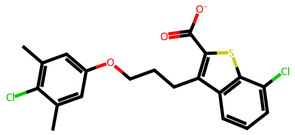

lig_65
kd = 7.50e-01 uM

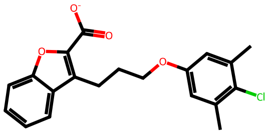

lig_67
kd = 3.00e+00 uM

Figure S25: Ligands of the MCL1 set.

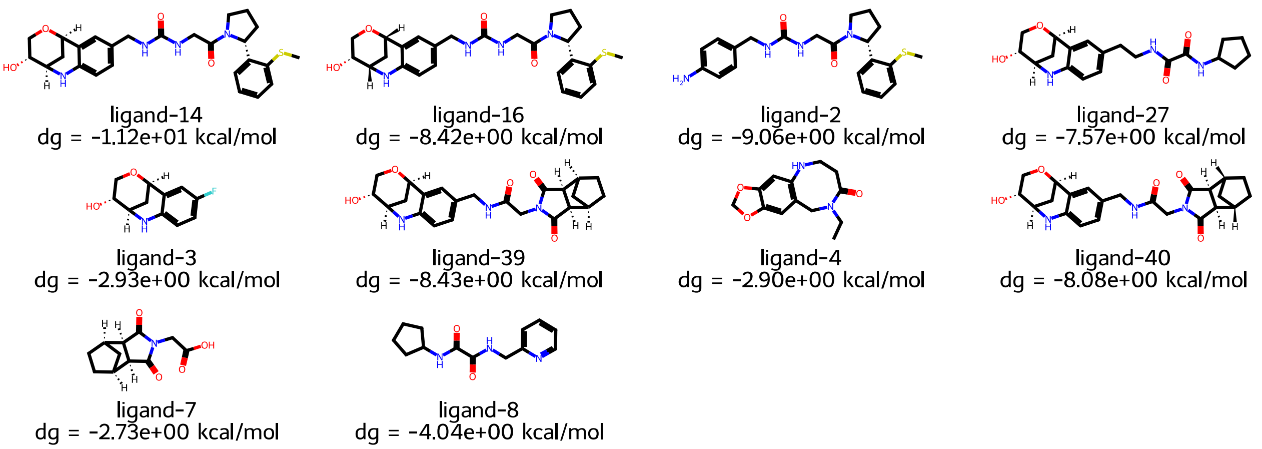


Figure S26: Ligands of the CyclophilinD set.

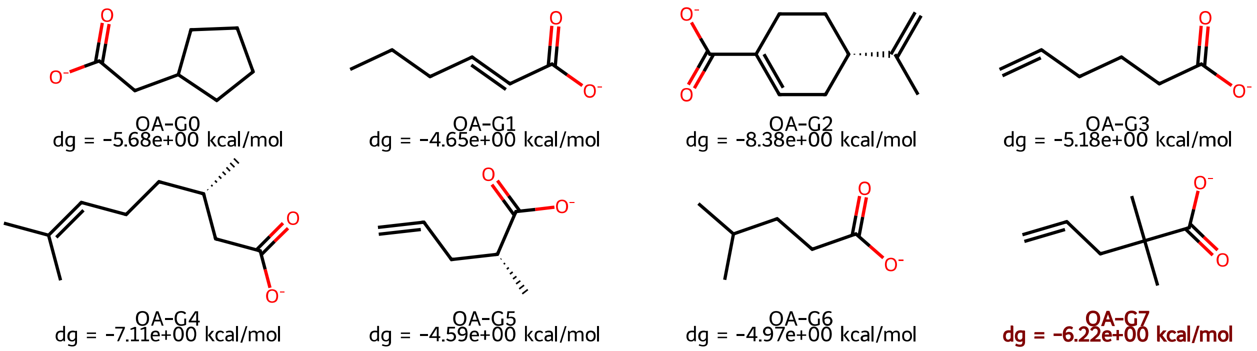


Figure S27: Ligands from the SAMPL6-OA set. OA-G7 ligand, highlighted by bold red font, was excluded from our analysis since FEP simulations were unstable.

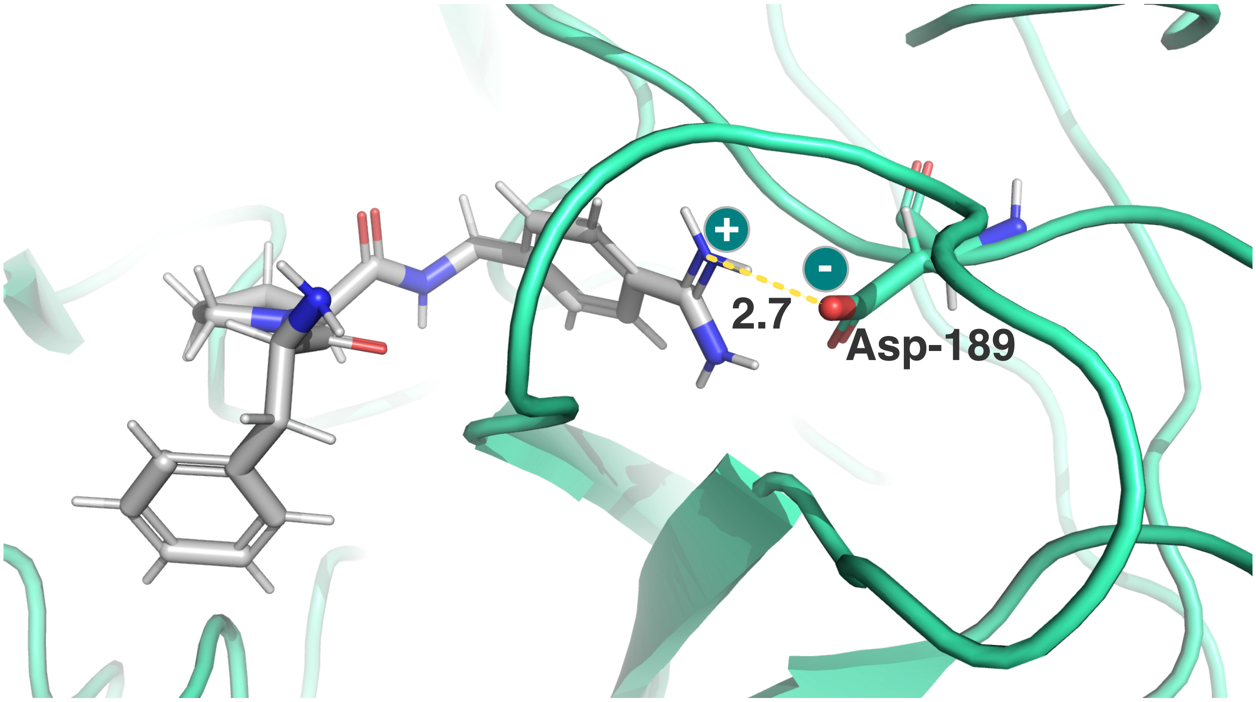


Figure S28: Salt bridge formation between the cationic amidinium moiety of lig_4 and the carboxylate group of Asp-189 in the lig_4/Thrombin complex. The distance between the two groups is approximately 2.7 Å.

Table S1: Agreement between $\Delta G_{\text{calc}}$ and $\Delta G_{\text{exp}}$ for eight receptors, quantified by Pearson $\rho$, Kendall's $\tau$, Spearman $r_S$, RMSE, MSE, and MUE. Color shades from blue to red highlight small to large values to guide the eye. Minor differences of confidence intervals (if any) relative to the insets of correlation plots are caused by different random seeds for bootstrapping.

| System | Simulation | Force Field | $\rho$ | $\tau$ | $r_S$ | RMSE | MSE | MUE |
|---|---|---|---|---|---|---|---|---|
| P38 | FEP | Espaloma | $0.28^{0.48}_{0.11}$ | $0.29^{0.44}_{0.14}$ | $0.37^{0.57}_{0.15}$ | $3.23^{3.53}_{2.90}$ | $-2.74^{-2.43}_{-3.06}$ | $2.85^{3.14}_{2.58}$ |
| | | GAFF | $0.61^{0.71}_{0.49}$ | $0.41^{0.53}_{0.29}$ | $0.57^{0.69}_{0.42}$ | $2.18^{2.36}_{1.99}$ | $-1.87^{-1.66}_{-2.07}$ | $1.92^{2.11}_{1.72}$ |
| | | OpenFF | $0.58^{0.69}_{0.46}$ | $0.40^{0.50}_{0.30}$ | $0.59^{0.69}_{0.46}$ | $2.92^{3.12}_{2.70}$ | $-2.62^{-2.39}_{-2.86}$ | $2.62^{2.86}_{2.39}$ |
| | MMGBSA | Espaloma | $0.39^{0.52}_{0.25}$ | $0.28^{0.38}_{0.17}$ | $0.41^{0.54}_{0.25}$ | $31.98^{32.69}_{31.30}$ | $-31.75^{-31.07}_{-32.48}$ | $31.75^{32.48}_{31.07}$ |
| | | GAFF | $0.38^{0.57}_{0.18}$ | $0.29^{0.42}_{0.17}$ | $0.43^{0.58}_{0.25}$ | $31.27^{31.92}_{30.59}$ | $-31.05^{-30.38}_{-31.71}$ | $31.05^{31.71}_{30.38}$ |
| | | OpenFF | $0.41^{0.57}_{0.26}$ | $0.34^{0.47}_{0.21}$ | $0.46^{0.62}_{0.28}$ | $33.06^{33.70}_{32.41}$ | $-32.86^{-32.20}_{-33.53}$ | $32.86^{33.53}_{32.20}$ |
| | MMGBSA-C2 | Espaloma | $0.23^{0.38}_{0.07}$ | $0.18^{0.31}_{0.05}$ | $0.26^{0.43}_{0.08}$ | $17.47^{18.36}_{16.51}$ | $-16.50^{-15.41}_{-17.56}$ | $16.50^{17.56}_{15.41}$ |
| | | GAFF | $0.24^{0.45}_{0.02}$ | $0.18^{0.33}_{0.02}$ | $0.21^{0.42}_{-0.00}$ | $15.40^{16.31}_{14.44}$ | $-14.59^{-13.68}_{-15.52}$ | $14.59^{15.52}_{13.68}$ |
| | | OpenFF | $0.27^{0.46}_{0.10}$ | $0.25^{0.38}_{0.12}$ | $0.35^{0.52}_{0.16}$ | $16.70^{17.58}_{15.77}$ | $-15.89^{-14.94}_{-16.83}$ | $15.89^{16.83}_{14.94}$ |
| | MMGBSA-IE | Espaloma | $0.36^{0.49}_{0.22}$ | $0.28^{0.38}_{0.17}$ | $0.40^{0.54}_{0.24}$ | $27.39^{28.08}_{26.65}$ | $-27.10^{-26.36}_{-27.83}$ | $27.10^{27.83}_{26.36}$ |
| | | GAFF | $0.37^{0.56}_{0.16}$ | $0.29^{0.43}_{0.16}$ | $0.41^{0.57}_{0.22}$ | $26.36^{27.01}_{25.68}$ | $-26.11^{-25.44}_{-26.77}$ | $26.11^{26.77}_{25.44}$ |
| | | OpenFF | $0.40^{0.57}_{0.23}$ | $0.36^{0.48}_{0.24}$ | $0.49^{0.64}_{0.32}$ | $28.08^{28.70}_{27.42}$ | $-27.85^{-27.19}_{-28.49}$ | $27.85^{28.49}_{27.19}$ |
| | MMPBSA | Espaloma | $0.29^{0.43}_{0.16}$ | $0.22^{0.33}_{0.11}$ | $0.33^{0.48}_{0.17}$ | $20.11^{20.70}_{19.50}$ | $-19.84^{-19.24}_{-20.44}$ | $19.84^{20.44}_{19.24}$ |
| | | GAFF | $0.35^{0.54}_{0.17}$ | $0.27^{0.39}_{0.14}$ | $0.41^{0.56}_{0.24}$ | $18.46^{19.06}_{17.83}$ | $-18.19^{-17.60}_{-18.77}$ | $18.19^{18.77}_{17.60}$ |
| | | OpenFF | $0.53^{0.65}_{0.41}$ | $0.43^{0.54}_{0.32}$ | $0.57^{0.69}_{0.42}$ | $19.53^{20.07}_{18.96}$ | $-19.27^{-18.68}_{-19.85}$ | $19.27^{19.85}_{18.68}$ |
| | MMPBSA-C2 | Espaloma | $0.16^{0.30}_{0.02}$ | $0.09^{0.21}_{-0.03}$ | $0.15^{0.31}_{-0.04}$ | $6.46^{7.12}_{5.75}$ | $-4.60^{-3.75}_{-5.45}$ | $5.51^{6.15}_{4.89}$ |
| | | GAFF | $0.20^{0.39}_{-0.01}$ | $0.10^{0.24}_{-0.05}$ | $0.14^{0.34}_{-0.06}$ | $4.85^{5.47}_{4.13}$ | $-1.73^{-0.89}_{-2.56}$ | $3.65^{4.25}_{3.06}$ |
| | | OpenFF | $0.38^{0.57}_{0.20}$ | $0.34^{0.47}_{0.21}$ | $0.46^{0.62}_{0.29}$ | $4.96^{5.46}_{4.40}$ | $-2.30^{-1.49}_{-3.12}$ | $4.09^{4.61}_{3.58}$ |
| | MMPBSA-IE | Espaloma | $0.28^{0.41}_{0.15}$ | $0.20^{0.31}_{0.09}$ | $0.30^{0.45}_{0.13}$ | $15.53^{16.10}_{14.91}$ | $-15.19^{-14.59}_{-15.78}$ | $15.19^{15.78}_{14.59}$ |
| | | GAFF | $0.33^{0.53}_{0.15}$ | $0.30^{0.43}_{0.18}$ | $0.45^{0.59}_{0.27}$ | $13.61^{14.21}_{12.97}$ | $-13.25^{-12.67}_{-13.82}$ | $13.25^{13.82}_{12.67}$ |
| | | OpenFF | $0.53^{0.66}_{0.39}$ | $0.43^{0.55}_{0.31}$ | $0.58^{0.71}_{0.42}$ | $14.58^{15.09}_{14.05}$ | $-14.26^{-13.70}_{-14.82}$ | $14.26^{14.82}_{13.70}$ |
| A2A | FEP | Espaloma | $0.54^{0.74}_{0.23}$ | $0.16^{0.44}_{-0.12}$ | $0.21^{0.55}_{-0.16}$ | $4.07^{4.54}_{3.55}$ | $-3.73^{-3.21}_{-4.25}$ | $3.73^{4.25}_{3.21}$ |
| | | GAFF | $0.93^{0.97}_{0.87}$ | $0.69^{0.85}_{0.53}$ | $0.85^{0.95}_{0.69}$ | $3.66^{4.10}_{3.17}$ | $-3.28^{-2.76}_{-3.80}$ | $3.28^{3.80}_{2.76}$ |
| | | OpenFF | $0.90^{0.96}_{0.73}$ | $0.56^{0.76}_{0.35}$ | $0.76^{0.89}_{0.55}$ | $4.05^{4.35}_{3.72}$ | $-3.92^{-3.60}_{-4.24}$ | $3.92^{4.24}_{3.60}$ |
| | MMGBSA | Espaloma | $0.94^{0.98}_{0.87}$ | $0.73^{0.90}_{0.56}$ | $0.88^{0.96}_{0.70}$ | $28.76^{29.21}_{28.32}$ | $-28.73^{-28.29}_{-29.18}$ | $28.73^{29.18}_{28.29}$ |
| | | GAFF | $0.90^{0.95}_{0.74}$ | $0.64^{0.80}_{0.49}$ | $0.81^{0.90}_{0.64}$ | $26.68^{27.26}_{26.07}$ | $-26.61^{-26.00}_{-27.21}$ | $26.61^{27.21}_{26.00}$ |
| | | OpenFF | $0.91^{0.96}_{0.73}$ | $0.60^{0.77}_{0.43}$ | $0.75^{0.89}_{0.52}$ | $29.63^{30.14}_{29.09}$ | $-29.58^{-29.04}_{-30.11}$ | $29.58^{30.11}_{29.04}$ |
| | MMGBSA-C2 | Espaloma | $0.93^{0.97}_{0.79}$ | $0.60^{0.79}_{0.41}$ | $0.81^{0.90}_{0.61}$ | $19.57^{20.04}_{19.08}$ | $-19.51^{-19.01}_{-20.00}$ | $19.51^{20.00}_{19.01}$ |
| | | GAFF | $0.89^{0.96}_{0.74}$ | $0.64^{0.81}_{0.48}$ | $0.79^{0.92}_{0.61}$ | $17.42^{18.05}_{16.79}$ | $-17.30^{-16.66}_{-17.97}$ | $17.30^{17.97}_{16.66}$ |
| | | OpenFF | $0.95^{0.98}_{0.82}$ | $0.60^{0.81}_{0.38}$ | $0.76^{0.91}_{0.51}$ | $19.91^{20.45}_{19.37}$ | $-19.83^{-19.26}_{-20.41}$ | $19.83^{20.41}_{19.26}$ |
| | MMGBSA-IE | Espaloma | $0.94^{0.97}_{0.83}$ | $0.69^{0.86}_{0.51}$ | $0.85^{0.95}_{0.68}$ | $25.02^{25.44}_{24.57}$ | $-24.98^{-24.53}_{-25.41}$ | $24.98^{25.41}_{24.53}$ |
| | | GAFF | $0.91^{0.96}_{0.78}$ | $0.64^{0.80}_{0.50}$ | $0.81^{0.90}_{0.63}$ | $23.01^{23.56}_{22.42}$ | $-22.93^{-22.34}_{-23.51}$ | $22.93^{23.51}_{22.34}$ |
| | | OpenFF | $0.92^{0.96}_{0.74}$ | $0.56^{0.75}_{0.37}$ | $0.73^{0.88}_{0.50}$ | $25.78^{26.27}_{25.26}$ | $-25.73^{-25.20}_{-26.24}$ | $25.73^{26.24}_{25.20}$ |
| | MMPBSA | Espaloma | $0.65^{0.81}_{0.43}$ | $0.47^{0.76}_{0.17}$ | $0.56^{0.86}_{0.21}$ | $16.81^{17.35}_{16.25}$ | $-16.72^{-16.18}_{-17.26}$ | $16.72^{17.26}_{16.18}$ |
| | | GAFF | $0.84^{0.92}_{0.58}$ | $0.51^{0.77}_{0.26}$ | $0.64^{0.87}_{0.33}$ | $15.05^{15.33}_{14.77}$ | $-15.02^{-14.74}_{-15.30}$ | $15.02^{15.30}_{14.74}$ |

| System | Simulation | Force Field | $\rho$ | $\tau$ | $r_S$ | RMSE | MSE | MUE |
|---|---|---|---|---|---|---|---|---|
| | | OpenFF | $0.70^{0.85}_{0.21}$ | $0.38^{0.66}_{0.10}$ | $0.47^{0.78}_{0.10}$ | $17.83^{18.16}_{17.49}$ | $-17.79^{-17.45}_{-18.14}$ | $17.79^{18.14}_{17.45}$ |
| | MMPBSA-C2 | Espaloma | $0.63^{0.84}_{0.32}$ | $0.42^{0.73}_{0.10}$ | $0.50^{0.84}_{0.14}$ | $7.76^{8.35}_{7.12}$ | $-7.51^{-6.89}_{-8.13}$ | $7.51^{8.13}_{6.89}$ |
| | | GAFF | $0.74^{0.86}_{0.50}$ | $0.56^{0.79}_{0.33}$ | $0.66^{0.89}_{0.39}$ | $5.86^{6.26}_{5.42}$ | $-5.72^{-5.31}_{-6.12}$ | $5.72^{6.12}_{5.31}$ |
| | | OpenFF | $0.71^{0.88}_{0.29}$ | $0.38^{0.67}_{0.10}$ | $0.54^{0.80}_{0.18}$ | $8.15^{8.56}_{7.71}$ | $-8.05^{-7.63}_{-8.45}$ | $8.05^{8.45}_{7.63}$ |
| | MMPBSA-IE | Espaloma | $0.64^{0.82}_{0.36}$ | $0.47^{0.76}_{0.18}$ | $0.56^{0.85}_{0.21}$ | $13.09^{13.62}_{12.54}$ | $-12.97^{-12.44}_{-13.52}$ | $12.97^{13.52}_{12.44}$ |
| | | GAFF | $0.81^{0.90}_{0.57}$ | $0.56^{0.80}_{0.30}$ | $0.68^{0.90}_{0.40}$ | $11.38^{11.69}_{11.06}$ | $-11.34^{-11.03}_{-11.65}$ | $11.34^{11.65}_{11.03}$ |
| | | OpenFF | $0.68^{0.84}_{0.20}$ | $0.33^{0.63}_{0.03}$ | $0.44^{0.76}_{0.05}$ | $13.99^{14.32}_{13.66}$ | $-13.95^{-13.62}_{-14.28}$ | $13.95^{14.28}_{13.62}$ |
| PTP1B | FEP | Espaloma | $0.40^{0.62}_{0.18}$ | $0.39^{0.58}_{0.22}$ | $0.52^{0.71}_{0.31}$ | $14.44^{14.93}_{13.92}$ | $-14.22^{-13.70}_{-14.74}$ | $14.22^{14.74}_{13.70}$ |
| | | GAFF | $0.59^{0.71}_{0.41}$ | $0.36^{0.50}_{0.22}$ | $0.48^{0.64}_{0.29}$ | $10.50^{10.94}_{10.05}$ | $-10.28^{-9.84}_{-10.73}$ | $10.28^{10.73}_{9.84}$ |
| | | OpenFF | $0.67^{0.76}_{0.58}$ | $0.43^{0.55}_{0.31}$ | $0.61^{0.73}_{0.45}$ | $14.48^{14.98}_{13.95}$ | $-14.27^{-13.73}_{-14.79}$ | $14.27^{14.79}_{13.73}$ |
| | MMGBSA | Espaloma | $0.68^{0.84}_{0.51}$ | $0.53^{0.67}_{0.40}$ | $0.67^{0.83}_{0.49}$ | $75.43^{77.36}_{73.44}$ | $-74.90^{-73.00}_{-76.79}$ | $74.90^{76.79}_{73.00}$ |
| | | GAFF | $0.63^{0.92}_{0.39}$ | $0.58^{0.72}_{0.44}$ | $0.68^{0.85}_{0.49}$ | $77.62^{79.40}_{75.80}$ | $-77.22^{-75.53}_{-78.89}$ | $77.22^{78.89}_{75.53}$ |
| | | OpenFF | $0.70^{0.90}_{0.51}$ | $0.60^{0.74}_{0.46}$ | $0.72^{0.89}_{0.54}$ | $77.68^{79.81}_{75.55}$ | $-77.11^{-75.12}_{-79.14}$ | $77.11^{79.14}_{75.12}$ |
| | MMGBSA-C2 | Espaloma | $0.41^{0.59}_{0.24}$ | $0.34^{0.50}_{0.19}$ | $0.47^{0.65}_{0.25}$ | $75.44^{78.18}_{72.59}$ | $74.28^{77.05}_{71.47}$ | $74.28^{77.05}_{71.47}$ |
| | | GAFF | $0.09^{0.32}_{-0.19}$ | $0.11^{0.31}_{-0.08}$ | $0.09^{0.35}_{-0.17}$ | $79.13^{81.01}_{77.22}$ | $78.64^{80.55}_{76.75}$ | $78.64^{80.55}_{76.75}$ |
| | | OpenFF | $0.38^{0.58}_{0.06}$ | $0.17^{0.33}_{0.00}$ | $0.21^{0.42}_{-0.02}$ | $78.06^{80.01}_{76.00}$ | $77.46^{79.49}_{75.36}$ | $77.46^{79.49}_{75.36}$ |
| | MMGBSA-IE | Espaloma | $0.67^{0.83}_{0.51}$ | $0.55^{0.69}_{0.41}$ | $0.68^{0.84}_{0.50}$ | $58.76^{60.66}_{56.83}$ | $-58.12^{-56.29}_{-59.99}$ | $58.12^{59.99}_{56.29}$ |
| | | GAFF | $0.61^{0.91}_{0.38}$ | $0.53^{0.69}_{0.38}$ | $0.64^{0.82}_{0.44}$ | $60.41^{62.09}_{58.67}$ | $-59.94^{-58.35}_{-61.51}$ | $59.94^{61.51}_{58.35}$ |
| | | OpenFF | $0.69^{0.90}_{0.50}$ | $0.60^{0.74}_{0.46}$ | $0.71^{0.87}_{0.54}$ | $60.76^{62.88}_{58.64}$ | $-60.06^{-58.13}_{-62.07}$ | $60.06^{62.07}_{58.13}$ |
| | MMPBSA | Espaloma | $0.62^{0.74}_{0.48}$ | $0.44^{0.56}_{0.32}$ | $0.61^{0.75}_{0.44}$ | $67.97^{68.87}_{67.07}$ | $-67.84^{-66.96}_{-68.74}$ | $67.84^{68.74}_{66.96}$ |
| | | GAFF | $0.48^{0.89}_{0.23}$ | $0.58^{0.72}_{0.44}$ | $0.68^{0.84}_{0.50}$ | $70.74^{71.52}_{69.93}$ | $-70.65^{-69.89}_{-71.39}$ | $70.65^{71.39}_{69.89}$ |
| | | OpenFF | $0.70^{0.86}_{0.53}$ | $0.54^{0.69}_{0.39}$ | $0.67^{0.82}_{0.49}$ | $70.31^{71.19}_{69.40}$ | $-70.19^{-69.32}_{-71.05}$ | $70.19^{71.05}_{69.32}$ |
| | MMPBSA-C2 | Espaloma | $0.16^{0.40}_{-0.05}$ | $0.28^{0.46}_{0.11}$ | $0.37^{0.60}_{0.13}$ | $82.33^{85.06}_{79.47}$ | $81.34^{84.03}_{78.59}$ | $81.34^{84.03}_{78.59}$ |
| | | GAFF | $-0.27^{-0.02}_{-0.50}$ | $0.06^{0.27}_{-0.14}$ | $0.02^{0.29}_{-0.25}$ | $85.84^{88.10}_{83.42}$ | $85.21^{87.40}_{82.93}$ | $85.21^{87.40}_{82.93}$ |
| | | OpenFF | $0.00^{0.30}_{-0.32}$ | $-0.01^{0.19}_{-0.20}$ | $-0.04^{0.22}_{-0.30}$ | $85.02^{87.20}_{82.77}$ | $84.38^{86.56}_{82.20}$ | $84.38^{86.56}_{82.20}$ |
| | MMPBSA-IE | Espaloma | $0.57^{0.69}_{0.44}$ | $0.42^{0.55}_{0.29}$ | $0.57^{0.72}_{0.39}$ | $51.24^{52.09}_{50.35}$ | $-51.07^{-50.19}_{-51.92}$ | $51.07^{51.92}_{50.19}$ |
| | | GAFF | $0.39^{0.74}_{0.13}$ | $0.42^{0.57}_{0.27}$ | $0.55^{0.72}_{0.36}$ | $53.49^{54.31}_{52.67}$ | $-53.37^{-52.60}_{-54.14}$ | $53.37^{54.14}_{52.60}$ |
| | | OpenFF | $0.67^{0.84}_{0.48}$ | $0.52^{0.66}_{0.37}$ | $0.65^{0.80}_{0.46}$ | $53.30^{54.19}_{52.38}$ | $-53.14^{-52.27}_{-54.00}$ | $53.14^{54.00}_{52.27}$ |
| TYK2 | FEP | Espaloma | $0.30^{0.55}_{0.07}$ | $0.28^{0.49}_{0.07}$ | $0.42^{0.65}_{0.14}$ | $4.51^{4.80}_{4.19}$ | $-4.36^{-4.04}_{-4.68}$ | $4.36^{4.68}_{4.04}$ |
| | | GAFF | $0.58^{0.77}_{0.38}$ | $0.51^{0.74}_{0.30}$ | $0.62^{0.87}_{0.35}$ | $1.05^{1.21}_{0.87}$ | $0.17^{0.46}_{-0.12}$ | $0.79^{0.99}_{0.60}$ |
| | | OpenFF | $0.50^{0.66}_{0.35}$ | $0.26^{0.44}_{0.07}$ | $0.48^{0.66}_{0.20}$ | $2.10^{2.33}_{1.84}$ | $-1.78^{-1.47}_{-2.09}$ | $1.86^{2.13}_{1.59}$ |
| | MMGBSA | Espaloma | $0.57^{0.73}_{0.43}$ | $0.41^{0.57}_{0.25}$ | $0.59^{0.73}_{0.38}$ | $28.40^{28.79}_{28.00}$ | $-28.37^{-27.97}_{-28.76}$ | $28.37^{28.76}_{27.97}$ |
| | | GAFF | $0.76^{0.95}_{0.58}$ | $0.54^{0.78}_{0.32}$ | $0.64^{0.92}_{0.36}$ | $26.10^{26.45}_{25.74}$ | $-26.07^{-25.72}_{-26.42}$ | $26.07^{26.42}_{25.72}$ |
| | | OpenFF | $0.69^{0.81}_{0.55}$ | $0.59^{0.75}_{0.44}$ | $0.74^{0.87}_{0.55}$ | $28.97^{29.27}_{28.68}$ | $-28.95^{-28.66}_{-29.25}$ | $28.95^{29.25}_{28.66}$ |
| | MMGBSA-C2 | Espaloma | $0.40^{0.63}_{0.22}$ | $0.26^{0.53}_{-0.01}$ | $0.31^{0.65}_{-0.04}$ | $17.30^{18.01}_{16.58}$ | $-16.99^{-16.11}_{-17.91}$ | $16.99^{17.91}_{16.11}$ |
| | | GAFF | $0.41^{0.62}_{0.27}$ | $0.44^{0.69}_{0.16}$ | $0.47^{0.76}_{0.15}$ | $15.29^{16.06}_{14.50}$ | $-14.43^{-13.06}_{-15.85}$ | $14.72^{15.86}_{13.61}$ |
| | | OpenFF | $0.39^{0.56}_{0.25}$ | $0.33^{0.53}_{0.14}$ | $0.50^{0.69}_{0.22}$ | $17.68^{18.44}_{16.89}$ | $-17.27^{-16.21}_{-18.33}$ | $17.27^{18.33}_{16.21}$ |
| | MMGBSA-IE | Espaloma | $0.58^{0.76}_{0.41}$ | $0.38^{0.58}_{0.19}$ | $0.52^{0.73}_{0.26}$ | $24.31^{24.70}_{23.90}$ | $-24.27^{-23.85}_{-24.67}$ | $24.27^{24.67}_{23.85}$ |
| | | GAFF | $0.69^{0.89}_{0.50}$ | $0.59^{0.84}_{0.35}$ | $0.64^{0.93}_{0.35}$ | $21.94^{22.37}_{21.49}$ | $-21.88^{-21.43}_{-22.32}$ | $21.88^{22.32}_{21.43}$ |
| | | OpenFF | $0.59^{0.74}_{0.47}$ | $0.46^{0.64}_{0.29}$ | $0.62^{0.80}_{0.38}$ | $24.86^{25.24}_{24.46}$ | $-24.82^{-24.42}_{-25.21}$ | $24.82^{25.21}_{24.42}$ |

| System | Simulation | Force Field | $\rho$ | $\tau$ | $r_S$ | RMSE | MSE | MUE |
|---|---|---|---|---|---|---|---|---|
| | MMPBSA | Espaloma | $0.65^{0.80}_{0.53}$ | $0.44^{0.58}_{0.31}$ | $0.65^{0.75}_{0.49}$ | $20.43^{20.76}_{20.10}$ | $-20.40^{-20.06}_{-20.74}$ | $20.40^{20.74}_{20.06}$ |
| | | GAFF | $0.81^{0.95}_{0.68}$ | $0.64^{0.84}_{0.45}$ | $0.76^{0.94}_{0.56}$ | $18.21^{18.54}_{17.86}$ | $-18.16^{-17.83}_{-18.50}$ | $18.16^{18.50}_{17.83}$ |
| | | OpenFF | $0.76^{0.84}_{0.69}$ | $0.56^{0.69}_{0.44}$ | $0.77^{0.85}_{0.62}$ | $20.80^{21.07}_{20.52}$ | $-20.77^{-20.50}_{-21.05}$ | $20.77^{21.05}_{20.50}$ |
| | MMPBSA-C2 | Espaloma | $0.44^{0.66}_{0.27}$ | $0.28^{0.56}_{0.01}$ | $0.32^{0.65}_{-0.03}$ | $9.51^{10.06}_{8.92}$ | $-9.02^{-8.21}_{-9.85}$ | $9.02^{9.85}_{8.21}$ |
| | | GAFF | $0.45^{0.66}_{0.32}$ | $0.46^{0.70}_{0.21}$ | $0.52^{0.79}_{0.23}$ | $8.10^{8.59}_{7.58}$ | $-6.53^{-5.21}_{-7.86}$ | $7.88^{8.39}_{7.36}$ |
| | | OpenFF | $0.45^{0.62}_{0.35}$ | $0.41^{0.56}_{0.26}$ | $0.60^{0.74}_{0.38}$ | $9.73^{10.25}_{9.18}$ | $-9.09^{-8.11}_{-10.07}$ | $9.39^{10.09}_{8.68}$ |
| | MMPBSA-IE | Espaloma | $0.68^{0.84}_{0.53}$ | $0.46^{0.63}_{0.29}$ | $0.63^{0.78}_{0.42}$ | $16.34^{16.65}_{16.01}$ | $-16.30^{-15.97}_{-16.62}$ | $16.30^{16.62}_{15.97}$ |
| | | GAFF | $0.76^{0.92}_{0.60}$ | $0.69^{0.88}_{0.51}$ | $0.77^{0.94}_{0.56}$ | $14.06^{14.47}_{13.64}$ | $-13.98^{-13.56}_{-14.40}$ | $13.98^{14.40}_{13.56}$ |
| | | OpenFF | $0.70^{0.79}_{0.62}$ | $0.54^{0.67}_{0.41}$ | $0.75^{0.83}_{0.59}$ | $16.68^{17.01}_{16.34}$ | $-16.64^{-16.29}_{-16.98}$ | $16.64^{16.98}_{16.29}$ |
| Thrombin | FEP | Espaloma | $0.66^{0.78}_{0.40}$ | $0.40^{0.52}_{0.27}$ | $0.52^{0.66}_{0.35}$ | $3.49^{4.22}_{2.52}$ | $-1.71^{-1.09}_{-2.33}$ | $2.59^{3.07}_{2.10}$ |
| | | GAFF | $0.47^{0.68}_{-0.05}$ | $0.09^{0.25}_{-0.07}$ | $0.13^{0.34}_{-0.10}$ | $2.56^{2.96}_{2.06}$ | $0.39^{0.91}_{-0.13}$ | $1.97^{2.30}_{1.63}$ |
| | | OpenFF | $0.65^{0.77}_{0.36}$ | $0.30^{0.45}_{0.16}$ | $0.42^{0.60}_{0.22}$ | $2.91^{3.54}_{2.10}$ | $-1.29^{-0.75}_{-1.85}$ | $2.01^{2.46}_{1.57}$ |
| | MMGBSA | Espaloma | $0.25^{0.53}_{-0.25}$ | $-0.04^{0.16}_{-0.24}$ | $-0.08^{0.19}_{-0.34}$ | $33.38^{34.16}_{32.56}$ | $-33.17^{-32.39}_{-33.93}$ | $33.17^{33.93}_{32.39}$ |
| | | GAFF | $0.24^{0.50}_{-0.17}$ | $0.01^{0.20}_{-0.18}$ | $0.01^{0.26}_{-0.23}$ | $30.51^{31.22}_{29.79}$ | $-30.30^{-29.60}_{-31.03}$ | $30.30^{31.03}_{29.60}$ |
| | | OpenFF | $0.41^{0.61}_{0.11}$ | $0.17^{0.35}_{-0.02}$ | $0.24^{0.47}_{-0.01}$ | $32.53^{33.32}_{31.73}$ | $-32.31^{-31.51}_{-33.11}$ | $32.31^{33.11}_{31.51}$ |
| | MMGBSA-C2 | Espaloma | $0.01^{0.24}_{-0.26}$ | $0.01^{0.18}_{-0.17}$ | $-0.01^{0.25}_{-0.25}$ | $18.39^{19.30}_{17.45}$ | $-16.25^{-14.43}_{-18.08}$ | $17.68^{18.74}_{16.64}$ |
| | | GAFF | $0.03^{0.26}_{-0.24}$ | $0.02^{0.20}_{-0.15}$ | $0.04^{0.28}_{-0.20}$ | $15.38^{16.54}_{14.12}$ | $-11.43^{-9.24}_{-13.56}$ | $14.38^{15.53}_{13.24}$ |
| | | OpenFF | $0.06^{0.23}_{-0.25}$ | $0.07^{0.23}_{-0.09}$ | $0.10^{0.32}_{-0.13}$ | $18.18^{19.70}_{16.48}$ | $-13.59^{-11.05}_{-16.13}$ | $16.96^{18.29}_{15.60}$ |
| | MMGBSA-IE | Espaloma | $0.26^{0.50}_{-0.10}$ | $0.06^{0.24}_{-0.11}$ | $0.07^{0.31}_{-0.17}$ | $28.48^{29.24}_{27.72}$ | $-28.25^{-27.51}_{-29.00}$ | $28.25^{29.00}_{27.51}$ |
| | | GAFF | $0.24^{0.46}_{-0.04}$ | $0.11^{0.30}_{-0.08}$ | $0.15^{0.39}_{-0.10}$ | $25.37^{26.03}_{24.69}$ | $-25.16^{-24.47}_{-25.84}$ | $25.16^{25.84}_{24.47}$ |
| | | OpenFF | $0.30^{0.48}_{0.09}$ | $0.23^{0.39}_{0.06}$ | $0.29^{0.51}_{0.06}$ | $27.36^{28.16}_{26.52}$ | $-26.99^{-26.05}_{-27.90}$ | $26.99^{27.90}_{26.05}$ |
| | MMPBSA | Espaloma | $0.35^{0.60}_{-0.42}$ | $-0.21^{-0.02}_{-0.40}$ | $-0.23^{0.03}_{-0.48}$ | $18.97^{20.14}_{17.72}$ | $-18.37^{-17.37}_{-19.34}$ | $18.37^{19.34}_{17.37}$ |
| | | GAFF | $0.41^{0.63}_{-0.17}$ | $-0.03^{0.15}_{-0.21}$ | $-0.07^{0.18}_{-0.31}$ | $16.62^{17.58}_{15.62}$ | $-16.11^{-15.27}_{-16.98}$ | $16.11^{16.98}_{15.27}$ |
| | | OpenFF | $0.48^{0.68}_{-0.12}$ | $0.09^{0.29}_{-0.11}$ | $0.11^{0.37}_{-0.14}$ | $18.61^{19.61}_{17.49}$ | $-18.13^{-17.25}_{-18.98}$ | $18.13^{18.98}_{17.25}$ |
| | MMPBSA-C2 | Espaloma | $0.12^{0.28}_{-0.07}$ | $-0.03^{0.14}_{-0.20}$ | $-0.01^{0.23}_{-0.25}$ | $7.18^{8.90}_{4.78}$ | $-1.44^{0.02}_{-2.87}$ | $4.98^{6.05}_{3.92}$ |
| | | GAFF | $0.16^{0.32}_{-0.01}$ | $0.12^{0.28}_{-0.04}$ | $0.16^{0.38}_{-0.07}$ | $8.82^{11.87}_{3.66}$ | $2.76^{4.48}_{0.98}$ | $4.28^{5.88}_{2.61}$ |
| | | OpenFF | $0.13^{0.23}_{-0.06}$ | $0.04^{0.19}_{-0.11}$ | $0.09^{0.30}_{-0.14}$ | $10.13^{13.87}_{3.49}$ | $0.58^{2.71}_{-1.56}$ | $4.58^{6.46}_{2.65}$ |
| | MMPBSA-IE | Espaloma | $0.40^{0.63}_{-0.25}$ | $-0.05^{0.13}_{-0.23}$ | $-0.03^{0.21}_{-0.27}$ | $14.05^{15.05}_{12.95}$ | $-13.45^{-12.60}_{-14.31}$ | $13.45^{14.31}_{12.60}$ |
| | | GAFF | $0.49^{0.68}_{-0.01}$ | $0.09^{0.27}_{-0.09}$ | $0.11^{0.35}_{-0.14}$ | $11.43^{12.19}_{10.67}$ | $-10.96^{-10.32}_{-11.64}$ | $10.96^{11.64}_{10.32}$ |
| | | OpenFF | $0.44^{0.65}_{-0.05}$ | $0.07^{0.26}_{-0.11}$ | $0.11^{0.36}_{-0.14}$ | $13.42^{14.32}_{12.45}$ | $-12.81^{-11.98}_{-13.64}$ | $12.81^{13.64}_{11.98}$ |
| MCL1 | FEP | Espaloma | $0.74^{0.82}_{0.65}$ | $0.55^{0.67}_{0.45}$ | $0.73^{0.83}_{0.59}$ | $2.35^{2.57}_{2.09}$ | $-1.89^{-1.62}_{-2.16}$ | $2.02^{2.26}_{1.79}$ |
| | | GAFF | $0.73^{0.80}_{0.67}$ | $0.61^{0.68}_{0.53}$ | $0.81^{0.86}_{0.71}$ | $1.71^{1.93}_{1.47}$ | $-0.87^{-0.58}_{-1.16}$ | $1.30^{1.52}_{1.08}$ |
| | | OpenFF | $0.63^{0.73}_{0.52}$ | $0.47^{0.58}_{0.35}$ | $0.64^{0.76}_{0.48}$ | $1.59^{1.75}_{1.41}$ | $-0.84^{-0.57}_{-1.11}$ | $1.33^{1.51}_{1.16}$ |
| | MMGBSA | Espaloma | $0.56^{0.67}_{0.44}$ | $0.44^{0.55}_{0.33}$ | $0.57^{0.69}_{0.42}$ | $37.85^{38.65}_{37.02}$ | $-37.62^{-36.79}_{-38.45}$ | $37.62^{38.45}_{36.79}$ |
| | | GAFF | $0.70^{0.78}_{0.61}$ | $0.54^{0.64}_{0.45}$ | $0.70^{0.79}_{0.58}$ | $38.55^{39.28}_{37.82}$ | $-38.37^{-37.63}_{-39.12}$ | $38.37^{39.12}_{37.63}$ |
| | | OpenFF | $0.57^{0.69}_{0.45}$ | $0.42^{0.53}_{0.31}$ | $0.58^{0.70}_{0.43}$ | $37.59^{38.27}_{36.91}$ | $-37.43^{-36.77}_{-38.13}$ | $37.43^{38.13}_{36.77}$ |
| | MMGBSA-C2 | Espaloma | $0.41^{0.57}_{0.24}$ | $0.31^{0.45}_{0.16}$ | $0.43^{0.60}_{0.23}$ | $11.37^{12.81}_{9.74}$ | $8.01^{9.61}_{6.42}$ | $8.66^{10.13}_{7.21}$ |
| | | GAFF | $0.72^{0.79}_{0.65}$ | $0.57^{0.66}_{0.49}$ | $0.77^{0.84}_{0.65}$ | $11.07^{12.15}_{9.88}$ | $8.93^{10.23}_{7.67}$ | $9.50^{10.64}_{8.40}$ |
| | | OpenFF | $0.69^{0.78}_{0.61}$ | $0.51^{0.61}_{0.41}$ | $0.70^{0.78}_{0.58}$ | $11.94^{13.11}_{10.62}$ | $9.45^{10.90}_{7.99}$ | $10.00^{11.29}_{8.69}$ |
| | MMGBSA-IE | Espaloma | $0.56^{0.67}_{0.43}$ | $0.42^{0.53}_{0.31}$ | $0.55^{0.67}_{0.38}$ | $28.99^{29.83}_{28.14}$ | $-28.67^{-27.81}_{-29.54}$ | $28.67^{29.54}_{27.81}$ |

| System | Simulation | Force Field | $\rho$ | $\tau$ | $r_S$ | RMSE | MSE | MUE |
|---|---|---|---|---|---|---|---|---|
| | | GAFF | $0.73^{0.80}_{0.65}$ | $0.59^{0.67}_{0.50}$ | $0.76^{0.83}_{0.65}$ | $29.44^{30.21}_{28.65}$ | $-29.17^{-28.39}_{-29.96}$ | $29.17^{29.96}_{28.39}$ |
| | | OpenFF | $0.61^{0.72}_{0.51}$ | $0.47^{0.58}_{0.37}$ | $0.65^{0.75}_{0.50}$ | $28.36^{29.09}_{27.62}$ | $-28.12^{-27.38}_{-28.85}$ | $28.12^{28.85}_{27.38}$ |
| | MMPBSA | Espaloma | $0.65^{0.74}_{0.55}$ | $0.51^{0.61}_{0.41}$ | $0.67^{0.76}_{0.54}$ | $32.78^{33.43}_{32.13}$ | $-32.62^{-31.97}_{-33.27}$ | $32.62^{33.27}_{31.97}$ |
| | | GAFF | $0.76^{0.82}_{0.68}$ | $0.59^{0.68}_{0.51}$ | $0.77^{0.84}_{0.67}$ | $32.97^{33.57}_{32.34}$ | $-32.82^{-32.20}_{-33.44}$ | $32.82^{33.44}_{32.20}$ |
| | | OpenFF | $0.65^{0.75}_{0.54}$ | $0.45^{0.55}_{0.34}$ | $0.63^{0.73}_{0.48}$ | $32.23^{32.78}_{31.69}$ | $-32.11^{-31.57}_{-32.67}$ | $32.11^{32.67}_{31.57}$ |
| | MMPBSA-C2 | Espaloma | $0.45^{0.60}_{0.28}$ | $0.35^{0.50}_{0.21}$ | $0.47^{0.63}_{0.29}$ | $14.90^{16.34}_{13.29}$ | $13.01^{14.48}_{11.54}$ | $13.01^{14.48}_{11.54}$ |
| | | GAFF | $0.75^{0.81}_{0.68}$ | $0.59^{0.67}_{0.51}$ | $0.78^{0.84}_{0.68}$ | $15.68^{16.78}_{14.52}$ | $14.47^{15.69}_{13.28}$ | $14.47^{15.69}_{13.28}$ |
| | | OpenFF | $0.71^{0.79}_{0.62}$ | $0.49^{0.59}_{0.39}$ | $0.69^{0.77}_{0.56}$ | $16.34^{17.59}_{15.01}$ | $14.76^{16.16}_{13.38}$ | $14.85^{16.22}_{13.50}$ |
| | MMPBSA-IE | Espaloma | $0.65^{0.74}_{0.55}$ | $0.51^{0.61}_{0.40}$ | $0.66^{0.77}_{0.53}$ | $23.91^{24.58}_{23.23}$ | $-23.66^{-22.98}_{-24.35}$ | $23.66^{24.35}_{22.98}$ |
| | | GAFF | $0.78^{0.84}_{0.71}$ | $0.61^{0.70}_{0.53}$ | $0.80^{0.86}_{0.70}$ | $23.85^{24.53}_{23.21}$ | $-23.63^{-22.98}_{-24.31}$ | $23.63^{24.31}_{22.98}$ |
| | | OpenFF | $0.68^{0.77}_{0.58}$ | $0.53^{0.63}_{0.42}$ | $0.71^{0.80}_{0.57}$ | $23.01^{23.63}_{22.38}$ | $-22.80^{-22.18}_{-23.42}$ | $22.80^{23.42}_{22.18}$ |
| CyclophilinD | FEP | Espaloma | $0.95^{0.98}_{0.92}$ | $0.82^{0.95}_{0.70}$ | $0.93^{0.99}_{0.81}$ | $3.08^{3.52}_{2.55}$ | $-2.75^{-2.31}_{-3.19}$ | $2.75^{3.19}_{2.31}$ |
| | | GAFF | $0.96^{0.98}_{0.93}$ | $0.73^{0.90}_{0.57}$ | $0.85^{0.96}_{0.70}$ | $1.33^{1.54}_{1.07}$ | $-0.96^{-0.66}_{-1.25}$ | $1.11^{1.34}_{0.87}$ |
| | | OpenFF | $0.97^{0.98}_{0.95}$ | $0.78^{0.95}_{0.62}$ | $0.89^{0.99}_{0.77}$ | $2.47^{2.69}_{2.23}$ | $-2.35^{-2.11}_{-2.59}$ | $2.35^{2.59}_{2.11}$ |
| | MMGBSA | Espaloma | $0.91^{0.97}_{0.87}$ | $0.64^{0.79}_{0.51}$ | $0.84^{0.91}_{0.68}$ | $35.37^{38.73}_{31.62}$ | $-33.57^{-30.05}_{-37.06}$ | $33.57^{37.06}_{30.05}$ |
| | | GAFF | $0.89^{0.94}_{0.84}$ | $0.60^{0.76}_{0.46}$ | $0.79^{0.89}_{0.62}$ | $31.05^{33.41}_{28.48}$ | $-30.09^{-27.64}_{-32.54}$ | $30.09^{32.54}_{27.64}$ |
| | | OpenFF | $0.96^{0.98}_{0.94}$ | $0.64^{0.83}_{0.46}$ | $0.82^{0.92}_{0.64}$ | $34.43^{37.08}_{31.53}$ | $-33.18^{-30.23}_{-36.08}$ | $33.18^{36.08}_{30.23}$ |
| | MMGBSA-C2 | Espaloma | $0.53^{0.70}_{0.35}$ | $0.33^{0.58}_{0.10}$ | $0.42^{0.70}_{0.08}$ | $18.19^{23.00}_{11.12}$ | $-4.25^{1.46}_{-10.08}$ | $13.47^{17.26}_{9.59}$ |
| | | GAFF | $0.40^{0.61}_{-0.10}$ | $0.16^{0.45}_{-0.12}$ | $0.21^{0.56}_{-0.18}$ | $26.70^{36.55}_{9.37}$ | $2.68^{11.28}_{-6.11}$ | $15.04^{22.33}_{7.57}$ |
| | | OpenFF | $0.54^{0.73}_{0.38}$ | $0.42^{0.69}_{0.15}$ | $0.56^{0.81}_{0.22}$ | $24.55^{31.59}_{14.27}$ | $-4.63^{3.33}_{-12.51}$ | $18.26^{23.50}_{13.01}$ |
| | MMGBSA-IE | Espaloma | $0.90^{0.96}_{0.86}$ | $0.69^{0.85}_{0.55}$ | $0.87^{0.93}_{0.71}$ | $28.99^{32.12}_{25.47}$ | $-26.93^{-23.54}_{-30.28}$ | $26.93^{30.28}_{23.54}$ |
| | | GAFF | $0.85^{0.92}_{0.79}$ | $0.64^{0.80}_{0.49}$ | $0.82^{0.91}_{0.65}$ | $24.53^{26.80}_{22.04}$ | $-23.12^{-20.55}_{-25.68}$ | $23.12^{25.68}_{20.55}$ |
| | | OpenFF | $0.94^{0.97}_{0.91}$ | $0.64^{0.84}_{0.47}$ | $0.82^{0.92}_{0.64}$ | $28.53^{31.11}_{25.65}$ | $-26.93^{-23.88}_{-29.92}$ | $26.93^{29.92}_{23.88}$ |
| | MMPBSA | Espaloma | $0.92^{0.97}_{0.88}$ | $0.60^{0.76}_{0.45}$ | $0.82^{0.89}_{0.65}$ | $23.58^{26.00}_{20.99}$ | $-22.21^{-19.75}_{-24.76}$ | $22.21^{24.76}_{19.75}$ |
| | | GAFF | $0.82^{0.90}_{0.73}$ | $0.56^{0.71}_{0.40}$ | $0.77^{0.86}_{0.58}$ | $19.96^{21.66}_{18.08}$ | $-19.10^{-17.28}_{-20.93}$ | $19.10^{20.93}_{17.28}$ |
| | | OpenFF | $0.93^{0.97}_{0.88}$ | $0.56^{0.74}_{0.38}$ | $0.77^{0.88}_{0.57}$ | $23.05^{25.06}_{20.88}$ | $-22.00^{-19.83}_{-24.19}$ | $22.00^{24.19}_{19.83}$ |
| | MMPBSA-C2 | Espaloma | $0.46^{0.65}_{0.13}$ | $0.24^{0.51}_{-0.02}$ | $0.30^{0.65}_{-0.08}$ | $16.41^{22.49}_{5.51}$ | $7.10^{11.80}_{2.19}$ | $8.78^{13.25}_{4.17}$ |
| | | GAFF | $0.35^{0.57}_{-0.36}$ | $0.02^{0.32}_{-0.28}$ | $-0.01^{0.40}_{-0.40}$ | $27.68^{37.86}_{9.66}$ | $13.67^{21.51}_{5.70}$ | $15.48^{22.99}_{7.78}$ |
| | | OpenFF | $0.49^{0.68}_{0.25}$ | $0.29^{0.56}_{0.00}$ | $0.39^{0.69}_{0.03}$ | $22.26^{30.59}_{7.01}$ | $6.55^{13.35}_{-0.58}$ | $11.89^{18.00}_{5.49}$ |
| | MMPBSA-IE | Espaloma | $0.93^{0.99}_{0.89}$ | $0.69^{0.85}_{0.55}$ | $0.87^{0.93}_{0.71}$ | $17.14^{19.39}_{14.65}$ | $-15.57^{-13.29}_{-17.93}$ | $15.57^{17.93}_{13.29}$ |
| | | GAFF | $0.83^{0.91}_{0.74}$ | $0.60^{0.76}_{0.46}$ | $0.79^{0.88}_{0.62}$ | $13.39^{15.05}_{11.50}$ | $-12.13^{-10.30}_{-13.98}$ | $12.13^{13.98}_{10.30}$ |
| | | OpenFF | $0.93^{0.97}_{0.88}$ | $0.60^{0.80}_{0.41}$ | $0.78^{0.90}_{0.58}$ | $17.11^{19.03}_{14.96}$ | $-15.76^{-13.64}_{-17.89}$ | $15.76^{17.89}_{13.64}$ |
| SAMPL6-OA | FEP | Espaloma | $0.97^{0.99}_{0.93}$ | $0.71^{1.00}_{0.47}$ | $0.86^{1.00}_{0.67}$ | $2.95^{3.17}_{2.71}$ | $-2.88^{-2.63}_{-3.13}$ | $2.88^{3.13}_{2.63}$ |
| | MMGBSA | Espaloma | $0.91^{0.97}_{0.89}$ | $0.71^{1.00}_{0.56}$ | $0.86^{1.00}_{0.67}$ | $8.92^{9.34}_{8.48}$ | $-8.85^{-8.43}_{-9.27}$ | $8.85^{9.27}_{8.43}$ |
| | MMGBSA-C2 | Espaloma | $0.86^{0.99}_{0.58}$ | $0.71^{1.00}_{0.53}$ | $0.86^{1.00}_{0.67}$ | $4.25^{4.72}_{3.72}$ | $3.89^{4.53}_{3.26}$ | $3.89^{4.53}_{3.26}$ |
| | MMGBSA-IE | Espaloma | $0.93^{0.97}_{0.90}$ | $0.81^{1.00}_{0.58}$ | $0.89^{1.00}_{0.70}$ | $4.81^{5.16}_{4.44}$ | $-4.70^{-4.34}_{-5.07}$ | $4.70^{5.07}_{4.34}$ |
| | MMPBSA | Espaloma | $0.97^{0.99}_{0.96}$ | $0.90^{1.00}_{0.79}$ | $0.96^{1.00}_{0.89}$ | $8.41^{9.04}_{7.71}$ | $-8.23^{-7.56}_{-8.89}$ | $8.23^{8.89}_{7.56}$ |
| | MMPBSA-C2 | Espaloma | $0.90^{0.98}_{0.72}$ | $0.81^{1.00}_{0.65}$ | $0.93^{1.00}_{0.76}$ | $5.13^{5.69}_{4.50}$ | $4.51^{5.46}_{3.59}$ | $4.75^{5.48}_{4.02}$ |
| | MMPBSA-IE | Espaloma | $0.97^{0.99}_{0.95}$ | $0.90^{1.00}_{0.79}$ | $0.96^{1.00}_{0.89}$ | $4.42^{5.00}_{3.77}$ | $-4.09^{-3.40}_{-4.77}$ | $4.09^{4.77}_{3.40}$ |

(7) Ries, B.; Alibay, I.; Anand, N. M.; Biggin, P. C.; Magarkar, A. Automated Absolute

Binding Free Energy Calculation Workflow for Drug Discovery. *Journal of Chemical Information and Modeling* **2024**, *64*, 5357–5364.

(8) Khalak, Y.; Tresadern, G.; Aldeghi, M.; Baumann, H. M.; Mobley, D. L.; de Groot, B. L.; Gapsys, V. Alchemical absolute protein-ligand binding free energies for drug design. *Chemical Science* **2021**, *12*, 13958–13971.

(9) Lin, Z.; Zou, J.; Liu, S.; Peng, C.; Li, Z.; Wan, X.; Fang, D.; Yin, J.; Gobbo, G.; Chen, Y.; others A cloud computing platform for scalable relative and absolute binding free energy predictions: New opportunities and challenges for drug discovery. *Journal of Chemical Information and Modeling* **2021**, *61*, 2720–2732.

(10) Li, Z.; Huang, Y.; Wu, Y.; Chen, J.; Wu, D.; Zhan, C. G.; Luo, H. B. Absolute Binding Free Energy Calculation and Design of a Subnanomolar Inhibitor of Phosphodiesterase-10. *Journal of Medicinal Chemistry* **2019**, *62*, 2099–2111.

(11) Deflorian, F.; Perez-Benito, L.; Lenselink, E. B.; Congreve, M.; van Vlijmen, H. W.; Mason, J. S.; Graaf, C. d.; Tresadern, G. Accurate prediction of GPCR ligand binding affinity with free energy perturbation. *Journal of Chemical Information and Modeling* **2020**, *60*, 5563–5579.
